# Mapping Brain-Behavior Space Relationships Along the Psychosis Spectrum

**DOI:** 10.1101/2020.09.15.267310

**Authors:** Jie Lisa Ji, Markus Helmer, Clara Fonteneau, Joshua B. Burt, Zailyn Tamayo, Jure Demšar, Brendan Adkinson, Aleksandar Savić, Katrin H. Preller, Flora Moujaes, Franz X. Vollenweider, William J. Martin, Grega Repovš, John D. Murray, Alan Anticevic

## Abstract

Difficulties in advancing effective patient-specific therapies for psychiatric disorders highlight a need to develop a stable neurobiologically-grounded mapping between neural and symptom variation. This gap is particularly acute for psychosis-spectrum disorders (PSD). Here, in a sample of 436 cross-diagnostic PSD patients, we derived and replicated a dimensionality-reduced symptom space across hallmark psychopathology symptoms and cognitive deficits. In turn, these symptom axes mapped onto distinct, reproducible brain maps. Critically, we found that multivariate brain-behavior mapping techniques (e.g. canonical correlation analysis) do not produce stable results. Instead, we show that a univariate brain-behavioral space (BBS) can resolve stable individualized prediction. Finally, we show a proof-of-principle framework for relating personalized BBS metrics with molecular targets via serotonin and glutamate receptor manipulations and gene expression maps. Collectively, these results highlight a stable and data-driven BBS mapping across PSD, which offers an actionable path that can be iteratively optimized for personalized clinical biomarker endpoints.

## Introduction

Mental health conditions cause profound disability with most treatments yielding limited efficacy across psychiatric symptoms (***Tohen et al., 2003***; ***McEvoy et al., 2007***; ***aan het Rot et al., 2010***; ***Cipriani et al., 2018***). A key step towards developing more effective therapies for specific psychiatric symptoms is reliably mapping them onto underlying neural systems. This goal is particularly challenging because neuropsychiatric diagnostics still operate under “legacy” categorical constraints, which were not quantitatively informed by neural or symptom data.

Critically, diagnostic systems in psychiatry [e.g. the Diagnostic and Statistical Manual of Mental Disorders (DSM) (***Association and on DSM-IV., 1994***)] were built to aid clinical consensus, but were not designed to guide quantitative mapping of symptoms onto neural alterations (***Phillips et al., 2008***; ***Gillihan and Parens, 2011***). Consequently, this system cannot optimally map onto patient-specific brain-behavioral alterations. This challenge is particularly evident along the psychosis spectrum disorders (PSD) where there is notable shared symptom variation across what distinct DSM diagnostic categories, including schizophrenia (SZP), schizo-affective (SADP), bipolar disorder with psychosis (BPP). For instance, despite BPP being a distinct diagnosis, BPP patients exhibit similar but attenuated psychosis symptoms and neural alterations similar to SZP (e.g. thalamic functional connectivity (FC) (***Anticevic et al., 2013b***)). It is essential to quantitatively map such shared clinical variation onto common neural alterations, to circumvent constraints for biomarker development (***Casey et al., 2013***; ***Phillips et al., 2008***; ***Gillihan and Parens, 2011***) - a key goal for development of neurobiologically-informed personalized therapies (***Anticevic et al., 2013b***; ***Casey et al., 2013***).

Recognizing the limitations of categorical frameworks, the NIMH’s Research Domain Criteria (RDoC) initiative introduced dimensional mapping of functional domains on to neural circuits (***Insel, 2014***). This motivated cross-diagnostic multi-site studies for mapping PSD symptom and neural variation (***Tamminga et al., 2014***; ***Koutsouleris et al., 2018***; ***Casey et al., 2018***; ***Di Martino et al., 2014***). Multivariate neuro-behavioral analyses across PSD and mood spectra reported brain-behavioral relationships across diagnoses, with the goal of informing individualized treatment (***Drysdale et al., 2017***). These studies attempted to address the challenge of moving beyond traditional *a priori* clinical scales, which provide composite scores (***Cronbach and Meehl, 1955***) that may not optimally capture neural variation (***Gillihan and Parens, 2011***; ***Barch et al., 2013***). For instance, despite many data-driven dimensionality-reduction symptom studies (***van der Gaag et al., 2006a***; ***Lindenmayer et al., 1994***; ***Emsley et al., 2003***; ***Dollfus et al., 1996***; ***Blanchard and Cohen, 2006***; ***Chen et al., 2019***; ***Lefort-Besnard et al., 2018***), a common approach in PSD neuroimaging research is still to sum “positive” or “negative” psychosis symptoms into a single score for relating to neural measures (***Ji et al., 2019b***; ***Anticevic et al., 2013b***). Importantly, neural alterations in PSD may reflect a more complex weighted combination of symptoms than *a priori* composite scores.

While multivariate neuro-behavioral studies have addressed this, such studies face the risk of failing to replicate due to overfitting (***Dinga et al., 2019***), arising from high dimensionality of behavioral and neural features and a comparatively limited number of subjects (***Helmer et al., 2020***). Notably, current state-of-the-art large-scale clinical neuroimaging studies have target enrollment totals of ~200-600 subjects (https://www.humanconnectome.org/disease-studies). We therefore wanted to test whether a stable, reproducible neuro-behavioral geometry can be derived with a sample size that is on par with current large-scale studies.

We hypothesized that a linearly-weighted low-dimensional symptom solution (capturing key disease-relevant information) produces a stable and robust univariate brain-behavioral mapping. Indeed, recent work used dimensionality reduction methods successfully to compute a neural mapping across canonical SZP symptoms (***Chen et al., 2019***). However, it remains unknown if this approach generalizes across PSD. Moreover, it is unknown if incorporating cognitive assessment, a hallmark and untreated PSD deficit (***Barch et al., 2013***), explains neural feature variation that is distinct from canonical PSD symptoms. Finally, prior work has not tested if a low-dimensional symptom-neural mapping can be predictive at the single patient level - a prerequisite for individualized clinical endpoints.

To inform these gaps, we first tested two key questions: i) Can data-reduction methods reliably reveal symptom axes across PSD that include both canonical symptoms and cognitive deficits? ii) Do these lower-dimensional symptom axes map onto a reproducible brain-behavioral solution across PSD? Specifically, we combined fMRI-derived resting-state measures with psychosis and cognitive symptoms (***Canuso et al., 2008***; ***Kay et al., 1987***) obtained from a public multi-site cohort of 436 PSD patients and 202 healthy individuals collected by the Bipolar-Schizophrenia Network for Intermediate Phenotypes (BSNIP-1) consortium across 6 sites in North America (***Tamminga et al., 2014***). The dataset included included 150 patients formally diagnosed with BPP, 119 patients diagnosed SADP, and 167 patients diagnosed with SZP (**Appendix 1 - Table 2**). This cohort enabled cross-site symptom-neural comparisons across multiple psychiatric diagnostic categories, which we then mapped onto specific neural circuits. First, we tested if dimensionality reduction of PSD symptoms revealed a stable solution for individual patient prediction. Next, we tested if this lowdimensional symptom solution yields novel, stable, and statistically robust neural mapping compared to canonical composite symptom scores or DSM diagnoses. We then tested if the computed symptom-neural mapping is reproducible across symptom axes and actionable for individual patient prediction. Finally, we anchor the derived symptom-relevant neural maps by computing their similarity against mechanistically-informed neural maps. We used independently collected pharmacological fMRI maps from healthy adults in response to putative PSD receptor treatment targets (glutamate via ketamine and serotonin via *LSD*) (***Preller et al., 2018***; ***Anticevic et al., 2015***). We also computed transcriptomic maps from the Allen Human Brain Atlas (AHBA) (***Hawrylycz et al., 2012***; ***Burt et al., 2018***) for genes implicated in PSD. The primary purpose of this paper is to derive a comprehensive, stable, and reproducible symptom-neural geometry across symptom domains in chronic PSD, that can be resolved at the single-subject level and mechanistically linked to molecular benchmarks. This necessitated the development of novel methods and rigorous testing (as we found that existing multivariate methods are drastically underpowered (***Helmer et al., 2020***)) which we have also presented in the paper for transparency and reproducibility. This approach can be iteratively improved upon to further elucidate the neural underpinnings of PSD variation, and expanded to apply to broader research questions and psychiatric spectra, symptom domains, and neural features.

To our knowledge, no study to date has mapped a data-reduced symptom geometry encompassing a transdiagnostic PSD cohort across combined cognitive and psychopathology symptom domains, with demonstrable statistical stability and reproducibility at the single subject level. Additionally, this symptom geometry can be mapped robustly onto neural data to achieve reproducible group-level effects as well as individual-level precision following feature selection. Furthermore, while other other studies have evaluated relationships between neural maps and complementary molecular datasets (e.g. transcriptomics), no study to our knowledge has benchmarked single-subject selected neural maps using reproducible neuro-behavioral features against both pharmacological neuroimaging maps and gene expression maps that may inform single subject selection for targeted treatment. Collectively, this study used data-driven dimensionality-reduction methods to derive stable orthogonal symptom dimensions across 436 PSD patients and mapped them onto novel and stable neuroimaging features. These effects were then benchmarked against molecular imaging targets, outlining an actionable path towards making personalized recommendations for clinical endpoints.

## Methods and Materials

### Overall Data Collection and Study Design

We used publicly available behavioral and neural data from the Bipolar-Schizophrenia Network on Intermediate Phenotypes (BSNIP) consortium (***Tamminga et al., 2014***). All data were obtained from the National Data Archive (NDA) repository (https://nda.nih.gov/edit_collection.html?id=2274). Participants were collected at 6 sites across the United States. Full recruitment details are provided in prior publications (***Tamminga et al., 2014***; ***Meda et al., 2015***; ***Sheffield et al., 2017***). In brief, participants were recruited through advertising in Baltimore MD, Boston MA, Chicago IL, Dallas TX, and Hartford CT. All assessments were standardized across sites. Participants were excluded if they had i) a history of seizures or head injury resulting in >10 minutes loss of consciousness, ii) positive drug screen on the day of testing, iii) a diagnosis of substance abuse in the past 30 days or substance dependence in the past 6 months, iv) history of serious medical or neurological disorder that would likely affect cognitive functioning, v), history of serious medical or neurological disorder that would likely affect cognitive functioning, vi) insufficient English proficiency, or vii) an age-corrected Wide-Range Achievement Test (4th edition) reading test standard score <65. Additionally, participants were required to have had no change in medication and been clinically stable over the past month. Participants completed a SCID interview and were given a diagnosis via consensus from study clinicians; participants with an Axis 1 clinical psychosis diagnosis were additionally given assessments including the Positive and Negative Symptom Scale (PANSS)(***Kay et al., 1987***). Note that apart from the measures used in this paper, the full BSNIP dataset includes a rich characterization of measures from multiple modalities, including electrophysiological, eye tracking, structural and diffusion neuroimaging, as well as a number of clinical batteries, which are not quantified in this study. We used data from a total of 638 participants with complete behavioral and neural data after preprocessing and quality control, including 202 healthy controls (CON), 167 patients diagnosed with schizophrenia (SZP), 119 patients with schizoaffective disorder (SADP), and 150 patients with bipolar disorder with psychosis (BPP). For full demographic and clinical characteristics of the sample see **Appendix 1 - Table 2**.

### Neural Data Acquisition and Preprocessing

Participants completed a neural magnetic resonance imaging (MRI) scan at 3T, including restingstate functional blood-oxygen-level-dependent imaging (BOLD) and a magnetization-prepared rapid gradient-echo (MP-RAGE) sequence for T1 weighted data. Full details on scanners and acquisition parameters used at each of the sites have previously been described (***Meda et al., 2012***) and are summarized here in **Appendix 1 - Table 3**. Neuroimaging data were preprocessed using the Human Connectome Project (HCP) minimal preprocessing pipeline (***Glasser et al., 2013***), adapted for compatibility with “legacy” data, which are now featured as a standard option in the HCP pipelines provided by our team (https://github.com/Washington-University/HCPpipelines/pull/156). These modifications to the HCP pipelines were necessary as the BSNIP data did not include a standard field map and did not incorporate a T2w high-resolution image. The adaptations for singleband BOLD acquisition have previously been described in detail (***Ji et al., 2019a***).

A summary of the HCP pipelines is as follows: the T1-weighted structural images were first aligned by warping them to the standard Montreal Neurological Institute-152 (MNI-152) brain template in a single step, through a combination of linear and non-linear transformations via the FM-RIB Software Library (FSL) linear image registration tool (FLIRT) and non-linear image registration tool (FNIRT) (***Jenkinson et al., 2002***). Next, FreeSurfer’s recon-all pipeline was used to segment brain-wide gray and white matter to produce individual cortical and subcortical anatomical segmentations (***Reuter et al., 2012***). Cortical surface models were generated for pial and white matter boundaries as well as segmentation masks for each subcortical grey matter voxel. Using the pial and white matter surface boundaries, a ‘cortical ribbon’ was defined along with corresponding subcortical voxels, which were combined to generate the neural file in the Connectivity Informatics Technology Initiative (CIFTI) volume/surface ‘grayordinate’ space for each individual subject (***Glasser et al., 2013***). BOLD data were motion-corrected by aligning to the middle frame of every run via FLIRT in the initial NIFTI volume space. In turn, a brain-mask was applied to exclude signal from non-brain tissue. Next, cortical BOLD data were converted to the CIFTI gray matter matrix by sampling from the anatomically-defined gray matter cortical ribbon and subsequently aligned to the HCP atlas using surface-based nonlinear deformation (***Glasser et al., 2013***). Subcortical voxels were aligned to the MNI-152 atlas using whole-brain non-linear registration and then the Freesurfer-defined subcortical segmentation applied to isolate the subcortical grayordinate portion of the CIFTI space.

After the HCP minimal preprocessing pipelines, movement scrubbing was performed (***Power et al., 2013***). As done in our prior work (***Anticevic et al., 2012c***), all BOLD image frames with possible movement-induced artifactual fluctuations in intensity were flagged using two criteria: frame displacement (the sum of the displacement across all six rigid body movement correction parameters) exceeding 0.5 mm (assuming 50 mm cortical sphere radius) and/or the normalized root mean square (RMS) (calculated as the RMS of differences in intensity between the current and preceding frame, computed across all voxels and divided by the mean intensity) exceeding 1.6 times the median across scans. Any frame that met one or both of these criteria, as well as the frame immediately preceding and immediately following, were discarded from further preprocessing and analyses. Subjects with more than 50% frames flagged using these criteria were excluded. Next, a high-pass filter (threshold 0.008 Hz) was applied to the BOLD data to remove low frequency signals due to scanner drift. In-house Matlab code was used to calculate the average variation in BOLD signal in the ventricles, deep white matter, and across the whole grey matter, as well as movement parameters. These signals, as well as their first derivatives to account for delayed effects, were then regressed out of the grey matter BOLD time series as nuisance variables (as any change in the BOLD signal due to these variables are persistent and likely not reflecting neural activity) (***Power et al., 2018***). It should be noted that using global signal regression to remove spatially persistent artifact is highly controversial in neuroimaging (***Power et al., 2017***; ***Yang et al., 2016***), but it remains the field-wide gold standard (though see other recent and emerging approaches at (***Glasser et al., 2018***; ***Aquino et al., 2019***)).

### Behavioral Symptom and Cognitive Data

The behavioral measures analyzed included the Positive and Negative Syndrome Scale (PANSS), an assessment of psychosis symptom severity (***Kay et al., 1987***), and the Brief Assessment of Cognition in Schizophrenia (BACS) battery, which provided an assessment of cognitive functioning (***Keefe et al., 2004***). Only control subjects with complete BACS measures were used for analyses; PANSS symptom scores were imputed for control subjects for whom the PANSS had not been administered under the assumption that these subjects were asymptomatic. Only patient subjects with complete PANSS and BACS measures were used in analyses. The BACS scores used here are presented as standardized Z-scores normalized to the mean and standard deviation of the control group for each BSNIP site, as done in prior work (***Tamminga et al., 2014***). The full PANSS battery is conventionally divided into three sub-scales: Positive symptom scale (7 items), Negative symptom scale (7 items), and General Psychopathology symptom scale (16 items). The BACS battery consists of 6 individual sub-scores (***Keefe et al., 2004***). In total, this yielded 36 symptom variables per participant. Effects of symptom variation across assessment sites have been rigorously characterized in prior publications (***Tamminga et al., 2013***). Nevertheless, we explicitly tested for site effects in our analyses, described in detail below.

### Principal Component Analysis of Behavioral Symptom Measures

The principal component analysis (PCA) of behavioral data was computed by including all 36 symptom variables across all N=436 patients. Variables were first scaled to have unit variance across patients before running the PCA. Significance of the derived principal components (PCs) was computed via permutation testing. For each permutation, patient order was randomly shuffled for each symptom variable before re-computing PCA. This permutation was repeated 5,000 times to establish the null model. PCs which accounted for a proportion of variance that exceeded chance (p<0.05 across all 5000 permutations) were retained for further analysis.

### Stability and Reproducibility of the Symptom PCA

Stability of the symptom PCA solution was assessed with within-sample cross-validation. To evaluate if there were site differences that uniquely drove the PCA solution, we performed a leave-site-out cross-validation analysis. Specifically, we re-ran the PCA using all patients except those from one site, which was held out. This process was repeated for each of the 6 sites. To further evaluate the stability of the derived PCA solutions, we performed 1,000 runs of *k*-fold cross-validation for values of *k* between 2 and 10. For each *k*-fold run, the full sample of patients was randomly divided into equally-sized *k* sets and a PCA was re-computed using subjects in *k*-1 sets (as the left-out set was used as the held-out sample).

For each run of leave-site-out and *k*-fold cross-validations significance was assessed via permutation testing as described above. The number of significant PCs and the total proportion of variance explained by these significant PCs remained highly stable across all runs (see **Fig. 2A** and **Appendix 1 - Fig. 5** and **Appendix 1 - Fig. 6**). Additionally, we compared observed and predicted PCA scores for the held-out sample of patients. Predicted scores for the held-out sample of patients were computed by weighing their raw symptom scores with the loadings from the PCA computed in all other patients. Observed scores for held-out patients were obtained from the original PCA computed on the full sample presented in the main text. The similarity between predicted and observed scores was high for all five significant PCs across all runs of leave-site-out and *k*-fold cross-validation, exceeding *r*=0.9 in most analyses (see **Fig. 2B-C** and **Appendix 1 - Fig. 5** and **6**). Notably, the PCA solution did not show medication status or dosage effects (**Appendix 1 - Fig. 7**). We also examined effects of age, sex, and SES in **Appendix 1 - Fig. 8**. Briefly, we observed a significant positive relationship between age and *PC3* scores, which may be because older patients (whom presumably have been ill for a longer time) exhibit more severe symptoms along the positive *PC3 - Psychosis Configuration* dimension. We also observed a significant negative relationship between Hollingshead index of SES and *PC1* and PC2 scores. Lower *PC1* and *PC2* scores indicate poorer general functioning and cognitive performance respectively, which is consistent with higher Hollingshead indices (i.e. lower-skilled jobs or unemployment and fewer years of education). We also found significant sex differences in *PC2 - Cognitive Functioning, PC4 - Affective Valence,* and *PC5 - Agitation/Excitement* scores.

**Figure 1.**
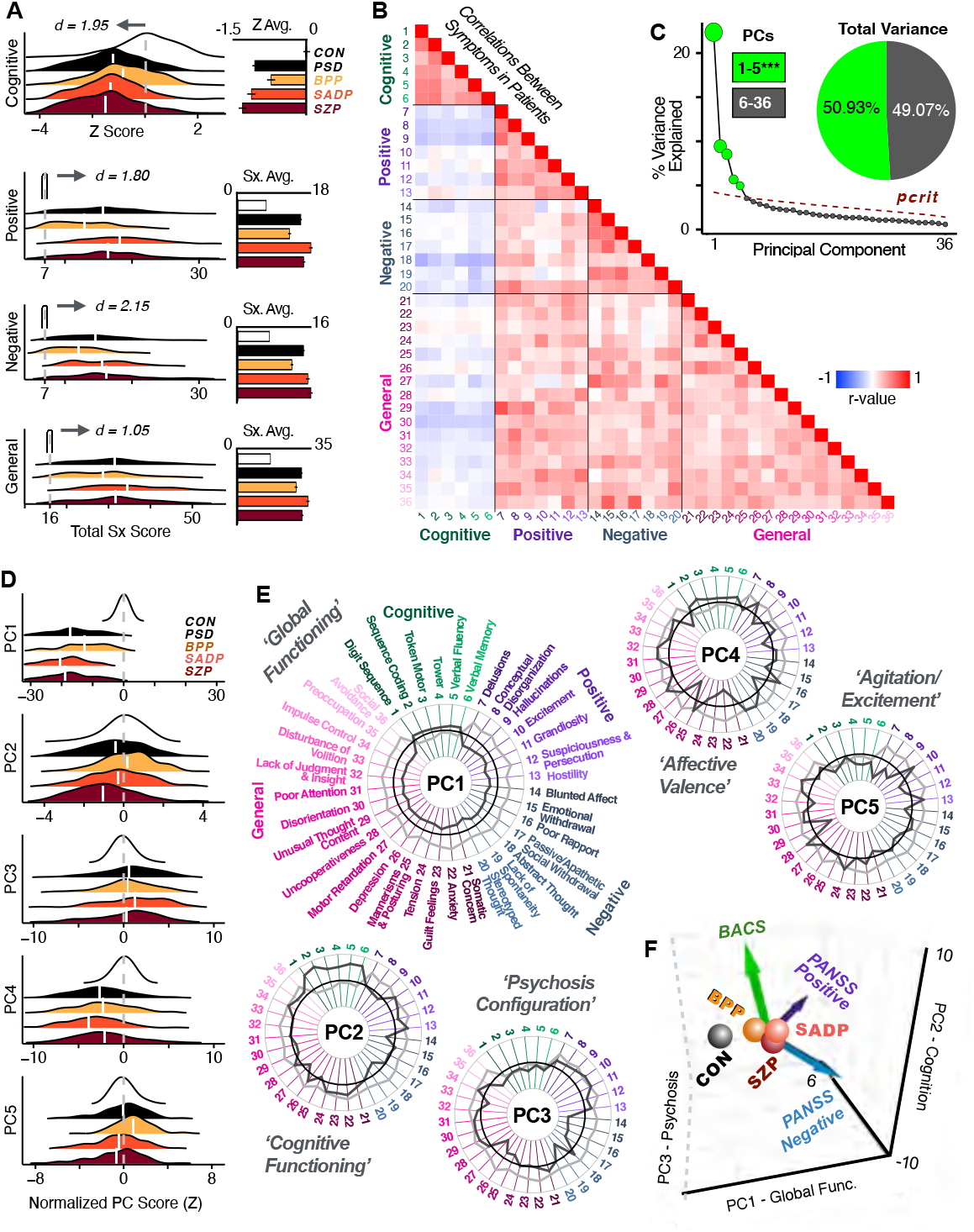
Quantifying data-driven low-dimensional variation of cross-diagnostic psychosis spectrum disorder symptoms and cognitive deficits. **A)** Distributions of symptom scores for each of the DSM diagnostic groups across core psychosis symptom meaures (PANSS positive, negative, and general symptoms tracking illness severity) and cognitive deficits (BACS composite cognitive performance). BPP: bipolar disorder with psychosis (yellow, N=150);SADP: schizo-affective disorder (orange, N=119); SZP - schizophrenia (red, N=167); All PSD patients (black, N=436);Controls (white, N=202). Bar plots show group means;error bars show standard deviations. **(B)** Correlations between 36 symptom measures across all PSD patients (N=436). **(C)** Screeplot shows the % variance explained by each of the principal components (PCs) from a PCA performed using all 36 symptom measures across 436 PSD patients. The size of each point is proportional to the variance explained. The first five PCs (green) survived permutation testing (p<0.05, 5,000 permutations). Together they capture 50.93% of all symptom variance (inset). **(D)** Distribution plots showing subject scores for the 5 significant PCs for each of the clinical groups, normalized relative to the control group. Note that control subjects (CON) were not used to derive the PCA solution;however, all subjects, including CON, can be projected into the data-reduced symptom geometry. **(E)** Loading profiles shown in dark grey for the 36 PANSS/BACS symptom measures on the 5 significant PCs. Each PC (“Global Functioning”, “Cognitive Functioning”, “Psychosis Configuration”, “Affective Valence”, “Agitation/Excitement”) was named based on the pattern of loadings on symptom measures. See **Appendix 1 - Fig. 3G** for numerical loading values. The PSD group mean score on each symptom measure is also shown, in light grey (scaled to fit on the same radarplots). Note that the group mean configuration resembles the *PC1* loading profile most closely (as *PC1* explains the most variance in the symptom measures). **(F)** PCA solution shown in the coordinate space defined by the first three PCs. Colored arrows show *a priori* composite PANSS/BACS vectors projected into the *PC1-3* coordinate space. The *a priori* composite symptom vectors do not directly align with data-driven PC axes, highlighting that PSD symptom variation is not captured fully by any one aggregate *a priori* symptom score. Spheres denote centroids (i.e. center of mass) for each of the patient diagnostic groups and control subjects. Alternative views showing individual patients and controls projected into the PCA solution are in **Appendix 1 - Fig. 3A-F**.

**Figure 2.**
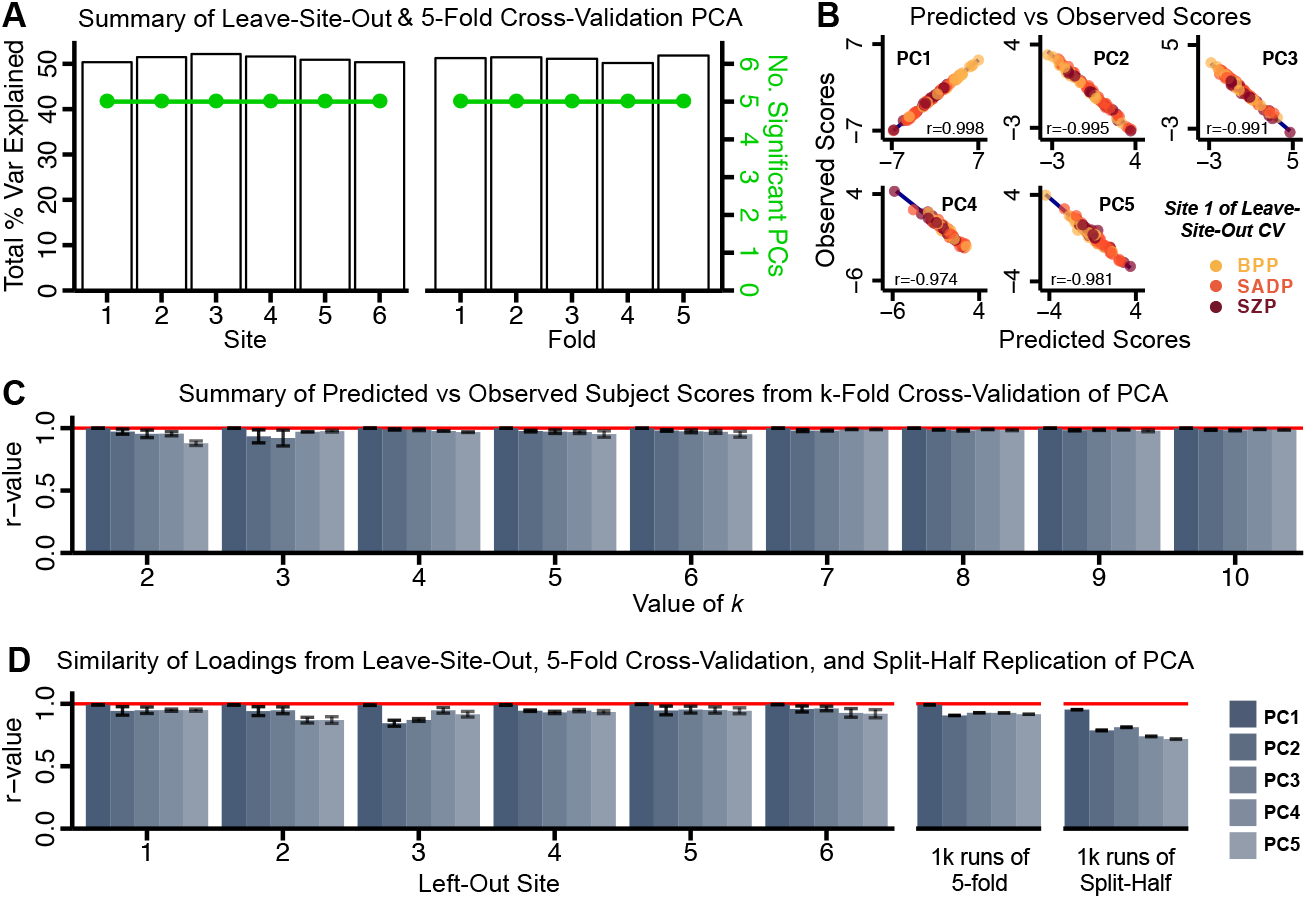
Dimensionality reduction of PSD symptom measures is highly stable and reproducible. **(A)** PCA solutions for leave-one-site out cross-validation (left) and 5-fold bootstrapping (right) explain a consistent total proportion of variance with a consistent number of significant PCs after permutation testing. Full results available in **Appendix 1 - Fig. 5** and **Appendix 1 - Fig. 6**. **(B)** Predicted versus observed single subject PC scores for all five PCs are shown for an example site (here Site 1). **(C)** Mean correlations between predicted and observed PC scores across all patients calculated via *k*-fold bootstrapping for *k*=2-10. For each *k,* patients were randomly split into *k* folds. For each fold, a subset of patients was held out and PCA was performed on the remaining patients. Predicted PC scores for the held-out patients were computed from the PCA obtained in all other retained patients. Original observed PC scores for the held-out patients were then correlated with the predicted PC scores. **(D)** Mean correlations between predicted and observed symptom measure loadings are shown for leave-one-site-out cross-validation (left), across 1,000 runs of 5-fold cross-validation (middle), and 1,000 split-half replications (right). For split-half replication, loadings were compared between the PCA performed in each independent half sample. Note: for panels **C-D** correlation values were averaged all *k* runs, all 6 leave-site-out-runs, or all 1,000 runs of the 5-fold cross-validation and split-half replications. Error bars indicate the standard error of the mean.

We also assessed the similarity of the PCA loadings using leave-site-out and 1,000 runs of 5-fold cross-validation frameworks (**Fig. 2D**). Importantly, this cross-validation was designed to test if the observed loadings remained stable (as opposed to predicted patient-level scores). The loadings for significant PCs from each leave-site-out PCA solution as well as each run of the 5-fold crossvalidation were highly correlated (**Fig. 2D**.

We also assessed the reproducibility of the PCA solution using independent split-half samples. The leave-site-out and *k*-fold cross-validation PCA analyses by definition use overlapping samples of patients for each iteration. Therefore, we additionally conducted a full split-half replication using entirely non-overlapping sets of patients in each iteration. For each split-half iteration, the full patient sample was randomly divided into two sets with equal proportions of each of the three diagnostic groups (BPP, SADP, SZP). Then, a PCA was computed using each of the split-half patient samples. The loadings from the two PCA solutions were then evaluated for reproducibility. This process was repeated 1,000 times. The loadings for significant PCs were again highly similar even when comparing PCA solutions derived from completely non-overlapping patient samples (**Fig. 2D**).

To predict individual patient PC scores for the leave-one-out analysis (**Appendix 1 - Fig. 27A**), a PCA was computed using all patients except one held-out patient (N=435). In turn, the derived loadings were then used to compute the predicted PC scores for the left-out patient. This process was repeated until predicted PC scores were calculated for every patient. Finally, the predicted score for each patient was evaluated for reproducibility relative to the observed score obtained from the PCA solution computed using the full N=436 sample of patients. In addition, we computed an independent component analysis (ICA) to evaluate the consistency of the behavioral data reduction solution across methods (see **Methods**).

### Global Brain Connectivity Calculation

Following preprocessing, the functional connectivity (FC) matrix was calculated for each participant by computing the Pearson’s correlation between every grayordinate in the brain with all other grayordinates. A Fisher’s r-to-Z transform was then applied. Global brain connectivity (GBC) was calculate by computing every grayordinate’s mean FC strength with all other grayordinates (i.e. the mean, per row, across all columns of the FC matrix). GBC is a data-driven summary measure of connectedness that is unbiased with regards to the location of a possible alteration in connectivity (***Cole et al., 2016***) and is therefore a principled way for reducing the number of neural features while assessing neural variation across the entire brain.

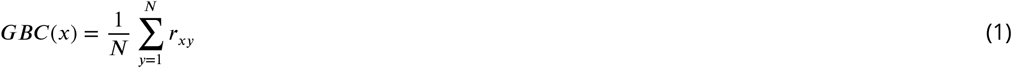

- where *GBC*(*x*) denotes the GBC value at grayordinate *x*;
- where *N* denotes the total number of grayordinates;
- where 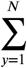 denotes the sum from *y* = 1 to *y* = *N*;
- where *r_xy_* denotes the correlation between the time-series of grayordinates *x* and *y*;

For parcel-wise GBC maps (described below) we first computed the mean BOLD signal within each parcel (see section below for parcellation details) for each participant and then computed the pairwise FC between all parcels. Finally, to obtain the parcellated GBC metric we computed the mean FC for each parcel and all other parcels. This order of operations (first parcellating the dense data series and then computing GBC) was chosen because it resulted in stronger statistical values due to increased within-parcel signal-to-noise of the BOLD data (**Fig. 4A**).

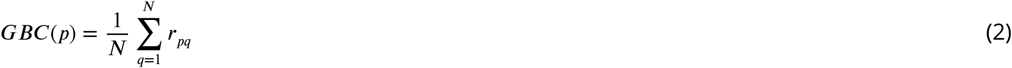

- where *GBC*(*p*) denotes the GBC value at parcel *p*;
- where *N* denotes the total number of parcels;
- where 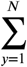 denotes the sum from *q* = 1 to *q* = *N*;
- where *r_pq_* denotes the correlation between the time-series of parcels *p* and *q*;

### Neural Data Reduction via Functional Brain-wide Parcellation

Here we applied a recently developed Cole-Anticevic Brain Network Parcellation (CAB-NP) parcellation (***Ji et al., 2019c***), which defines functional networks and regions across cortex and subcortex that leveraged the Human Connectome Project’s Multi-Modal Parcellation (MMP1.0) (***Glasser et al., 2016***; ***Ji et al., 2019c***). The final published CAB-NP 1.0 parcellation solution can be visualized via the Brain Analysis Library of Spatial maps and Atlases (BALSA) resource (https://balsa.wustl.edu/rrg5v) and downloaded from the public repository (https://github.com/ColeLab/ColeAnticevicNetPartition). The cortex component of the parcellation solution is comprised of 180 bilateral parcels (a total of 360 across both left and right hemispheres), consistent with the Human Connectome Project’s Multi-Modal Parcellation (MMP1.0) (***Glasser et al., 2016***). The subcortex component is comprised of 358 parcels defined using resting-state functional BOLD covariation with the cortical network solution (***Ji et al., 2019c***).

### Mass Univariate Symptom-Neural Mapping

Behavioral scores (*a priori* and PCA) were quantified in relation to individual GBC variation (either dense or parcellated) via a mass univariate regression procedure. The resulting map of regression coefficients reflected the strength of the relationship between patients’ behavioral PC score and GBC at every neural location, across all 436 patients. The greater the magnitude of the coefficient for a given location, the stronger the statistical relationship between GBC and the behavioral variation across patients. The coefficients were then Z-scored for each map. Significance of the maps was assessed via nonparametric permutation testing, 2000 random shuffles with TFCE (***Smith and Nichols, 2009***) type-I error-protection computed via the Permutation Analysis of Linear Models program (***Winkler et al., 2014***).

### Cross-validation and Split-half Replication of Mass Univariate Symptom-Neural Mapping

Stability of the symptom-neural mapping was assessed following the same cross-validation logic as described for the symptom-driven PCA solutions. Specifically, five-fold cross-validation was performed by first randomly partitioning all patients (N=436) into 5 subsets. Regression of the behavioral PC scores onto GBC across patients was performed while holding out 1/5 of the patient sample (N=349). The correlation between the resulting neural coefficient map was then computed with the neural map obtained from the full sample calculation. For leave-site-out cross-validation, all subjects except those from one site were used when calculating the symptom-neural regression model. The resulting maps were compared to the map from the full sample regression. This process was repeated 6 times, each time leaving out subjects in one of the 6 sites.

Reproducibility of the symptom-neural mapping was assessed with 1,000 iterations of split-half replication. For each iteration, the full sample of subjects was first randomly split into two halves (referred to as H1 and H2) with the proportion of subjects in each diagnostic group (BPP, SADP, SZP) preserved within each half. For each iteration we used the H1 PCA solution and loadings to compute the predicted PCA scores for H2. In turn, the observed H1 scores were computed from a PCA loadings on the same H1 half-sample of patients. These H1 scores were then regressed against parcellated GBC for patients in H2. This coefficient GBC map reflects the strength of the relationship between the predicted PC score and GBC across H2 patients. Finally, the GBC coefficient maps derived from the H1 observed and H2 predicted PCA scores were then correlated for each PC axis. This process was then repeated 1,000 times and evaluated for reproducibility.

### Principal Component Analysis of Neural Data

To evaluate the consistency of the neural GBC, we computed a PCA solution for the parcellated neural GBC data for all 202 control subjects as well as for all 436 patients (**Appendix 1 - Fig. 20**). The resulting neural GBC-derived PCs capture the striking consistency of the neural variance components irrespective of clinical status, which highlights that the bulk of the neural variance is not symptom-relevant. As with the behavioral PCA, significance of the neural PCA solution was assessed via permutation testing (1,000 random shuffles of parcels within subject). To evaluate the effect of shared neural variance on the symptom-neural solution, we parsed out common neural PCs from the PSD GBC data. We first conducted a PCA using the parcellated GBC data from all 436 PSD and 202 CON (a matrix with dimensions 638 subjects x 718 parcels). This GBC-PCA resulted in 637 independent GBC-PCs. Since PCs are orthogonal to each other, we then partialled out the variance attributable to GBC-PC1 from the PSD data by reconstructing the PSD GBC matrix using only scores and coefficients from the remaining 636 GBC-PCs 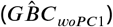. We then reran the univariate regression as described in **Appendix 1 - Fig. 3**, using the same five symptom PC scores across 436 PSD. We also repeated the symptom-neural regression with the first 2 GBC-PCs partialled out of the PSD data 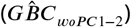, with the first 3 PCs parsed out 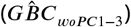, and with the first 4 neural PCs parsed out 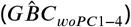.

### Canonical Correlation Analysis

Canonical correlation analysis (CCA) is a multivariate statistical technique which examines simultaneously the relationships between multiple independent variables and multiple dependent variables by computing linear combinations of each variable set that maximizes the correlations between the two sets (**Fig. 5A**). Here, the two variates used were symptom and neural features across all subjects. Each feature was Z-scored prior to computing the CCA solution. Given the size of the ‘dense’ neural feature space reduced the number of neural features in a principled manner via the described parcellation. This reduced the number of neural features to 180 symmetrized cortical parcels (i.e. GBC was averaged for each pair of analogous parcels in the left and right hemispheres). Critically, corresponding cortical parcels in the left and right hemispheres of this parcellation have been shown to be highly similar (***Glasser et al., 2013***; ***Burt et al., 2018***). First, we computed the CCA solution using all 36 item-level symptom measures as behavioral features. In turn, we computed an additional CCA solution using scores from the significant 5 PCs.

Each symptom PC is a weighted linear composite of the original symptom measures and each behavioral CV is a weighted linear composite of the symptom PCs. Therefore, to compute the loadings of the 36 symptom measures on each of the behavioral CVs (computed using the symptom PCs), we multiplied the matrix of loadings from the CCA with the matrix of loadings from the PCA (**Fig. 5J**).

### CCA Cross-validationand Split-half Replication

Stability of the CCA solution was assessed via cross-validation. Five-fold cross-validation of the CCA was performed by first randomly partitioning all patients (N=436) into 5 subsets. The CCA was then performed between neural and behavioral features using all but one of the subsets. The results of this CCA was then compared to the full sample CCA. **Appendix 1 - Fig. 18A-D** shows the comparisons across all 5 five-fold cross-validation runs of the CCA compared to the full model, including neural factor loadings, behavioral factor loadings, and projected symptom measure loadings. For leave-site-out cross-validation, all subjects except those from one site were used in the CCA. The resulting outputs were then compared to those from the full sample CCA. This process was repeated for all 6 sites. These data are shown in **Appendix 1 - Fig. 18E-H**.

For leave-one-out cross-validation of the CCA, one subject was held out as a CCA was performed using neural and behavioral features from the other 435 subjects. The loadings matrices *Ψ* and *Θ* from the CCA were then used to calculate the “predicted” neural and behavioral latent scores for all 5 canonical modes for the holdout subject. This was repeated for every subject, such that predicted neural and behavioral latent score matrices (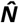 and 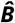) were computed, of the same dimensions as (***N*** and ***B***) respectively. The corresponding CVs in 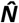 and 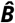 were then correlated across subjects, as shown in **Fig. 5M**. If the CCA solution is stable, these correlations should be comparable to the canonical correlations of the full CCA (**Fig. 5D**).

Reproducibility of the CCA solution was assessed via split-half replication. This was performed by first randomly splitting the full sample into two halves (H1 and H2) with the proportion of subjects in each diagnostic group (BPP, SADP, SZP) preserved within each half. A CCA was then performed separately in each half and the resulting outputs were then compared to each other. This process was repeated 1,000 times to obtain mean and standard deviation performance metrics. These data are shown in **Fig. 5L**.

### Multivariate CCA Power Analysis

Multivariate power analyses to estimate the minimum sample size needed to sufficiently power a CCA were computed using methods described in (***Helmer et al., 2020***), using the Generative Modeling of Multivariate Relationships tool (*gemmr,* https://github.com/murraylab/gemmr (v0.1.2)). Briefly, a model was built by: 1) Generating synthetic datasets for the two input data matrices, by sampling from a multivariate normal distribution with a joint covariance matrix that was structured to encode CCA solutions with specified properties;2) Performing CCAs on these synthetic datasets. Because the joint covariance matrix is known, the true values of estimated association strength, weights, scores, and loadings of the CCA, as well as the errors for these four metrics, can also be computed. In addition, statistical power that the estimated association strength is different from 0 is determined through permutation testing;3) Varying parameters of the generative model (number of features, assumed true between-set correlation, within-set variance structure for both datasets) the required sample size *n*_req_ is determined in each case such that statistical power reaches 90% and all of the above described error metrics fall to a target level of 10%;and 4) Fitting and validating a linear model to predict the required sample size *n*_req_ from parameters of the generative model. This linear model was then used to calculate *n*_req_ for CCA in three data scenarios: i) 718 neural vs. 5 symptom features;ii) 180 neural vs. 5 symptom features; iii) 12 neural vs. 5 symptom features.

### Independent Replication Dataset

To illustrate the generalizability of the single-subject prediction results across independent datasets and DSM disorders, we use an independently collected dataset consisting of 30 patients with a formal diagnosis of schizophrenia (SZP) and 39 patients diagnosed with obsessive compulsive disorder (OCD). These patients were recruited via clinician referral and regional advertising and assessed at the Connecticut Mental Health Center in New Haven, CT. Demographic and symptom data are shown in **Appendix 1 - Table 4**.

The behavioral assessment for the replication dataset included the PANSS and the Penn Computerized Neurocognitive Battery (PennCNB)(***Moore et al., 2015***). Items from the PennCNB were matched to items from the BACS as follows: BACS digital sequencing (working memory) -> LNB (N-back Total score); BACS symbol coding (attention) -> Penn Continuous Performance Test); BACS token motor task (motor) -> CTAP; BACS Tower of London (abstraction and mental flexibility) -> PCET or RAVEN (Penn Conditional Exclusion Test); BACS verbal fluency (verbal ability) -> SPVRT (Penn Verbal Reasoning Test); BACS verbal memory test -> PLLT (Penn List Learning Test) (***Gur et al., 2010***).

Neural data were collected using a Siemens 3T scanner with a 64 channel head coil at the Yale Center for Biomedical Imaging. Imaging acquisition parameters were aligned with those of the Human Connectome Project (HCP) (***Van Essen et al., 2013***). High-resolution T1w and T2w structural images were acquired in 224 AC-PC aligned slices, 0.8mm isotropic voxels. T1w images were collected with a magnetization-prepared rapid gradient-echo (MP-RAGE) pulse sequence (TR=2400 ms, TE=2.07 ms, flip angle=8o, field of view=256 x 256 mm). T2w images were collected with a SCP pulse sequence [TR=3200 ms, TE=564 ms, flip angle mode=T2 var, field of view=256 x 256 mm]. Resting-state BOLD images were collected with a multi-band accelerated fast gradient-echo, echo-planar sequence (acceleration factor=6, time repetition (TR)=800 ms, time echo (TE)=31.0 ms, flip angle=55o, field of view=210 x 210 mm, matrix=84 x 84, bandwidth=2290 Hz); 54 interleaved axial slices aligned to the anterior-posterior commissure (AC-PC) with 2.5mm isotropic voxels. Additionally, a pair of reverse phase-encoded spin-echo field maps (anterior-to-posterior and posterior-to-anterior) were acquired (voxel size=2.5 mm isotropic, TR=7220 ms, TE=73 ms, flip angle=90o, field of view=210 x 210 mm, bandwidth=2290 Hz). These neural data were preprocessed using the HCP minimal preprocessing pipeline as described in the section “Neural data acquisition and preprocessing” above, with the same motion scrubbing and nuisance signal regression parameters as were used for the BSNIP dataset.

### Neural Parcel *Δ*GBC Feature Selection and Prediction

For the purposes of patient selection, we were focused on individual differences in neural features as they co-vary in relation to individual symptom scores. However, this individual variation in neural features may be small compared to the overall group mean. Hence for each patient, we compute the difference in each brain location (parcel) between the patient’s actual GBC and the group mean GBC for that location. This is denoted by *Δ*GBC. Importantly, using a de-meaned GBC metric “standardizes” the data and helps to correct for possible differences in scanners/protocols across different datasets. Next, we developed an optimized univariate regression framework leveraging a “dot product” metric to relate a vector of neural features with a single symptom scalar value. This process for neural feature selection (results in **Appendix 1 - Fig. 27**) is shown as a systems flow diagram in **Appendix 1 - Fig. 25**. The observed dot product GBC metric (*dpGBC^obs^*) is computed as follows:

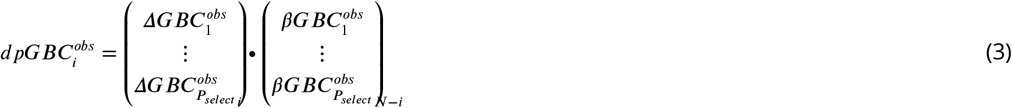

- where for a given subject 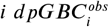 denotes the dot product GBC value of the two vectors 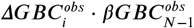 across all *P_select_* parcels,
- 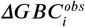 is a vector of length *P_select_* denoting a difference map of subject *i*’s *GBC^obs^* map relative to the group mean *GBC^obs^* map, within a given number of parcels *P_select_,*
- the 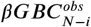 vector denotes the PCA-to-GBC statistical group-level *β* map for a lowdimensional PC symptom score across selected parcels *P_select_* for *N* subjects excluding subject *i*.

This calculation is then repeated for each subject *i*, resulting in a final vector 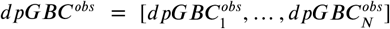 for *N* subjects. There are several key properties of the *dpGBC^obs^* statistic: i) it is not inflated by individual GBC map similarity to the group map because each subject’s *ΔGBC^obs^* map is demeaned relative to the reference group computed independently of the left-out subject; ii) this statistic is not biased by the parcel number (which drops with iterative selection) because the resulting *dpGBC^obs^* value variation is quantified relative to the low-dimensional symptom score across all subjects (see Eq. 2). Put differently, the final evaluation considers the relationship between *dpGBC^obs^* and the low-dimensional PC symptom scores across individuals; iii) The dot product statistic can yield both positive and negative values - a property which some map similarity measures lack (e.g. *η*^2^); iv) It is unbounded (unlike a correlation), which is key to maximize co-variation with low-dimensional symptom scores across individuals (see Eq. 2); v)The 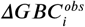 map for a given individual is projected onto the basis set of the 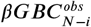 map, which is independent of the left-out individual but directly related to the low-dimensional PC symptom score variance, thus maximizing the dot product optimization.

Next, we select *N* – *i* individuals and compute a univariate regression where participants’ low dimensional symptom scores are regressed onto the 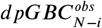 values:

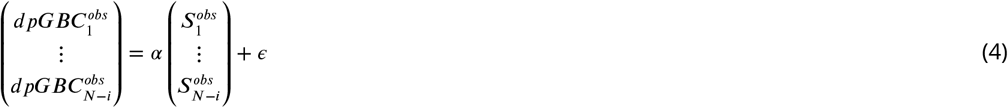

- where for *dpGBC^obs^* each element denotes the dot product value of the 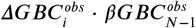 vectors per subject across *P* parcels,
- *α* denotes the regression coefficient in the univariate linear model,
- where for *S^obs^* each element denotes the observed low-dimensional symptom score (e.g. *PC3* score) per subject,
- *ϵ* denotes the error term in the univariate linear model.

After the *dpGBC^obs^* = *αS^obs^* + *ϵ* regression is computed on *N* – *i* subjects it is applied to the left-out-subject *i*. This is repeated for all *N* subjects and the model is evaluated for the number of parcels that maximize two key dot product evaluation metrics (**Fig. 6B**):

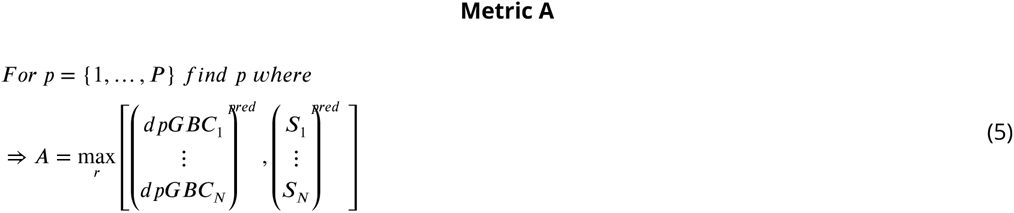

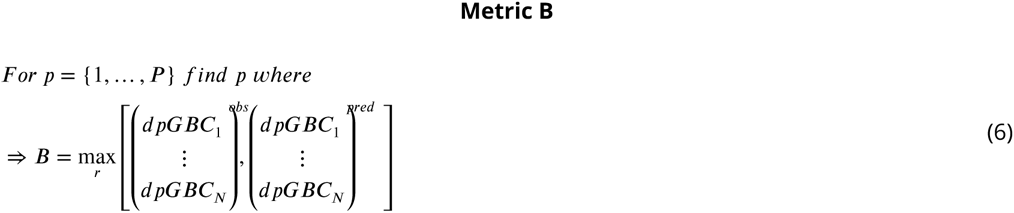

- where *A* denotes the maximum *r* correlation value for the two vectors of *dpGBC^pred^* values and the predicted *S^pred^* low-dimensional symptom scores (e.g. *PC3* axis) for *N* subjects from the leave-one-out cross-validation,
- where *B* denotes the maximum *r* correlation value for the two vectors of *obs* and *pred dpGBC* values for *N* subjects.

In the initial step in the step-down model all *P =* 718 parcels are retained in the initial dot product calculation. For each iteration of P selected parcels, the least predictable parcel *P* (i.e. the parcel with the weakest value in the *PC3* map) is eliminated from the map. Then, the step-down regression is repeated until *P* = 1.

### Pharmacological Neuroimaging Acquisition in Healthy Volunteers - LSD

Methods for the lysergic acid diethylamide (LSD) neuroimaging study are described in detail in prior publications (***Preller et al., 2018***). The use of LSD in humans was authorized by the Swiss Federal Office of Public Health, Bern, Switzerland. The study protocol was approved by the Can-tonal Ethics Committee of Zurich (KEK-ZH_No: 2014_0496). Participants received both written and oral descriptions of the study procedures and the effects and possible risks of LSD. All participants provided written informed consent in accordance with the Declaration of Helsinki. The study employed a fully double-blind, randomized, within-subject cross-over design with 3 conditions: (1) placebo + placebo (Pla) condition: placebo (179 mg Mannitol and Aerosil 1 mg po) after pretreatment with placebo (179 mg Mannitol and Aerosil 1 mg po); (2) Pla+LSD (LSD) condition: LSD (100 *μ*g po) after pretreatment with placebo (179 mg Mannitol and Aerosil 1 mg po), or (3) Ketanserin+LSD (Ket+LSD) condition: LSD (100 *μ*g po) after pretreatment with the 5-HT2A antagonist Ket (40 mg po). Data were collected for all subjects in a randomized counterbalanced order at three different sessions each two weeks apart. For all conditions, the first substance was administered 60 minutes before the second substance, and the first neural scan was conducted 75 minutes after the second administration, with a second scan conducted at 300 minutes post-administration. In the present study, only data from the two neural scans for the LSD and Pla conditions were evaluated.

Briefly, neuroimaging data acquisition details for the LSD study are as follows. MRI data were acquired on a Philips Achieva 3.0T whole-body scanner (Best, The Netherlands). A32-channel receiver head coil and MultiTransmit parallel radio frequency transmission was used. Images were acquired using a whole-brain gradient-echo planar imaging (EPI) sequence (repetition time=2,500 ms; echo time=27 ms; slice thickness=3 mm; 45 axial slices; no slice gap; field of view=240 mm^2^; in-plane resolution=3 mm × 3 mm; sensitivity-encoding reduction factor=2.0). 240 volumes were acquired per resting state scan resulting in a scan duration of 10 mins. Additionally, two high-resolution anatomical images were acquired using T1-weighted and T2-weighted sequences. T1-weighted images were collected via a 3D magnetization-prepared rapid gradient-echo sequence (MP-RAGE) with the following parameters: voxel size=0.7 mm^3^, time between two inversion pulses=3123 ms, inversion time=1055 ms, inter-echo delay=12 ms, flip angle=8°, matrix=320×335, field of view=224×235 mm, 236 sagittal slices. Furthermore T2-weighted images were collected using via a turbo spinecho sequence with the following parameters: voxel size=0.7 mm^3^, repetition time=2500 ms, echo time=415 ms, flip angle=90°, matrix=320 × 335, field of view=224 mm × 235 mm, 236 sagittal slices.

### Pharmacological Neuroimaging Acquisition in Healthy Volunteers - Ketamine

Similar to the LSD study, the ketamine pharmacological neuroimaging protocol employed a within-subject design where all healthy volunteer participants underwent a single scanning session consisting of two infusions: i) placebo (saline solution) followed by ii) ketamine infusion. The study was approved by the Yale University institutional review board. Participants received both written and oral descriptions of the study procedures and the effects and possible risks of ketamine, and all participants provided written informed consent before beginning the study. Healthy volunteers were informed prior to scanning that they would undergo one placebo run and one ketamine run but were blinded to the order of administration. Because of the sustained effects of ketamine, this infusion was always the second of the two runs, consistent with prior work (***Anticevic et al., 2012c***). A doctor and two nurses as well as a study coordinator remained present for the duration of the scan. A bolus of ketamine (0.3 mg/kg of bodyweight) or saline were delivered via infusion 5 sec after the start of the run and then continuously at a rate of 0.65 mg/kg per hour through the duration of the session. The sequence of scans in each for either the placebo or ketamine infusion was as follows: i) resting state (4.67 min); ii) blood draw (sham if saline condition); iii) a cognitive working memory task (total 14 min); iv) blood draw (sham if saline condition); v) a cognitive working memory task (total 14 min); vii) blood draw (sham if saline condition); vii) a cognitive working memory task(total 8.63 min). Data from the the cognitive working memory task were not used in the present study and are actively undergoing a distinct analysis. Participants were scanned at Yale University on a Siemens Trio 3T whole-body scanner and 32-channel receiver head coil. High-resolution structural T1-weighted images were acquired using an MP-RAGE sequence and the following parameters: voxel size=0.8 mm^3^, time between two inversion pulses=3123 ms, inversion time=1055 ms, interecho delay=12 ms, flip angle=8°, matrix=320×335, field of view=227×272 mm, 227 sagittal slices. T2-weighted images were acquired with the following parameters:voxel size=0.8 mm^3^, repetition time=2500 ms, echo time=415 ms, flip angle=90°, matrix=320 × 335, field of view=227×272 mm, 227 sagittal slices. BOLD images were acquired using a whole-brain gradient-echo planar imaging (EPI) sequence (400 frames at TR=0.7 ms; TE=30 ms; slice thickness=2.5 mm; 54 axial slices; no slice gap; field of view=250 mm^2^; in-plane resolution=2.5 mm × 2.5 mm). In addition, a field map and pair of spin-echo gradients were collected at the end of every scanning session.

### Pharmacological Neuroimaging Processing and Analysis for LSD and Ketamine Samples

All data were preprocessed to be consistent with the BSNIP and replication dataset processing steps. Specifically, we used the HCP minimal preprocessing pipeline and with the same motion scrubbing and nuisance signal regression parameters as were used for the BSNIP dataset as described in the section “Neural data acquisition and preprocessing” above. Parcel-wise GBC maps were computed as described in the section “Resting-state functional connectivity and global brain connectivity” above, by first parcellating the dense time-series for each subject and then computing the parcel-level GBC. As with the BSNIP and replication analyses, parcel GBC was first calculated by parcellating the dense time-series for every subject and then computing GBC across all 718 parcels from the whole-brain functional parcellation (***Ji et al., 2019c***; ***Glasser et al., 2016***). Group-level GBC maps were computed for every participant for all conditions. We computed the contrast map for “LSD-Placebo” as well as “Ketamine-Placebo” conditions as a t-test between both the pharmacological scans versus placebo scans. The Z-scored t-contrast maps between pharmacological and placebo conditions were used as a pharmacological target map in relation to the PC-derived neural GBC maps (**Fig. 7** and **Appendix 1 - Fig. 30**).

### Neural Gene Expression Mapping

The gene mapping analyses in this study utilize the procedure described in (***Burt et al., 2018***). Briefly, we used cortical gene expression data from the publicly available Allen Human Brain Atlas (AHBA, RRID:SCR_007416), mapped to cortex (***Burt et al., 2018***). Specifically, the AHBA quantified expression levels across 20,737 genes obtained from six postmortem human brains using DNA microarray probes sampled from hundreds of neuroanatomical loci. Recent studies demonstrated the ability to map expression of each gene onto neuroimaging-compatible templates (***Hawrylycz et al., 2012***; ***Burt et al., 2018***). Building on these innovations we mapped gene expression on to 180 symmetrized cortical parcels from the HCP atlas (***Glasser et al., 2016***) in line with recently published methods (***Burt et al., 2018***). This yielded a group-level map for each gene where the value in each parcel reflected the average expression level of that gene in the AHBA dataset. These group-level maps were in turn as a gene expression target map in relation to the PC-derived neural GBC maps.

### Data Availability

All primary results derive from data that is publicly available from sources described above. All symptom and neuroimaging data for the BSNIP-1 consortium was obtained via the NDA (https://nda.nih.gov/edit_collection.html?id=2274).

### Code Availability

Code and core results presented in this paper are available at https://bitbucket.org/jielisa_ji/bbs_manuscript/src/master/.

## Results

The key questions and results of this study are summarized in **Appendix 1 - Fig. 1** and an overview of the workflow is presented in **Appendix 1 - Fig. 2**. Additionally, for convenient reference, a glossary of the key terms and abbreviations used throughout the study are provided in **Appendix 1 - Table 1**.

### Dimensionality-Reduced PSD Symptom Variation is Stable and reproducible

First, to evaluate PSD symptom variation we examined core PSD psychopathology metrics captured by two instruments: the Brief Assessment of Cognition in Schizophrenia (BACS) and the Positive and Negative Syndrome Scale (PANSS). We refer to these 36 items as “symptom measures” throughout the rest of the paper. We observed group mean differences across DSM diagnoses (**Fig. 1A**, p<0.05 Bonferroni corrected); however, symptom measure distributions revealed notable overlap that crossed diagnostic boundaries (***Tamminga et al., 2014***; ***Keshavan et al., 2011***). Furthermore, we observed marked collinearity between symptom measures across the PSD sample (**Fig. 1B**), indicating that a dimensionality-reduced solution may sufficiently capture meaningful varation in this symptom space. We hypothesized that such a dimensionality-reduced symptom solution may improve PSD symptom-neural mapping as compared to pre-existing symptom scales.

Here we report results from a principal component analysis (PCA) as it produces a deterministic solution with orthogonal axes (i.e. no *a priori* number of factors needs to be specified) and explains all variance in symptom measures. Results were highly consistent with prior symptom-reduction studies in PSD: we identified 5 PCs (**Fig. 1C**), which captured ~50.93% of all variance (see **Methods** & **Appendix 1 - Fig. 3**) (***Chen et al., 2019***).

Critically, the key innovation here is the combined analysis across PSD diagnoses of canonical PSD symptoms and cognitive deficits, a fundamental PSD feature (***Barch et al., 2013***). The 5 PCs revealed few distinct boundaries between DSM categories (**Fig. 1D**). Furthermore, we observed substantial overlap in PC scores between controls and patients in all DSM diagnostic groups, with PC1 showing the greatest differentiation between PSD and the “normal-spectrum”. This overlap is not unexpected, given that PSD symptoms are present to some extent in “healthy” individuals not meeting DSM diagnostic criteria (***Verdoux and van Os, 2002***; ***Nuevo et al., 2012***; ***Kelleher and Cannon, 2011***; ***Stefanis et al., 2002***).

**Fig. 1E** highlights loading configurations of symptom measures forming each PC. To aid interpretation, we assigned a name for each PC based on its most strongly weighted symptom measures. This naming is qualitative but informed by the pattern of loadings of the original 36 symptom measures. For example, *PC1* was highly consistent with a general impairment dimension (i.e. “Global Functioning”); *PC2* reflected more exclusively variation in cognition (i.e. “Cognitive Functioning”); *PC3* indexed a complex configuration of psychosis-spectrum relevant items (i.e. “Psychosis Configuration”); *PC4* generally captured variation mood and anxiety related items (i.e. “Affective Valence”); finally, *PC5* reflected variation in arousal and level of excitement (i.e. “Agitation/Excitation”). For instance, a generally impaired patient would have a highly negative *PC1* score, which would reflect low performance on cognition and elevated scores on most other symptomatic items. Conversely, an individual with a high positive *PC3* score would exhibit delusional, grandiose, and/or hallucinatory behavior, whereas a person with a negative *PC3* score would exhibit motor retardation, social avoidance, possibly a withdrawn affective state with blunted affect (***Gelenberg, 1976***). Comprehensive loadings for all 5 PCs are shown in **Appendix 1 - Fig. 3G**. **Fig. 1F** highlights the mean of each of the 3 diagnostic groups (colored spheres)and healthy controls (black sphere) projected into a 3-dimensional orthogonal coordinate system for *PCs 1,2 & 3* (x,y,z axes respectively; alternative views of the 3-dimensional coordinate system with all patients projected are shown in **Appendix 1 - Fig. 3**). Critically, PC axes were not parallel with traditional aggregate symptom scales. For instance, *PC3* is angled at ~45° to the dominant direction of PANSS Positive and Negative symptom variation (purple and blue arrows respectively in **Fig. 1F**).

Because *PC3* loads most strongly on to hallmark symptoms of PSD (including strong positive loadings across PANSS Positive symptom measures in the PANSS and strong negative loadings onto most Negative measures), we focus on this PC as an opportunity to quantify an innovative, fully data-driven dimension of symptom variation that is highly characteristic of the PSD patient population. Additionally, this bi-directional symptom axis captured shared variance from measures in other traditional symptoms factors, such the PANSS General factor and cognition. We found that the *PC3* result provided a powerful empirical demonstration of how using a data-driven dimensionality-reduced solution (via PCA) can reveal novel patterns intrinsic to the structure of PSD psychopathology.

Notably, independent component analysis (ICA), an alternative dimensionality reduction procedure which does not enforce component orthogonality, produced similar effects for this PSD sample, see **Appendix 1 - Note 1** & **Appendix 1 - Fig. 4A**). Certain pairs of components between the PCA and ICA solutions appear to be highly similar and exclusively mapped (IC5 and PC4; IC4 and PC5) (**Appendix 1 - Fig. 4B**). On the other hand, PCs 1-3 and ICs 1-3 do not exhibit a one-to-one mapping. For example, *PC3* appears to correlate positively with *IC2* and equally strongly negatively with *IC3,* suggesting that these two ICs are oblique to the PC and perhaps reflect symptom variation that is explained by a single PC. The orthogonality of the PCA solution forces the resulting components to capture maximally separated, unique symptom variance, which in turn map robustly on to unique neural circuits. We observed that the data may be distributed in such a way that in the ICA highly correlated independent components emerge, which do not maximally separate the symptom variance associate with neural variance. We demonstrate this by plotting the relationship between parcel beta coefficients for the *β*_*PC*3_*GBC* map versus the *β*_*IC*2_*GBC* and *β*_*IC*3_*GBC* maps (Appendix 1 - Fig. 13G). The sigmoidal shape of the distribution indicates an improvement in the Z-statistics for the *β*_*PC*3_*GBC* map relative to the *β*_*IC*2_*GBC* and *β*_*IC*3_*GBC* maps.

Next, we show that the PCA solution was highly ***stable*** when tested across sites, and k-fold cross-validations, and was ***reproducible*** in independent split-half samples. First, we show that the symptom-derived PCA solution remained highly robust across all sites (**Fig. 2A**) and 5-fold crossvalidation iterations (see **Methods**). Notably, the total proportion of variance explained as well as the total number of significant PCs remained stable (**Fig. 2A**). Second, PCA loadings accurately and reliably computed the scores of subjects in a hold-out sample (**Fig. 2B**). Specifically, for each left-out site, we tested if single-subject predicted scores were similar to the originally observed scores of these same subjects in the full PCA (i.e. performed with all 436 subjects). This similarity was high for all sites (**Appendix 1 - Fig. 6**). Third, we verified that the predicted-versus-observed PCA score similarities also hold via a *k*-fold cross-validation (for *k*=2 to 10, **Fig. 2C** & **Appendix 1 - Fig. 5**). Finally, across all 5 PCs, predicted-to-observed similarity of PCA loadings was remarkably high. Moreover, PCA loadings were stable when testing via leave-site-out analysis and 5-fold cross-validation. We furthermore demonstrated the reproducibility of the solution using 1,000 independent split-half replications (**Fig. 2D**). For each run of split-half replication, PSD subjects were split into 2 sets and a PCA was performed independently in both sets. The loadings from both PCAs were then compared using a Pearson’s correlation. Importantly, results were not driven by medication status or dosage (**Appendix 1 - Fig. 7**). Collectively, these data-reduction analyses strongly support a stable and reproducible low-rank PSD symptom geometry.

### Dimensionality-Reduced PSD Symptom Geometry Reveals Novel and Robust Neuro-Behavioral Relationships

Next, we tested if the dimensionality-reduced symptom geometry can identify robust and novel patterns of neural variation across PSD. We chose to use global brain connectivity (GBC), a summary FC metric, to measure neural variance because it yields a parsimonious measure reflecting how globally coupled an area is to the rest of the brain (***Cole et al., 2010***) (see **Methods**). Furthermore, we selected GBC because: i) the metric is agnostic regarding the location of dysconnectivity as it weights each area equally; ii) it yields an interpretable dimensionality-reduction of the full FC matrix; iii) unlike the full FC matrix or other abstracted measures, GBC produces a neural map, which can be related to other independent neural maps (e.g. gene expression or pharmacology maps, discussed below). Furthermore, GBC has been shown to be sensitive to altered patterns of global neural connectivity in PSD cohorts (***Anticevic et al., 2013a***; ***Fornito et al., 2011***; ***Hahamy et al., 2014***; ***Cole et al., 2011***) as well as in models of psychosis (***Preller et al., 2018***; ***Driesen et al., 2013***).

All 5 PCs captured unique patterns of GBC variation across the PSD (**Appendix 1 - Fig. 10**), which were not observed in CON (**Appendix 1 - Fig. 11**). Again we highlight the hallmark *“Psychosis Configuration”* dimension (i.e. *PC3*), here to illustrate the benefit of the low-rank PSD symptom geometry for symptom-neural mapping relative to traditional aggregate PANSS symptom scales. The relationship between total PANSS Positive scores and GBC across N=436 PSD patients (*β_Pos_GBC*, **Fig. 3A**) was statistically modest (**Fig. 3B**) and no areas survived whole-brain type-I error protection (**Fig. 3C**, *p*<0.05). In contrast, regressing *PC3* scores onto GBC across N=436 patients revealed a robust symptom-neural *β*_*PC*3_*GBC* map (**Fig. 3E-F**), which survived whole-brain type-I error protection. Of note, the *PC3 “Psychosis Configuration”* axis is bi-directional whereby individuals who score either highly positively or negatively are symptomatic. Therefore, a high positive *PC3* score was associated with both reduced GBC across insular and superior dorsal cingulate cortices, thalamus, and anterior cerebellum and elevated GBC across precuneus, medial prefrontal, inferior parietal, superior temporal cortices and posterior lateral cerebellum - consistent with the default-mode network (***Fox et al., 2005***). A high negative *PC3* score would exhibit the opposite pattern. Critically, this robust symptom-neural mapping emerged despite no differences between mean diagnostic group *PC3* scores (**Fig. 3D**). These two diverging directions may be captured separately in the ICA solution, when orthogonality is not enforced (**Appendix 1 - Fig. 13**). Moreover, the *PC3* symptom-neural map exhibited improved statistical properties relative to other GBC maps computed from traditional aggregate PANSS symptom scales (**Fig. 3G-J**). Notably, the *PC2 - Cognitive Functioning* dimension, which captured a substantial proportion of cognitive performance-related symptom variance independent of other symptom axes, revealed a circuit that was moderately (anti-)correlated with other PC maps but strongly anti-correlated with the BACS composite cognitive deficit map (*r*=-0.81, **Appendix 1 - Fig. 9O**). This implies that the *PC2* map reflects unique neural circuit variance that is relevant for cognition, independent of the other *PC* symptom dimensions.

**Figure 3.**
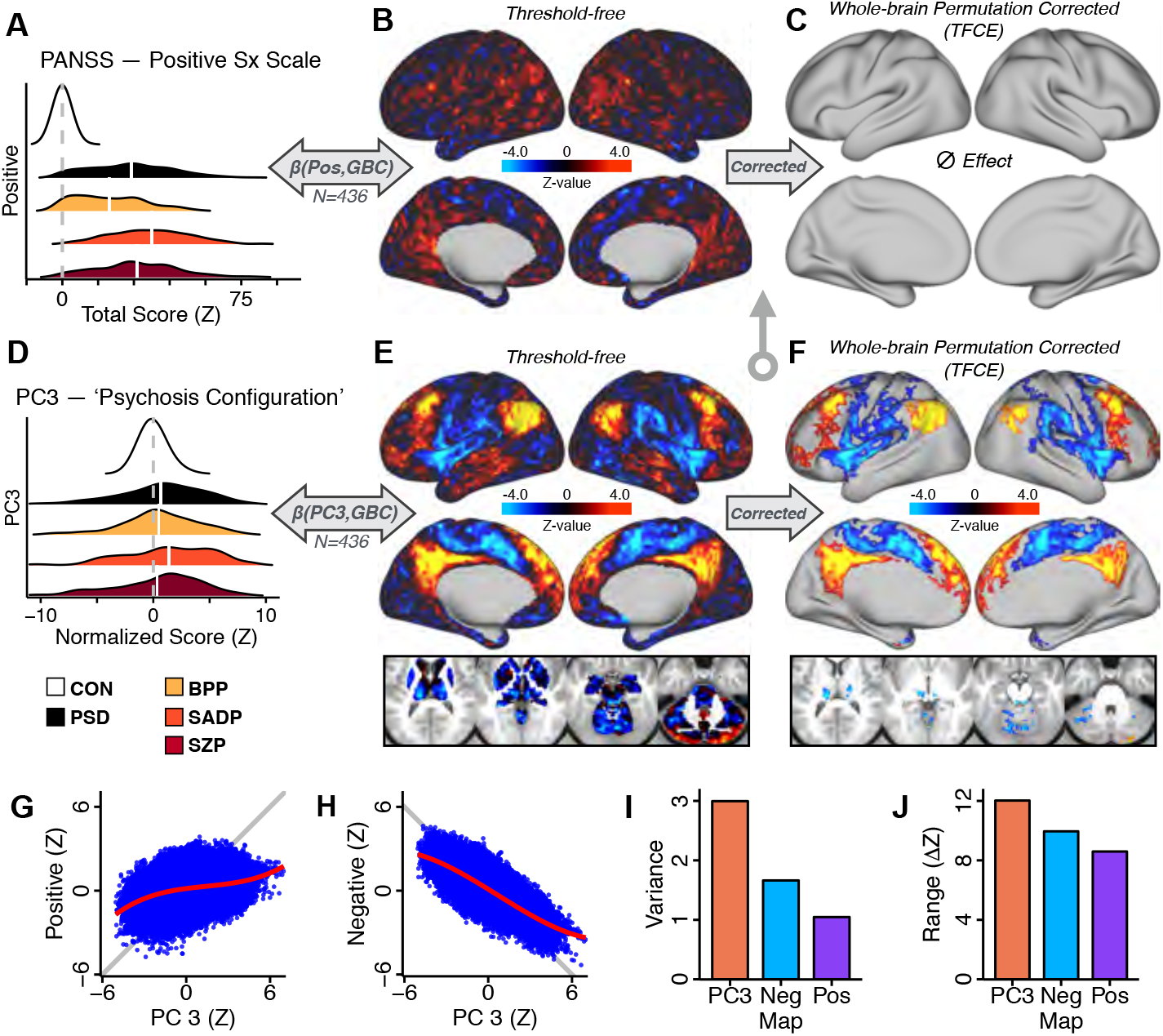
Dimensionality-reduced symptom variation reveals robust neuro-behavioral mapping. **(A)** Distributions of total PANSS Positive symptoms for each of the clinical diagnostic groups normalized relative to the control group (white=CON;black=all PSD patients;yellow=BPP;orange=SADP;red=SZP). **(B)** *β_Positive_GBC* map showing the relationship between the aggregate PANSS Positive symptom score for each patient regressed onto global brain connectivity (GBC) across all patients (N=436). **(C)** No regions survived non-parametric family-wise error (FWE) correction at p<0.05 using permutation testing with threshold-free cluster enhancement (TFCE). **(D)** Distributions of scores for *PC3 “Psychosis Configuration”* across clinical groups, again normalized to the control group. **(E)** *β*_*PC*3_*GBC* map showing the relationship between the *PC3 “Psychosis Configuration”* score for each patient regressed onto GBC across all patients (N=436). **(F)** Regions surviving p<0.05 FWE whole-brain correction via TFCE showed clear and robust effects. **(G)** Comparison between the *Psychosis Configuration* symptom score versus the aggregate PANSS Positive symptom score GBC map for every datapoint in the neural map (i.e. greyordinate in the CIFTI map). The sigmoidal pattern indicates an improvement in the Z-statistics for the *Psychosis Configuration* symptom score map (panel **E**) relative to the aggregate PANSS Positive symptom map (panel **B**). **(H)** A similar effect was observed when comparing the *Psychosis Configuration* GBC map relative to the PANSS Negative symptoms GBC map (**Appendix 1 - Fig. 9**). **(I)** Comparison of the variances for the *Psychosis Configuration,* PANSS Negative and PANSS Positive symptom map Z-scores. **(J)** Comparison of the ranges between the *Psychosis Configuration,* Negative and Positive symptom map Z-scores. Symptom-neural maps for all 5 PCs and all 4 traditional symptom scales (BACS and PANSS subscales) are shown in **Appendix 1 - Fig. 9**.

### Univariate Neuro-Behavioral Map of Psychosis Configuration is Reproducible

After observing improved symptom-neural *PC3* statistics, we tested if these neural maps replicate. Recent attempts to derive stable symptom-neural mapping using multivariate techniques, while inferentially valid, have not replicated (***Dinga et al., 2019***). This fundamentally relates to the tradeoff between the sample size needed to resolve multivariate neuro-behavioral solutions and the size of the feature space. To mitigate the feature size issue we re-computed the *β_PC_GBC* maps using a functionally-derived whole-brain parcellation via the recently-validated CAB-NP atlas (***Glasser et al., 2016***; ***Ji et al., 2019c***) (**Methods**). Here, a functional parcellation is a principled way of neural feature reduction (to 718 parcels) that can also appreciably boost signal-to-noise (***Glasser et al., 2016***; ***Ji et al., 2019c***). Indeed, parcellating the full-resolution “dense” resting-state signal for each subject prior to computing GBC statistically improved the group-level symptom-neural maps compared to parcellating after computing GBC (**Fig. 4A-C**, all maps in **Appendix 1 - Fig. 10**). Results demonstrate that the univariate symptom-neural mapping was indeed stable across 5-fold boot-strapping, leave-site-out, and split-half cross-validations (**Fig. 4D-N**, see **Appendix 1 - Note 1**), yielding consistent symptom-neural *PC3* maps. Importantly, the symptom-neural maps computed using ICA showed comparable results (**Appendix 1 - Fig. 13**).

**Figure 4.**
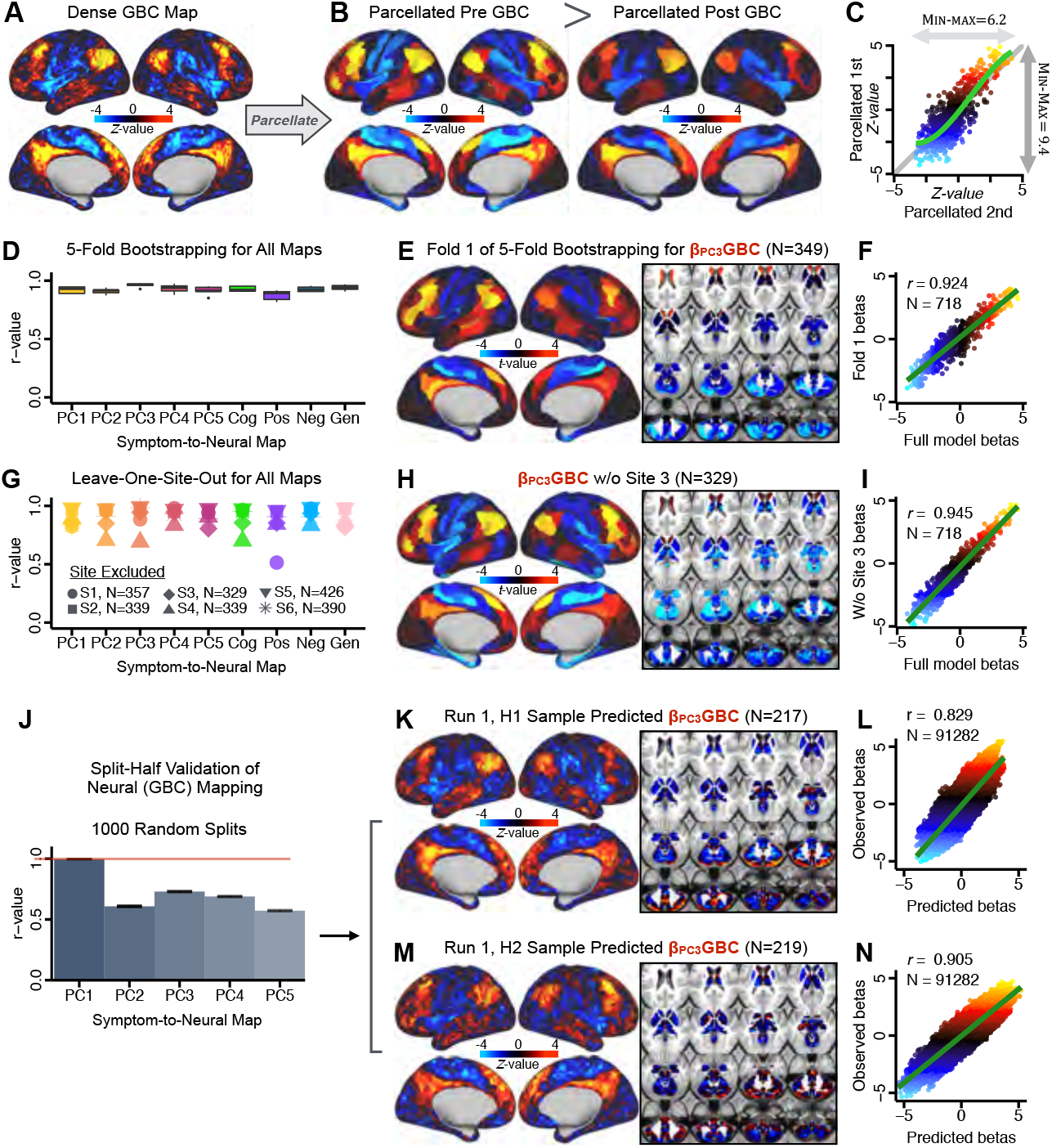
Parcellated symptom-neural GBC maps reflecting psychosis configuration are statistically robust and reproducible. **(A)** Z-scored *PC3 Psychosis Configuration* GBC neural map at the ‘dense’ (full CIFTI resolution) level. **(B, C)** Neural data parcellated using a whole-brain functional partition (***Ji et al., 2019c***) before computing subject-level GBC yielded stronger statistical values in the Z-scored *Psychosis Configuration* GBC neural map as compared to when parcellation was performed *after* computing GBC for each subject. **(D)** Summary of similarities between all symptom-neural *βGBC* maps (PCs and traditional symptom scales) across 5-fold cross-validation. Boxplots show the range of *r* values between *βGBC* maps for each fold and the full model. **(E)** Normalized *βGBC* map from regression of individual patients’ *PC3* scores onto parcellated GBC data, shown here for a subset of patients from Fold 1 out of 5 (N=349). The greater the magnitude of the coefficient for a parcel, the stronger the statistical relationship between GBC of that parcel and *PC3* score. **(F)** Correlation between the *βGBC* value of each parcel in the regression model computed using patients in Fold 1 and the full PSD sample (N=436) model indicates that the leave-one-fold-out *βGBC* map was highly similar to the *βGBC* map obtained from the full PSD sample model (*r*=0.924). **(G)** Summary of leave-one-site-out regression for all symptom-neural maps. Regression of PC symptom scores onto parcellated GBC data, each time leaving out subjects from one site, resulted in highly similar maps. This highlights that the relationship between *PC3* scores and GBC is robust and not driven by a specific site. **(H)** *βGBC* map for all PSD except one site. As an example, Site 3 is excluded here given that it recruited the most patients (and therefore may have the greatest statistical impact on the full model). **(I)** Correlation between the value of each parcel in the regression model computed using all patients minus Site 3, and the full PSD sample model. **(J)** Split-half replication of *β*_*PC*3_*GBC* map. Bar plots show the mean correlation across 1,000 runs;error bars show standard error. Note that the split-half effect for *PC1* was exceptionally robust. The split-half consistency for *PC3*, while lower, was still highly robust and well above chance. **(K)** *β*_*PC*3_*GBC* map from *PC3*-to-GBC regression for the first half (H1) patients, shown here for one exemplar run out of 1000 split-half validations. **(L)** Correlation across 718 parcels between the H1 predicted coefficient map (i.e. panel **K**) and the observed coefficient map for H1. **(M-N)** The same analysis as **K-L** is shown for patients in H2, indicating a striking consistency in the *Psychosis Configuration β*_*PC*3_*GBC* map correspondence.

### A Multivariate PSD Neuro-Behavioral Solution Can be Computed but is Not Reproducible with the Present Sample Size

Several recent studies have reported “latent” neuro-behavioral relationships using multivariate statistics (***Xia et al., 2018***; ***Drysdale et al., 2017***; ***Yu et al., 2019***), which would be preferable because they simultaneously solve for maximal covariation across neural and behavioral features. Given the possibility of deriving a stable multivariate effect, here we tested if results improve with canonical correlation analysis (CCA) (***Hardoon et al., 2004***) (**Fig. 5A**).

**Figure 5.**
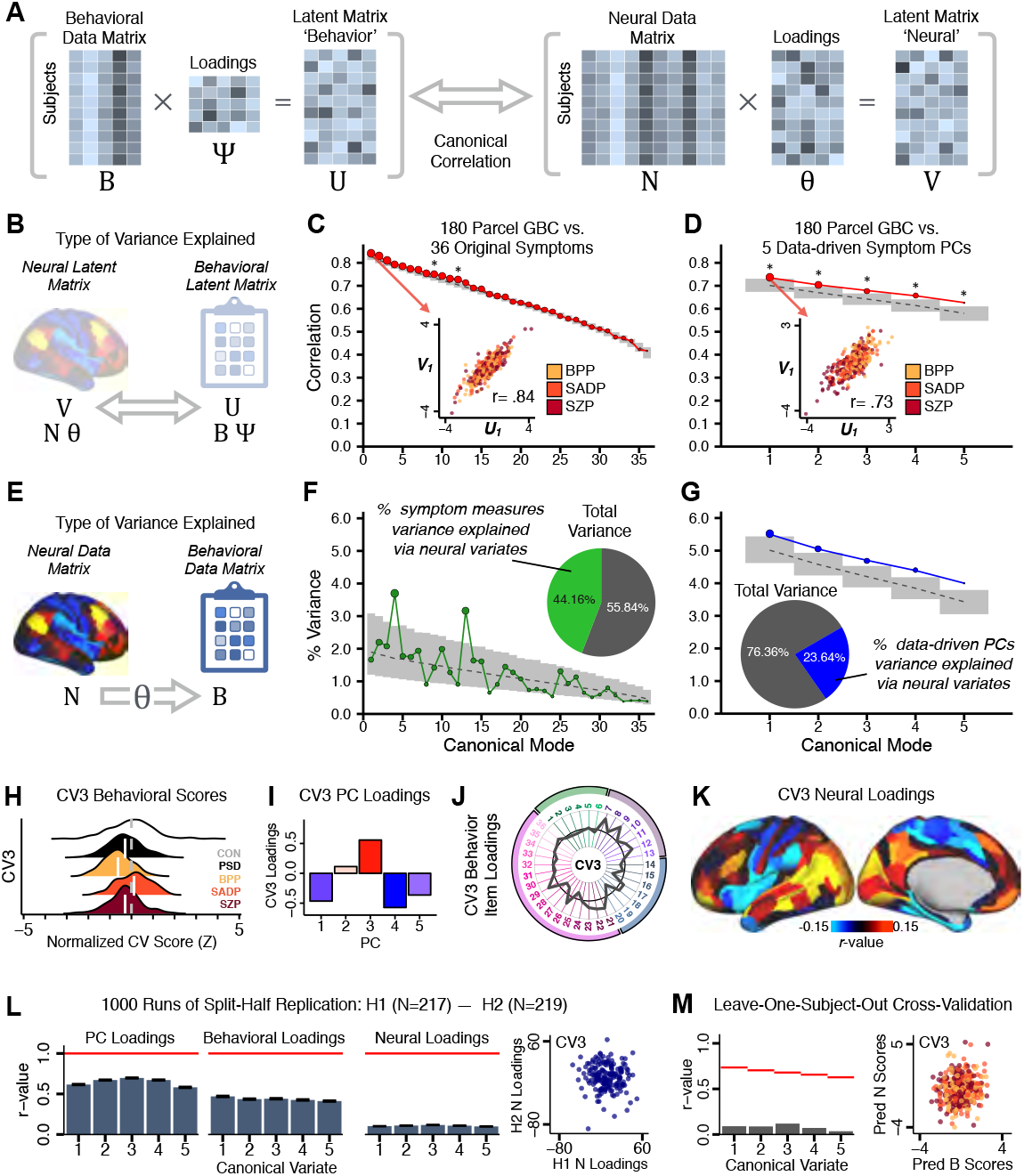
Multivariate symptom-neural feature mapping using canonical correlation analysis (CCA). **(A)** Schematic of CCA data (***B***, ***N***), transformation (*Ψ*, *Θ*), and transformed ‘latent’ (***U***, ***V***) matrices. Here each column in ***U*** and ***V*** is referred to as a canonical variate (CV);each corresponding pair of CVs (e.g. ***U_1_*** and ***V_1_***) is referred to as a canonical mode. **(B)** CCA maximized correlations between the CVs (***U*** and ***V***) **(C)** Screeplot showing canonical modes obtained from 180 neural features (cortical GBC symmetrized across hemispheres) and 36 symptom measures (‘180 vs. 36 CCA’). Inset illustrates the correlation (*r*=0.85) between the CV of the first mode, ***U_1_*** and ***V_1_*** (note that the correlation was not driven by a separation between diagnoses). Modes 9 and 12 remained significant after FDR correction. **(D)** CCA computed with 180 neural features and 5 PC symptom features (‘180 vs. 5 CCA’). Here all modes remained significant after FDR correction. Dashed black line shows the null calculated via a permutation test with 5,000 shuffles; grey bars show 95% confidence interval. **(E)** Correlation between ***B*** and ***NΘ*** reflects how much of the symptom variation can be explained by the latent neural features. **(F)** Proportion of symptom variance explained by each of the neural CVs in the 180 vs. 36 CCA. Inset shows the total proportion of behavioral variance explained by the neural CVs. **(G)** Proportion of total symptom variance explained by each of the neural CVs in the 180 vs. 5 CCA. While CCA using symptom PCs has fewer dimensions and thus accounts for lower total variance (see inset), each neural variate explains a higher amount of symptom variance than in **F**, suggesting that CCA could be optimized by first obtaining a low-rank symptom solution. Dashed black line indicates the null calculated via a permutation test with 5,000 shuffles;grey bars show 95% confidence interval. Neural variance explained by symptom CVs are plotted in **Appendix 1 - Fig. 16**. **(H)** Distributions of *CV3* scores from the 180 vs. 5 CCA are shown here as an example of characterizing CV configurations. Scores for all diagnostic groups are normalized to CON. Additionally, **(I)** symptom canonical factor loadings, **(J)** loadings of the original 36 symptom measures, and **(K)** neural canonical factor loadings for *CV3* are shown. **(L)** Within-sample CCA cross-validation appeared robust (see **Appendix 1 - Fig. 18**). However, a split-half replication of the 180 vs. 5 CCA (using two independent non-overlapping samples) was not reliable. Bar plots show the mean correlation for each CV between the first half (H1) and the second half (H2) CCA, each computed across 1,000 runs. *Left:* split-half replication of the symptom PC loadings matrix *Ψ*; *Middle:* individual symptom measure loadings; *Right:* the neural loadings matrix *Θ*, which in particular was not stable. Error bars show the standard error of the mean. Scatterplot shows the correlation between *CV3* neural loadings for H1 vs. H2 for one example CCA run, illustrating lack of reliability. **(M)** Leave-one-subject-out cross-validation further highlights CCA instability.

To evaluate if the number of neural features affects the solution, we computed CCA using GBC from: i) 180 symmetrized cortical parcels; ii) 359 bilateral subcortex-only parcels; iii) 192 symmetrized subcortical parcels; iv) 12 functional networks, all from the brain-wide cortico-subcortical CAB-NP parcellation (***Glasser et al., 2016***; ***Ji et al., 2019c***). We did not compute a solution using 718 bilateral whole-brain parcels, as this exceeded the number of samples and rendered the CCA insolvable. Notably, the 359 subcortex-only solution did not produce stable results according to any criterion, whereas the 192 symmetrized subcortical features (**Appendix 1 - Fig. 14**) and 12 networklevel features (**Appendix 1 - Fig. 15**) solutions captured statistically modest effects relative to the 180 symmetrized cortical features (**Fig. 5B-D**). Therefore, we characterized the 180-parcel solution further. We examined two CCA solutions using these 180 neural features in relation to symptom scores: i) all 36 item-level symptom measure scores from the PANSS/BACS (‘180 vs. 36 CCA’, **Fig. 5C&F**); and ii) 5 PC symptom scores (‘180 vs. 5 CCA’, **Fig. 5D&G**, see **Methods**). Only 2 out of 36 modes for the 180 vs. 36 CCA solution survived permutation testing and false discovery rate (FDR) correction (**Fig. 5C**). In contrast, all 5 modes of the 180 vs. 5 CCA survived (**Fig. 5D**). Critically, we found that no single CCA mode computed on item-level symptom measures captured more variance than the CCA modes derived from PC symptom scores, suggesting that the PCA-derived dimensions capture more neurally-relevant variation than anyone single clinical item (**Fig. 5E-G**). Additional CCA details are presented in **Appendix 1 - Note 1** and **Appendix 1 - Fig. 16**.

We highlight here an example canonical variate, *CV3*, across both symptom and neural effects, with data for all *CVs* shown in **Appendix 1 - Fig. 17**. *CV3* scores across diagnostic groups normalized to controls are shown **Fig. 5H** in addition to how *CV3* loads onto each *PC* (**Fig. 5I**). The negative loadings on to *PCs 1, 4, and 5* and the high positive loadings on to *PC3* in **Fig. 5I** indicate that *CV3* captures some shared variation across symptom *PCs*. This can also be visualized by computing how *CV3* projects onto the original 36 symptom measures (**Fig. 5J**). Finally, the neural loadings for *CV3* are shown in **Fig. 5K**.

Lastly, we tested if the 180 vs. 5 CCA solution is stable and reproducible, as done with PC-to-GBC univariate results. The CCA solution was robust when tested with *k*-fold and leave-site-out crossvalidation (**Appendix 1 - Fig. 18**) likely because these methods use CCA loadings derived from the full sample. However, the CCA loadings did not replicate in non-overlapping split-half samples (**Fig. 5L**, see **see Appendix 1 - Note 1**). Moreover, a leave-one-subject-out cross-validation revealed that removing a single subject from the sample affected the CCA solution such that it did not generalize to the left-out subject (**Fig. 5M**). This is in contrast to the PCA-to-GBC univariate mapping, which was substantially more reproducible for all attempted cross-validations relative to the CCA approach. This is likely because substantially more power is needed to resolve a stable multivariate neuro-behavioral effect with this many features (***Dinga et al., 2019***). Indeed, a multivariate power analysis using 180 neural features and 5 symptom features and assuming a true canonical correlation of *r* = 0.3 suggests that a minimal sample size of *N* = 8,145 is needed to sufficiently detect the effect (***Helmer et al., 2020***), which is an order of magnitude greater than the available sample size (**Appendix 1 - Fig. 19**). Therefore, we leverage the univariate symptom-neural result for subsequent subject-specific model optimization and comparisons to molecular neuroimaging maps.

### A Major Proportion of Overall Neural Variance May Not be Relevant for Psychosis Symptoms

Most studies look for differences between clinical and control groups, but to our knowledge no study has tested whether both PSD and healthy controls actually share a major portion of neural variance that may be present across all people. If the bulk of the neural variance is similar across both PSD and CON groups then including this clinically-irrelevant neural signal might obscure clinical-relevant neuro-behavioral relationships. To test this, we examined the shared variance structure of the neural signal for all PSD patients (N=436) and all controls (N=202) independently by conducting a PCA on the GBC maps (see **Methods**). Patients’ and controls’ neural signals were highly similar for each of the first 3 neural PCs (>30% of all neural variance in each group)(**Appendix 1 - Fig. 20A-J**). These neural *PCs* may reflect a “core” shared symptom-irrelevant neural variance that generalizes across all people. These data suggest that the bulk of neural variance in PSD patients may actually not be symptom-relevant, which highlights the importance of optimizing symptom-neural mapping. Under this assumption, removing the shared neural variance between PSD and CON should not drastically affect the reported symptom-neural univariate mapping solution, because this common variance *does not map to clinical features.* Thus, we conducted a PCA using the parcellated GBC data from all 436 PSD and 202 CON (see **Methods**), which we will refer to as “GBC-PCA” to avoid confusion with the symptom/behavioral PCA. This GBC-PCA resulted in 637 independent GBC-PCs. Since PCs are orthogonal to each other, we then partialled out the variance attributable to GBC-PC1 from the PSD data by reconstructing the PSD GBC matrix using only scores and coefficients from the remaining 636 GBC-PCs 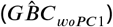. We then reran the univariate regression as described in **Fig. 3**, using the same five symptom PC scores across 436 PSD (**Appendix 1 - Fig. 21**). Removing the first PC of shared neural variance (which accounted for about 15.8% of the total GBC variance across CON and PSD) from PSD data attenuated the statistics slightly (not unexpected as the variance was by definition reduced) but otherwise did not strongly affect the univariate mapping solution. We repeated the symptom-neural regression next with the first two GBC-PCs partialled out of the PSD data **Appendix 1 - Fig. 22**, with the first three PCs parsed out **Appendix 1 - Fig. 23**, and with the first four neural PCs parsed out **Appendix 1 - Fig. 24**. The symptom-neural maps remain fairly robust, although the similarity with the original *β_PC_GBC* maps does drop as more common neural variance is parsed out.

### Optimizing Symptom-Neural Mapping Features for Personalized Prediction via Dimensionality-reduced Symptoms

Above we demonstrated that symptom *PC* scores can be reliably computed across sites and cross-validation approaches (**Fig. 2**). Next we show that leave-one-subject-out cross-validation yields reliable effects for the low-rank symptom PCA solution (**Fig. 6A**). This stable single-subject PC score prediction provides the basis for testing if the derived neural maps can yield an individually-reliable subset of features. To this end, we developed a framework for univariate neural feature selection (**Appendix 1 - Fig. 25**) based on PC scores (e.g. *PC3* score). Specifically, we computed a dot product GBC metric (*dpGBC*) that provides an index of similarity between an individual *ΔGBC* topography relative to a “reference” group-level *β_PC_GBC* map (see **Methods** and **Appendix 1 - Fig. 25**). Using this *dpGBC* index via a feature selection step-down regression, we found a subset of parcels for which the symptom-neural statistical association was maximal (**Fig. 6A**). For *PC3* we found *P* = 39 maximally predictive parcels out of the group neural *β_PC_GBC* map. Specifically, the relationship between *PC3* symptom scores and *dpGBC* values across subjects was maximal (**Fig. 6B**, top panel, *r* = 0.36) as was the relationship between predicted *dpGBC^pred^* vs. observed *dpGBC^obs^* (**Fig. 6B**, bottom panel, *r* = 0.35) (see **Appendix 1 - Fig. 26** for all PCs). Importantly, the “subset” feature map (i.e. [*β*_*PC*3_*GBC*^*obs*^]^*P*=39^, **Fig. 6C**) exhibited improved statistical properties relative to the full map (i.e. [*β*_*PC*3_*GBC*^*obs*^]^*P*=718;^, **Fig. 6D**). Furthermore, the relationship between observed vs. predicted subset feature maps (i.e. *r*[*dpGBC^obs^, dpGBC^pred^*]) was highly consistent across DSM diagnoses (**Fig. 6E**). Finally, a single patient is highlighted for whom the correlation between predicted and observed subset feature maps was high (i.e. *r*[*ΔGBC^pred^, ΔGBC^obs^*], **Fig. 6F**), demonstrating that the dimensionality-reduced symptom scores can be used to quantitatively optimize symptom-neural map features for individuals.

**Figure 6.**
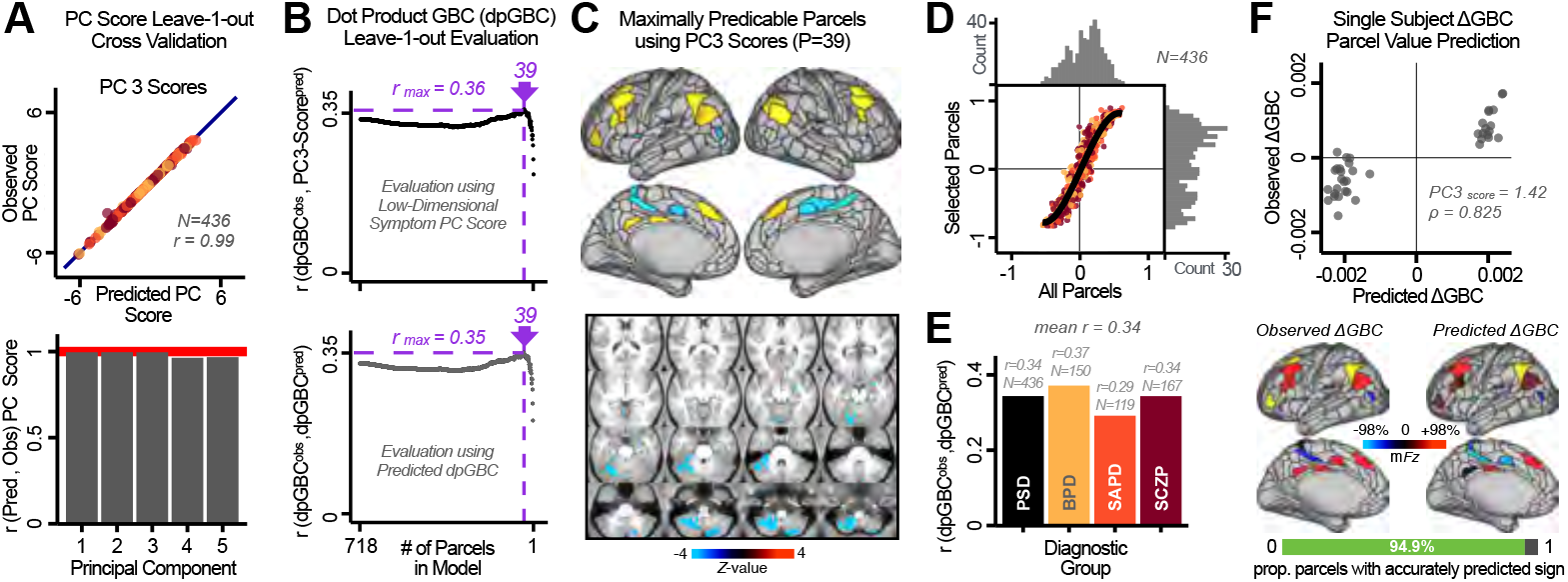
Optimizing neural feature selection to inform single-subject prediction via a low-dimensional symptom solution. **(A)** Leave-one-out cross-validation for the symptom PCA analyses indicates robust individual score prediction. Top panel: Scatterplot shows the correlation between each subject’s predicted *PC3* score from a leave-one-out PCA model and their observed *PC3* score from the full-sample PCA model, *r*=0.99. Bottom panel: Correlation between predicted and observed individual PC scores was above 0.99 for each of the significant PCs (see **Fig. 1**). The red line indicates *r*=1. **(B)** We developed a univariate step-down feature selection framework to obtain the most predictive parcels using a subject-specific approach via the *dpGBC* index. Specifically, the ‘observed’ patient-specific *dpGBC^obs^* was calculated using each patient’s *ΔGBC^obs^* (i.e. the patient-specific GBC map vs. the group mean GBC for each each parcel) and the ‘reference’ symptom-to-GBC *PC3* map (described in **Fig. 4B**) [*dpGBC^obs^* = *ΔGBC^obs^* · *β*_*PC*3_*GBC*^*obs*^]. See **Methods & Appendix 1 - Fig. 25** for complete feature selection procedure details. In turn, we computed the predicted *dpGBC* index for each patient by holding their data out of the model and predicting their score (*dpGBC^pred^*). We used two metrics to evaluate the maximally predictive feature subset: i) The correlation between *PC3* symptom score and *dpGBC^obs^* across all N=436, which was maximal for *P* = *39* parcels [*r*=0.36, purple arrow]; ii) The correlation between *dpGBC^obs^* and *dpGBC^pred^*, which also peaked at *P* = *39* parcels [*r*=0.35, purple arrow]. (C) The *P* = *39* maximally predictive parcels from the *β*_*PC*3_*GBC*^*obs*^ map are highlighted (referred to as the “selected” map). (D) Across all n=436 patients we evaluated if the selected parcels improve the statistical range of similarities between the *ΔGBC^obs^* and the *β*_*PC*3_*GBC*^*obs*^ reference for each patient. For each subject the value on the X-axis reflects a correlation between their *ΔGBC^obs^* map and the *β*_*PC*3_*GBC*^*obs*^ map across all 718 parcels;the Y-axis reflects a correlation between their *ΔGBC^obs^* map and the *β*_*PC*3_*GBC*^*obs*^ map only within the “selected” 39 parcels. The marginal histograms show the distribution of these values across subjects. (E) Each DSM diagnostic group showed comparable correlations between predicted and observed *dpGBC* values. The *r*-value shown for each group is a correlation between the *dpGBC^obs^* and *dpGBC^pred^* vectors, each of length N. (F) Scatterplot for a single patient with a positive behavioral loading (*PC*3 score= 1.42) and also with a high correlation between predicted *ΔGBC^pred^* versus observed *ΔGBC^obs^* values for the “selected” 39 parcels (*ρ* = 0.825). Right panel highlights the observed vs. predicted *ΔGBC* map for this patient, indicating that 94.9% of the parcels were predicted in the correct direction (i.e. in the correct quadrant).

### Single Patient Evaluation via Neuro-Behavioral Target Map Similarity

We showed a quantitative framework for computing a neuro-behavioral model at the single-patient level. This brain-behavior space (BBS) model was optimized along a single dimensionality-reduced symptom PC axis. Next, we tested a hybrid patient selection strategy by first imposing a PC-based symptom threshold, followed by a target neural similarity threshold driven by the most highly pre-dictive symptom-neural map features. This is described in **Methods**, **Appendix 1 - Note 1**, and **Appendix 1 - Fig. 27-28**. We found that for patients with a high (either positive or negative) *PC* symptom score the symptom-neural relationship was robust (**Appendix 1 - Fig. 27A-D**), and these patients could be reliably selected for that particular dimension (**Appendix 1 - Fig. 27E,G**). Conversely, patients with a low absolute *PC* score showed a weak relationship with symptom-relevant neural features. This is intuitive because these individuals do not vary along the selected PC symptom axis. We also tested this patient selection strategy on an independent cross-diagnostic “replication” sample, which yielded consistent results (**Appendix 1 - Fig. 27F,H**). Collectively, these results show that data-driven symptom scores can pinpoint individual patients for whom neural variation strongly maps onto a target neural reference map. These data also highlight that both symptom and neural information for an independent patient can be quantified in the reference “discovery” BBS using their symptom data alone.

### Subject-Specific PSD Neuro-Behavioral Features Track Neuropharmacological Map Patterns

Next, we use the personalized BBS selection in a proof-of-concept framework for informing molecular mechanism of possible treatment response by relating subject-specific BBS to independently-acquired pharmacological neuroimaging maps. Here we examine two mechanisms implicated in PSD neuropathology via ketamine, a N-methyl-D-aspartate (NMDA) receptor antagonist (***Krystal et al., 1994***), and lysergic acid diethylamide (LSD), primarily a serotonin receptor agonist (***Preller et al., 2018***; ***González-Maeso et al., 2007***; ***Egan et al., 1998***). We first quantified individual subjects’ BBS “locations” in the established reference neuro-behavioral geometry. The radarplot in **Fig. 7A** shows original symptoms whereas **Fig. 7B** shows *ΔGBC^obs^* maps for two patients from the replication dataset (denoted here with *X*_*PC*3_ and *Y*_*PC*3_, see **Appendix 1 - Fig. 12** for other ex-ample patients). Both of these patients exceeded the neural and behavioral BBS selection indices for *PC3* (defined independently in the “discovery” dataset, **Appendix 1 - Fig. 27C**). Furthermore, both patients exhibited neuro-behavioral variation in line with their expected locations in the BBS geometry. Specifically, patient *X*_*PC*3_ from the replication dataset scored highly negatively on the *PC3* axis defined in the “discovery” PSD sample (**Fig. 7A**). In contrast, patient *Y*_*PC*3_ scored positively on the *PC3* axis. Importantly, the correlation between the *ΔGBC^obs^* map for each patient and the group-reference *β*_*PC*3_*GBC* was directionally consistent with their symptom PC score **Fig. 7B-C**).

**Figure 7.**
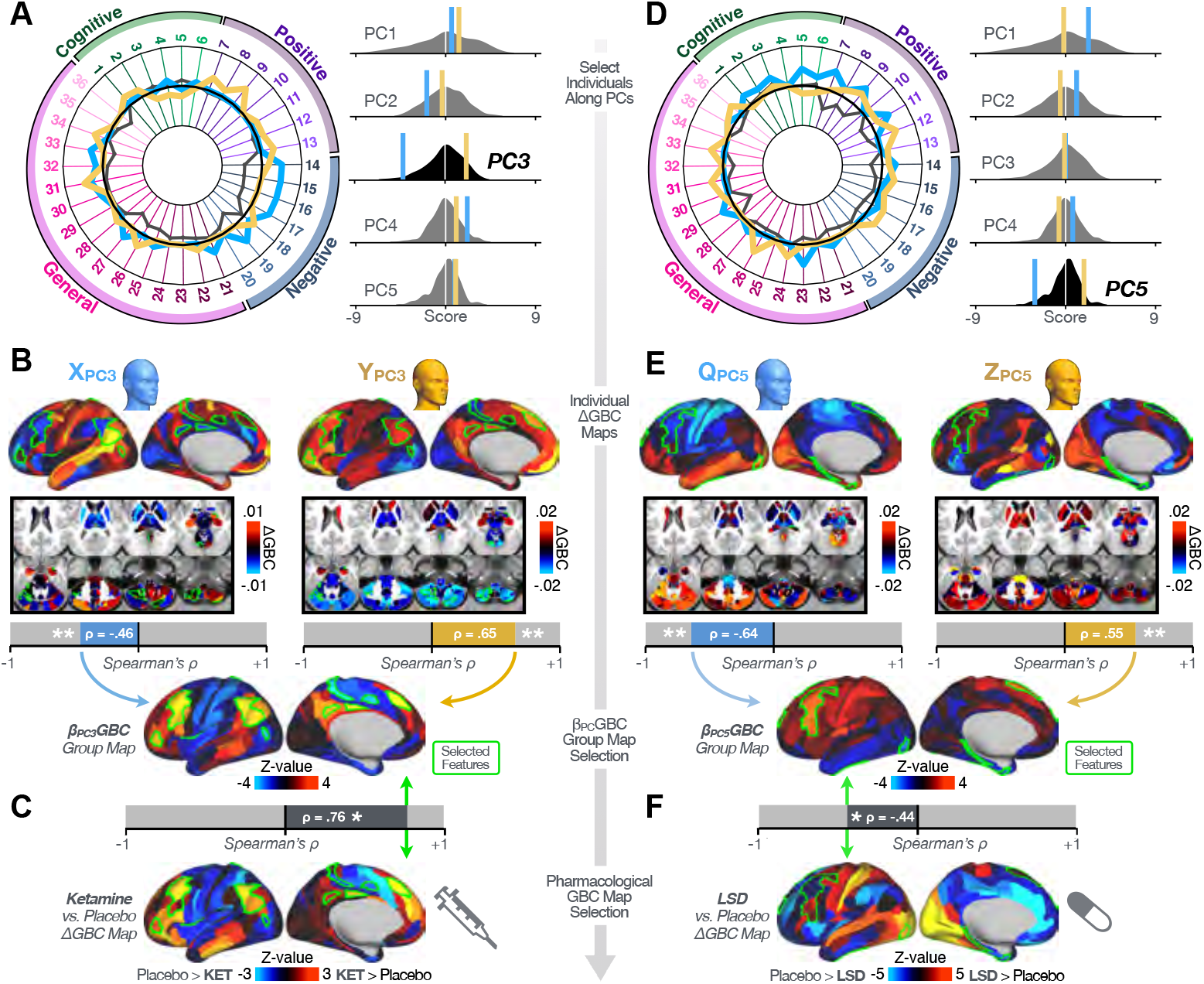
Leveraging subject-specific brain-behavioral maps for molecular neuroimaging target selection. **(A)** Data for two individual patients from the replication dataset are highlighted for PC3: *X*_*PC*3_ (blue) and *Y*_*PC*3_ (yellow). Both of these patients scored above the neural and behavioral thresholds for *PC3* defined in the “discovery” PSD dataset. Patient *X*_*P*3_ loads highly negatively on the *PC5* axis and Patient *Y*_*PC*3_ loads highly positively. Density plots show the projected PC scores for Patients *X*_*PC*3_ and *Y*_*PC*3_ overlaid on distributions of PC scores from the discovery PSD sample. **(B)** Neural maps show cortical and subcortical *ΔGBC^obs^* for the two patients *X*_*PC*3_ and *Y*_*PC*3_ specifically reflecting a difference from the mean *PC3*. The similarity of *ΔGBC^obs^* and the *β*_*PC*3_*GBC^obs^* map within the most predictive neural parcels for PC3 (outlined in green). Note that the sign of neural similarity to the reference *PC3* map and the sign of the *PC3* score is consistent for these two patients. **(C)** The selected *PC3* map (parcels outlined in green) is spatially correlated to the neural map reflecting the change in GBC after ketamine administration (*ρ*=0.76, **Methods**). Note that Patient *X*_*PC*3_ who exhibits *ΔGBC^obs^* that is anti-correlated to the ketamine map also expresses depressive moods symptoms (panel **A**). This is consistent with the possibility that this person may clinically benefit from ketamine administration, which may elevate connectivity in areas where they show reductions (***Berman et al., 2000***). In contrast, Patient *Y*_*PC*3_ may exhibit an exacerbation of their psychosis symptoms given that their *ΔGBC^obs^* is positively correlation with the ketamine map. **(D)** Data for two individual patients from the discovery dataset are highlighted for PC5: *Q*_*PC*5_ (blue) and *Z*_*PC*5_ (yellow). Note that no patients in the replication dataset were selected for *PC5* so both of these patients were selected from “discovery” PSD dataset for illustrative purposes. Patient *Q*_*PC*5_ loads highly negatively on the *PC5* axis and Patient *Z*_*PC*5_ loads highly positively. Density plots show the projected PC scores for Patients *Q*_*PC*5_ and *Z*_*PC*5_ overlaid on distributions of PC scores from the discovery PSD sample. **(E)** Neural maps show cortical and subcortical *ΔGBC^obs^* for Patients *Q*_*PC*5_ and *Z*_*PC*5_, which are highly negatively and positively correlated with the selected *PC5* map respectively. **(F)** The selected *PC5* map (parcels outlined in green) is spatially anti-correlated with the LSD response map (*ρ*=-0.44, see **Methods**), suggesting that circuits modulated by LSD (i.e. serotonin, in particular 5-HT2A) may be relevant for the *PC5* symptom expression. Here a serotonin receptor agonist may modulate the symptom-neural profile of Patient *Q*_*PC*5_, whereas an antagonist may be effective for Patient *Z*_*PC*5_.

We then tested if the single-subject BBS selection could be quantified with respect to a neural map reflecting glutamate receptor manipulation, a hypothesized mechanism underlying PSD symptoms (***Moghaddam and Javitt, 2012***). Specifically, we used an independently collected ke-tamine infusion dataset, collected in healthy adult volunteers during resting-state fMRI (***Anticevic et al., 2012b***). As with the clinical data, here we computed a *ΔGBC* map reflecting the effect of ketamine on GBC relative to placebo (**Methods**). The maximally-predictive *PC3* parcels exhibited high spatial similarity with the ketamine map (*p*=0.76, see **Methods**), indicating that the *ΔGBC* pattern induced by ketamine tracks with the GBC pattern reflecting *PC3* symptom variation.

Critically, because *X*_*PC*3_ is negatively loaded on the *PC3* symptom axis, an NMDA receptor antagonist like ketamine may modulate symptom-relevant circuits in a way that reduces similarity with the *PC3* map. This may in turn have an impact on the *PC3*-like symptoms. Consistent with this hypothesis, *X*_*PC*3_ expresses predominantly depressive symptoms (**Fig. 7A**), and ketamine has been shown to act as an anti-depressant (***Berman et al., 2000***). This approach can be applied for patients that load along another axis, such as *PC5*. **Fig. 7D-E** shows the symptom and neural data for two patients whom met thresholds for *PC5* selection (**Appendix 1 - Fig. 27C**). Notably, the selected *PC5* map is anti-correlated with a *ΔGBC* map reflecting LSD vs. placebo effects (***Preller et al., 2018***) (*p*=-0.44, **Fig. 7F**). Hence areas modulated by LSD may map onto behavioral variation along *PC5*. Consequently, serotonergic modulation may be effective for treating *Q*_*PC*5_ and *Z*_*PC*5_, via an antagonist or an agonist respectively. These differential similarities between pharmacological response maps and BBS maps (**Appendix 1 - Fig. 30**) can be refined for quantitative patient segmentation.

### Group-Level PSD Neuro-Behavioral Features Track Neural Gene Expression Patterns

To further inform molecular mechanism for the derived BBS results, we compared results with neural gene expression profiles derived from the Allen Human Brain Atlas (AHBA) (***Hawrylycz et al., 2012***; ***Burt et al., 2018***)(**Fig. 8A**, **Methods**). Specifically, we tested if BBS cortical topographies, which reflect stable symptom-neural mapping along PSD, covary with the expression of genes implicated in PSD neuropathology. We focus here on serotonin receptor subunits (HTR1E, HTR2C, HTR2A), GABA receptor subunits (GABRA1, GABRA5), and the interneuron markers somatostatin (SST) and parvalbumin (PVALB). Serotonin agonists such as LSD have been shown to induce PSD-like symptoms in healthy adults (***Preller et al., 2018***) and the serotonin antagonism of “second-generation” antipsychotics are thought to contribute to their efficacy in targeting broad PSD symptoms (***Geyer and Vollenweider, 2008***; ***Y Meltzer et al., 2012***; ***Abi-Dargham et al., 1997***). Abnormalities in GABAergic interneurons, which provide inhibitory control in neural circuits, may contribute to cognitive deficits in PSD (***Benes and Berretta, 2001***; ***Inan et al., 2013***; ***Dienel and Lewis, 2019***) and additionally lead to downstream excitatory dysfunction that underlies other PSD symptoms (***Lisman et al., 2008***; ***Grace, 2016***). In particular, a loss of prefrontal parvalbumin-expression fast-spiking interneurons has been implicated in PSD (***Enwright III et al., 2018***; ***Lodge et al., 2009***; ***Beasley and Reynolds, 1997***; ***Lewis et al., 2012***). **Fig. 8B** shows the distribution of correlations between the *PC3* map and the cortical expression patterns of 20,200 available AHBA genes (results for other PCs are shown in **Appendix 1 - Fig. 31**). Seven genes of interest are highlighted, along with their cortical expression topographies and their similarity with the *PC3* BBS map (**Fig. 8C-E**). This BBS-to-gene mapping can potentially reveal novel therapeutic molecular targets for neuro-behavioral variation. For example, the HTR1E gene, which encodes the serotonin 5-HT_1E_ receptor, is highly correlated with the *PC3* BBS map. This could drive further development of novel selective ligands for this receptor, which are not currently available (***Kitson, 2007***).

**Figure 8.**
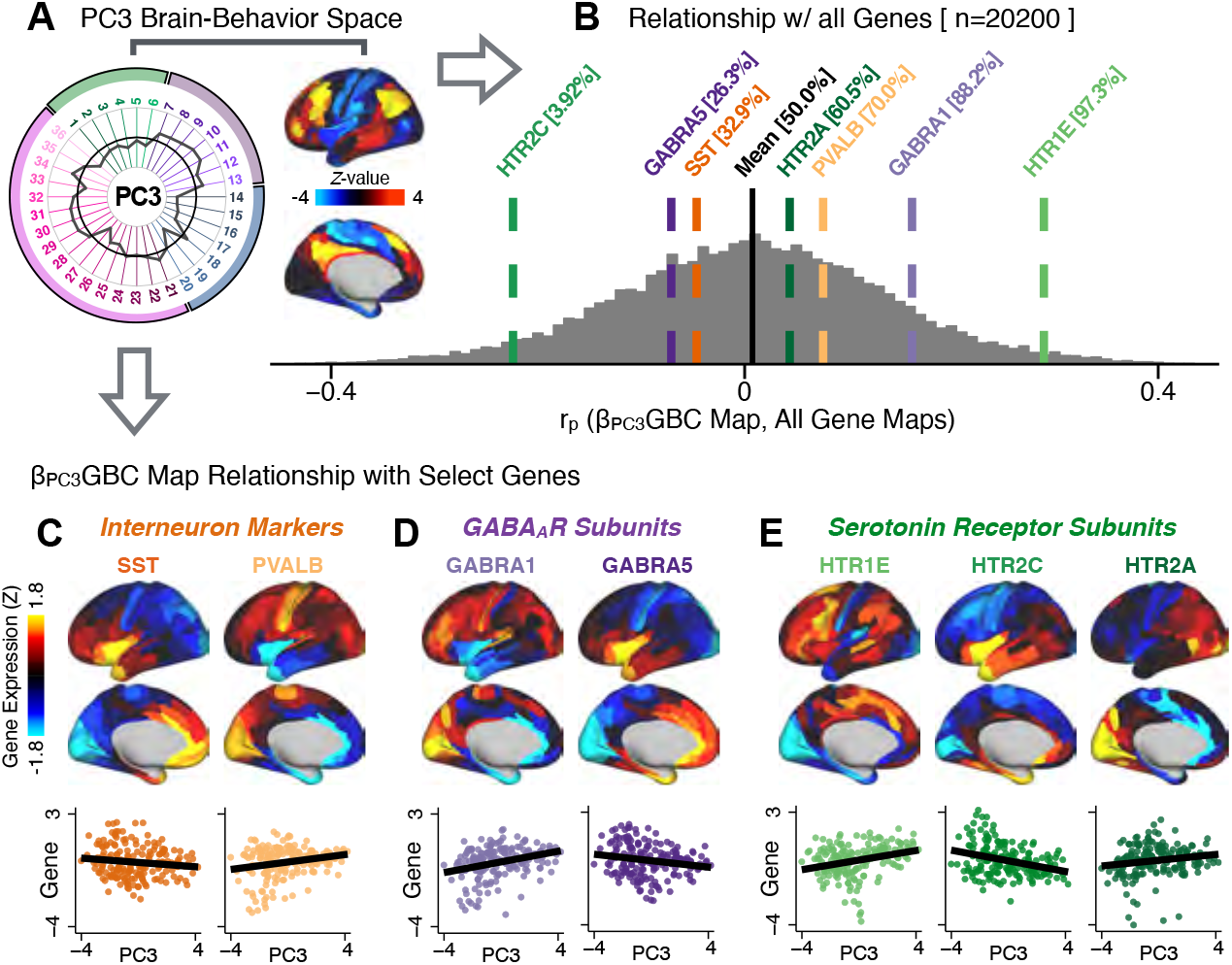
Psychosis spectrum symptom-neural maps track neural gene expression patterns computed from the Allen Human Brain Atlas (AHBA). **(A)** The symptom loadings and the associated neural map jointly reflect the *PC3* brain-behavioral space (BBS) profile, which can be quantitatively related to human cortical gene expression patterns obtained from the AHBA (***Burt et al., 2018***). **(B)** Distribution of correlation values between the *PC3* BBS map and ~20,000 gene expression maps derived from the AHBA dataset. Specifically, AHBA gene expression maps were obtained using DNA microarrays from six postmortem brains, capturing gene expression topography across cortical areas. These expression patterns were then mapped onto the cortical surface models derived from the AHBA subjects’ anatomical scans and aligned with the Human Connectome Project (HCP) atlas, described in prior work and methods (***Burt et al., 2018***). Note that because no significant inter-hemispheric differences were found in cortical gene expression all results were symmetrized to the left hemisphere, resulting in 180 parcels. We focused on a select number of psychosis-relevant genes - namely genes coding for the serotonin and GABA receptor subunits and interneuron markers. Seven genes of interest are highlighted with dashed lines. Note that the expression pattern of HTR2C (green dashed line) is at the low negative tail of the entire distribution, i.e. highly anti-correlated with *PC3* BBS map. Conversely, GABRA1 and HTR1E are on the far positive end, reflecting a highly similar gene-to-BBS spatial pattern. **(C)** Upper panels show gene expression patterns for two interneuron marker genes, somatostatin (SST) and parvalbumin (PVALB). Positive (yellow) regions show areas where the gene of interest is highly expressed, whereas negative (blue) regions indicate low expression values. Lower panels highlight all gene-to-BBS map spatial correlations where each value is a symmetrized cortical parcel (180 in total) from the HCP atlas parcellation. **(D)** Gene expression maps and spatial correlations with the *PC3* BBS map for two GABAA receptor subunit genes: GABRA1 and GABRA5. **(E)** Gene expression maps and spatial correlations with the *PC3* BBS map for three serotonin receptor subunit genes: HTR1E, HTR2C, and HTR2A.

## Discussion

We found a robust and reproducible symptom-neural mapping across the psychosis spectrum that emerged from a low-dimensional symptom solution. Critically, this low-rank symptom solution was predictive of a neural circuit pattern, which reproduced at the single-subject level. In turn, we show that the derived PSD symptom-neural feature maps exhibit spatial correspondence with independent pharmacological and gene expression neural maps that are directly relevant for PSD neurobiology. We demonstrate, for the first time in the literature, a transdiagnostic data-reduced PSD symptom geometry across hallmark psychopathology symptoms and cognitive measures, that shows stable and robust relationships with neural connectivity with singles subject-level precision. Critically, we anchor these symptom-neural maps to mechanistically-informed molecular imaging benchmarks that can be linked to relevant pathophysiology, and furthermore demonstrate how these methods can be combined to inform personalized patient selection decisions.

### Deriving an Individually Predictive Low-Dimensional Symptom Representation Across the Psychosis Spectrum

Psychosis spectrum is associated with notable clinical heterogeneity such deficits in cognition as well as altered beliefs (i.e. delusions), perception (i.e. hallucinations), and affect (i.e. negative symptoms) (***Lefort-Besnard et al., 2018***). This heterogeneity is captured by clinical instruments that quantify PSD symptoms across dozens of specific questions and ratings. This yields a high-dimensional symptom space that is intractable for reliable mapping of neural circuits (***Helmer et al., 2020***). Here we show that a low-rank solution captures principal axes of PSD symptom variation, a finding in line with prior work in SZP (***van der Gaag et al., 2006a***; ***Lindenmayer et al., 1994***; ***Emsley et al., 2003***; ***White et al., 1997***; ***Dollfus et al., 1996***; ***Blanchard and Cohen, 2006***; ***Chen et al., 2019***; ***Lefort-Besnard et al., 2018***).

These results highlights two key observations: i) Existing symptom reduction studies (even those in SZP specifically) have not evaluated solutions that include cognitive impairment - a hallmark deficit across the psychosis spectrum (***Barch et al., 2013***). Here we show that cognitive performance captures a notable portion of the symptom variance independent of other axes. We observed that cognitive variation captured 10% of PSD sample variance even after accounting for “Global Functioning” psychopathology. ii) No study has quantified cognitive deficit variation along with core psychosis symptoms via dimensionality reduction across multiple PSD diagnoses. While existing studies have evaluated stability of data-reduced solutions within a single DSM category (***van der Gaag et al., 2006a***,b; ***Lindenmayer et al., 1995***; ***Lefort-Besnard et al., 2018***), current results show that dimensionality-reduced PSD symptom solutions can be reproducibly obtained across DSM diagnoses.

Importantly, on each data-reduced symptom axis, some subset of PSD patients received a score near zero. This does not imply that these patients were unimpaired; rather, the symptom configurations for these patients were orthogonal to variation along this specific axis. While in the current paper we demonstrate individual selection based on symptom scores along a single axis, it may also be possible to use a combination of several symptom dimensions to further inform individual treatment selection. For example, for patients with above-threshold symptom scores along several dimensions, a combination of these *PCs* may be more predictive of their neural features and in turn be a better indication of treatment selection for these individuals. Alternatively, the symptom dimensions can be used as exclusionary criteria for selectively picking out individuals along only one particular axis (e.g. filtering out patients above-threshold on *PCs 1, 3, 4,* and *5* to selectively study patients with only severe *PC2* cognitive deficits). Additionally, the ICA solution revealed oblique axes which are linear combinations of the *PCs*, e.g. *IC2* which is oblique to *PCs 1, 2,* and *3* (**Appendix 1 - Fig. 4**). However, these *ICs* do not result in as robust or unique a neural circuit mapping as do the *PCs*, as shown in **Appendix 1 - Fig. 13**. The observation that PSD are associated with multiple complex symptom dimensions highlights an important intuition that may extend to other mental health spectra. Additionally, the PSD symptom axes reported here are neither definitive nor exhaustive. In fact, close to 50% of all clinical variance was not captured by the symptom PCA - an observation often overlooked in symptom data-reduction studies, which focus on attaining “predictive accuracy”. Such studies rarely consider how much variance remains unexplained in the final data-reduced model and, relatedly, if the proportion of explained variance is reproducible across samples. This is a key property for reliable symptom-to-neural mapping. Thus, we tested if this reproducible low-dimensional PSD symptom space robustly mapped onto neural circuit patterns.

### Leveraging a Robust Low-Dimensional Symptom Representation for Mapping Brain-Behavior Relationships

We show that the dimensionality-reduced symptom space improved the mapping onto neural circuit features (i.e. GBC), as compared to *a priori* item-level clinical symptom measures (**Fig. 3**). This symptom-neural mapping was highly reproducible across various cross-validation procedures and split-half replication (**Fig. 4**). The observed statistical symptom-neural improvement after dimensionality reduction suggests that data-driven clinical variation more robustly covaried with neural features. As noted, the low-rank symptom axes generalized across DSM diagnoses. Consequently, the mapping onto neural features (i.e. GBC) may have been restricted if only a single DSM category or clinical item was used. Importantly, as noted, traditional clinical scales are uni-directional (i.e. zero is asymptomatic, hence there is an explicit floor). Here, we show that data-driven symptom axes (e.g. *PC3*) were associated with bi-directional variation (i.e. no explicit floor effect). Put differently, patients who score highly on either end of these data-driven axes are severely symptomatic but in very different ways. If these axes reflect clinically meaningful phenomena at both tails, then they should more robustly map to neural feature variation, which is in line with reported effects. Therefore, by definition, the derived map for each of the *PCs* will reflect the neural circuitry that may be modulated by the behaviors that vary along that PC (but not others). For example, we named the *PC3* axis *“Psychosis Configuration”* because of its strong loadings onto conventional “positive” and “negative” PSD symptoms. This *PC3 “Psychosis Configuration”* showed strong positive variation along neural regions that map onto the canonical default mode network (DMN), which has frequently been implicated in PSD (***Fryer et al., 2013***; ***Anticevic et al., 2012a***; ***Öngür et al., 2010***; ***Woodward et al., 2011***; ***Baker et al., 2014***; ***Meda et al., 2014***) suggesting that individuals with severe “positive” symptoms exhibit broadly elevated connectivity with regions of the DMN. On the contrary, this bi-directional *“Psychosis Configuration”* axis also showed strong negative variation along neural regions that map onto the sensory-motor and associative control regions, also strongly implicated in PSD (***Ji et al., 2019b***; ***Anticevic et al., 2013b***). The “bi-directionality” property of the *PC* symptom-neural maps may thus be desirable for identifying neural features that support individual patient selection. For instance, it may be possible that *PC3* reflects residual untreated psychosis symptoms in this chronic PSD sample, which may reveal key treatment neural targets. In support of this circuit being symptom-relevant, it is notable that we observed a mild association between GBC and *PC* scores in the CON sample (**Appendix 1 - Fig. 11**). The symptom-neural mapping of dimensionality-reduced symptom scores also produced notable neural maps along other *PC* axes (**Appendix 1 - Fig. S10**). Higher *PC1* scores, indicating higher general functioning, may be associated with lower global connectivity in sensory/cingulate cortices and thalamus, but higher global connectivity with the temporal lobe. Higher *PC2* scores, indicating better cognitive functioning were positively associated with higher medial prefrontal and cerebellar GBC, and negatively with visual cortices and striatum. Importantly, the unique neural circuit variance shown in the *β*_*PC*2_*GBC* map suggest that there are PSD patients with more (or less) severe cognitive deficits independent of any other symptom axis, which would be in line with the observation that these symptoms are not treatable with antipsychotic medication (and therefore should not correlate with symptoms that are treatable by such medications; i.e. *PC3*). Of note, the statistics in the *β*_*P*2_*GBC* map were relatively moderate, opening up the possibility that a more targeted measure of neural connectivity (e.g. seed FC, see below) would reveal a robust circuit map for this particular symptom dimension. Given the key role of cognitive deficits in PSD (***Barch et al., 2013***), this will be a particularly important direction to pursue, especially in prodromal or early-stage individuals where cognitive dysfunction has been shown to predict illness trajectories (***Fusar-Poli et al., 2012***; ***Bora et al., 2014***; ***Bora, 2015***; ***Antshel et al., 2017***). The *PC4 - Affective Valence* axis was positively associated with higher GBC in cingulate and sensori-motor cortices and subcortically with the thalamus, basal ganglia, and anterior cerebellum, which may be implicated in the deficits in social functioning and affective aspects of this symptom dimension. Lastly, positive scores on *PC5,* which we named *“Agitation/Excitement”,* were associated with broad elevated global connectivity in the frontal associative and sensori-motor cortices, thalamus, and basal ganglia. Interestingly, however, the nucleus accumbens (as well as hippocampus, amygdala, cerebellum, and temporal lobe) appear to be negatively associated with *PC5* score (i.e. more severe grandiosity, unusual thought, delusional and disorganized thinking), illustrating another example of a bi-directional axis.

### Deriving Individually Actionable Brain-Behavior Mapping Across the Psychosis Spectrum

Deriving a neuro-behavioral mapping that is resolvable and stable at the individual patient level is a necessary benchmark for deploying symptom-neural “biomarkers” in a clinically useful way. Therefore, there is increasing attention placed on the importance of achieving reproducibility in the psychiatric neuroimaging literature (***Noble et al., 2019***; ***Balsters et al., 2016***; ***Cao et al., 2019***; ***Woo et al., 2017***), which becomes especially important for individualized symptom-neural mapping. Recently, several efforts have deployed multivariate methods to quantify symptom-neural relationships (***Drysdale et al., 2017***; ***Xia et al., 2018***; ***Smith et al., 2015***; ***Moser et al., 2018***; ***Rodrigue et al., 2018***; ***Yu et al., 2019***), highlighting how multivariate techniques may perhaps provide clinically innovative insights. However, such methods face the risk of overfitting for high-dimensional but underpowered datasets (***Dinga et al., 2019***), as recently shown via formal generative modeling (***Helmer et al., 2020***).

Here we attempted to use multivariate solutions (i.e. CCA) to quantify symptom and neural feature co-variation. In principle, CCA is well-suited to address the brain-behavioral mapping problem. However, symptom-neural mapping using CCA in our sample was not reproducible even when using a low-dimensional symptom solution and parcellated neural data as a starting point. Therefore, while CCA (and related multivariate methods such as partial least squares) are theoretically appropriate and may be helped by regularization methods such as sparse CCA, in practice many available psychiatric neuroimaging datasets may not provide sufficient power to resolve stable multivariate symptom-neural solutions (***Helmer et al., 2020***). A key pressing need for forthcoming studies will be to use multivariate power calculators to inform sample sizes needed for resolving stable symptom-neural geometries at the single subject level. Of note, though we were unable to derive a stable CCA in the present sample, this does not imply that the multivariate neuro-behavioral effect may not be reproducible with larger effect sizes and/or sample sizes. Critically, this does highlight the importance of power calculations prior to computing multivariate brain-behavioral solutions (***Helmer et al., 2020***).

Consequently, we tested if a low-dimensional symptom solution can be used in a univariate symptom-neural model to optimize individually predictive features. Indeed, we found that a univariate brain-behavioral space (BBS) relationship can result in neural features that are stable for individualized prediction. Critically, we found that if a patient exhibited a high PC symptom score, they were more likely to exhibit a topography of neural *Δ*GBC that was topographically similar to the *β_PC_GBC* map. This suggests that optimizing such symptom-neural mapping solutions (and ultimately extending them to multivariate frameworks) can inform cross-diagnostic patient segmentation with respect to symptom-relevant neural features. Importantly, this could directly inform patient identification based on neural targets that are of direct symptom relevance for clinical trial design.

### Utilizing Independent Molecular Neuroimaging Maps to “Benchmark” Symptom-Relevant Neural Features

Selecting single patients via stable symptom-neural mapping of BBS solutions is necessary for individual patient segmentation, which may ultimately inform treatment indication. However, it is also critical to be able to relate the derived symptom-neural maps to a given mechanism. Here we highlight two ways to “benchmark” the derived symptom-neural maps by calculating their similarity against independent pharmacological neuroimaging and gene expression maps. We show a proof-of-principle framework for quantifying derived symptom-neural reference maps with two PSD-relevant neuropharmacological manipulations derived in healthy adults via LSD and ketamine. These analyses revealed that selecting single patients, via the derived symptom-neural mapping solution, can yield above-chance quantitative correspondence to a given molecular target map. These data highlight an important effect: it is possible to construct a “strong inference” (***Platt, 1964***) evaluation of single patients’ differential similarity to one molecular target map versus another. For instance, this approach could be applied to maps associated with already approved PSD treatments (such as clozapine, olanzapine, or chlorpromazine (***Lally and MacCabe, 2015***; ***Miyamoto et al., 2005***)) to identify patients with symptom-neural configurations that best capture available treatment-covarying neural targets.

Relatedly, AHBA gene expression maps (***Burt et al., 2018***) may provide an *a priori* benchmark for treatment targets that may be associated with a given receptor profile. Here we show that identified BBS maps exhibit spatial correspondence with neural gene expression maps implicated in PSD - namely serotonin, GABA and interneuron gene expression. This gene-to-BBS mapping could be then used to select those patients that exhibit high correspondence to a given gene expression target.

Collectively, this framework could inform empirically testable treatment selection methods (e.g. a patient may benefit from ketamine, but not serotonin agonists such as LSD/psilocybin). In turn, this independent molecular benchmarking framework could be extended to other approaches (e.g. positron emission tomography (PET) maps reflecting specific neural receptor density patterns (***Farde et al., 1986***; ***Arakawa et al., 2008***)) and iteratively optimized for quantitative patient-specific selection against actionable molecular targets.

### Considerations for Generalizing Solutions Across Time, Severity and Mental Health Spectra

There are several constraints of the current result that require future optimization - namely the ability to generalize across time (early course vs. chronic patients), across a range of symptom severity (e.g. severe psychotic episode or persistent low-severity psychosis) and across distinct symptom spectra (e.g. mood). This applies to both the low-rank symptom solution and the resulting symptom-neural mapping. It is possible that the derived lower-dimensional symptom solution, and consequently the symptom-neural mapping solution, exhibits either time-dependent (i.e. state) or severity-dependent (i.e. trait) re-configuration. Relatedly, medication dose, type, and timing may also impact the solution. Another important aspect that will require further characterization is the possibility of oblique axes in the symptom-neural geometry. While orthogonal axes derived via PCA were appropriate here and similar to the ICA-derived axes in this solution, it is possible that oblique dimensions more clearly reflect the geometry of other psychiatric spectra and/or other stages in disease progression. For example, oblique components may better capture dimensions of neuro-behavioral variation in a sample of prodromal individuals, as these patients are exhibiting early-stage psychosis-like symptoms and may show signs of diverging along different trajectories.

Critically, these factors should constitute key extensions of an iteratively more robust model for individualized symptom-neural mapping across the PSD and other psychiatric spectra. Relatedly, it will be important to identify the ‘limits’ of a given BBS solution - namely a PSD-derived effect may not generalize into the mood spectrum (i.e. both the symptom space and the resulting symptom-neural mapping is orthogonal). It will be important to evaluate if this framework can be used to initialize symptom-neural mapping across other mental health symptom spectra, such as mood/anxiety disorders.

These types of questions will require longitudinal and clinically diverse studies that start prior to the onset of full-blown symptoms (e.g. the North American Prodrome Longitudinal Study (NAPLS) (***Addington et al., 2012***; ***Seidman et al., 2010***)). A corollary of this point is that ~50% of unexplained symptom variance in the current PCA solution necessitates larger samples with adequate power to map this subtle, but perhaps clinically essential, PSD variation.

Notably, the current cohort was adequately powered for symptom data reduction that drove univariate neural mapping. However, this sample was insufficiently powered for resolving stable multivariate symptom-neural relationships even with dimensionality-reduced symptom features. Consequently, the limited sample size necessitated choices for dimensionality reduction of the neural feature space in this study even for univariate analyses. While both parcellation and GBC constitute principled choices, symptom-relevant neural information may have been lost (which may be embedded in a higher-dimensional space). One obvious solution is to increase sample sizes (e.g. via datasets such as the UK Biobank (***Sudlow et al., 2015***)). However, in parallel, it will be critical to develop neurobiologically-informed feature space reduction and/or to optimize the stability of multivariate solutions via regularization methods. Another improvement would be to optimize neural data reduction sensitivity for specific symptom variation (***Shehzad et al., 2014***). We chose to use GBC for our initial geometry characterizations as it is a principled and assumption-free data-reduction metric that captures (dys)connectivity across the whole brain and generates neural maps (where each region can be represented by a value, in contrast to full functional connectivity matrices) that are necessary for benchmarking against molecular imaging maps. However, GBC is a summary measure that by definition does not provide information regarding connectivity between specific pairs of neural regions, which may prove to be highly symptom-relevant and informative. Thus symptom-neural relationships should be further explored with higher-resolution metrics, such as restricted GBC (***Anticevic et al., 2013a***) which can summarize connectivity information for a specific network or subset of neural regions, or seed-based FC using regions implicated in PSD (e.g. thalamus (***Anticevic et al., 2013b***; ***Woodward et al., 2011***)).

Here we focused on the neural blood oxygen level dependent (BOLD) signal from fMRI. However, other modalities such as diffusion-weighted imaging (DWI), PET imaging, or electroencephalography (EEG) could be leveraged. Additional clinically-relevant information could be derived from genetics (such as polygenic risk scores (***Hyman, 2011***; ***O’Donovan et al., 2009***; ***Cao et al., 2020***)) or ecological momentary assessment (EMA) (***Corcoran et al., 2018***; ***Niendam et al., 2018***), especially to parse state vs. trait biomarker variation.

Lastly, building on the proof-of-concept molecular neuroimaging comparisons, it will be imperative to eventually test such predictions in formal clinical trials. An actionable next step would be to optimize patient selection against existing treatments, which could result in higher success rates for drug development trials and potentially have massive impact for developing new interventions. Critically, the opportunity to develop, validate, and refine an individual-level quantitative framework could deliver a more rapid and cost-effective way of pairing patients with the right treatments.

## Conclusions

We show that complex and highly heterogeneous PSD symptom variation can be robustly reduced into a low-rank symptom solution that is cross-diagnostic, individually predictive, generalizable and incorporates cognitive deficits. In turn, the derived PSD symptom axes robustly mapped onto distinct yet reproducible neural patterns, which were predictive at the single-patient level. Leveraging these stable results, we show a proof-of-principle framework for relating the derived symptom-relevant neural maps, at the individual patient level, with molecular targets implicated in PSD via LSD and ketamine neuro-pharmacological manipulations. Lastly, we used AHBA gene expression maps to show that identified PSD symptom-relevant neural maps co-vary with serotonin, GABA and interneuron neural gene expression patterns. This set of symptom-neural mapping results can be iteratively and quantitatively optimized for personalized treatment segmentation endpoints.

## Acknowledgements

We thank J.H. Krystal and I. Levy for useful discussions. This work was supported by the Heffter Research Institute (Grant No. 1-190420), Swiss Neuromatrix Foundation (Grant No. 2016-0111), Swiss National Science Foundation under the framework of Neuron Cofund (Grant No. 01EW1908), Slovenian Research Agency (ARRS) (Grant Nos. J7-8275, J7-6829, P3-0338 to GR), 1U01MH121766 (to AA), R01MH112746 (to JDM), SFARI Pilot Award (to JDM & AA), 5R01MH112189 (to AA), 5R01MH108590 (to AA), and by BlackThorn Therapeutics (J.D.M. & A.A.). Data were provided by the Bipolar-Schizophrenia Network on Intermediate Phenotypes (BSNIP) consortium via the National Institute of Mental Health Data Archive (NDA).

