## Appendix for "Mapping Brain-Behavior Space Relationships Along the Psychosis Spectrum"

#### Appendix 1

##### Note 1 - ICA of Behavioral Symptom Data

We performed an independent component analysis (ICA) on the symptom data as a comparison to our PCA data-reduction solution. Specifically, an ICA was conducted on the behavioral data as an alternative dimensionality reduction method to the PCA. Five components were selected to allow for a direct comparison to be made to the results of the PCA (**Appendix 1 - Fig. 4**). ICA decomposition was computed on the *436 x 36* (patients x symptom variables) data matrix using Hyvarinen's (***Hyvarinen, 1999***) FastICA algorithm, executed via the "icafast" package in R (***Helwig and Hong, 2013***). Data were mean-centered before decomposition. After decomposition, the source signal estimate ("ICA score") for all subjects for each independent component (IC) was regressed on to the neural data across patients, as with *a priori* and PCA scores, described in the methods section. The proportion of variance explained by each independent component (IC) is shown in **Appendix 1 - Fig. 4A**. These percentages are comparable to those observed across all the PCs in **Fig. 1C**. **Appendix 1 - Fig. 4B** shows the correlations between subject scores across all five ICs and five PCs (N=436). Notably, certain pairs of components between the two solutions appear to be highly similar and exclusively mapped to each other (*IC5* and *PC4*; *IC4* and *PC5*). The correspondence between PCs 1-3 and ICs 1-3 appears to be more distributed as opposed to following a one-to-one mapping, likely because PCs are orthogonal to each other whereas ICs are not necessarily orthogonal. For example, *PC3* appears to correspond to *IC2*, but is negatively related to *IC3*, suggesting that these two ICs are oblique to the PC and perhaps share behavioral symptom variation in the ICA solution that is explained by a single *PC3*. Of note, *PC3* (the "Psychosis Configuration" axis) is a bi-directional axis whereby individuals who score either highly positively or highly negatively are highly symptomatic in different ways. The diverging nature of this axis may be not be captured well in the ICA solution, when orthogonality is not enforced. **Appendix 1 - Fig. 13F** shows the correlations across parcels (N=718 parcels) between the five IC maps (**Appendix 1 - Fig. 13A-E**) and five PC maps. Again, the neural pattern resembles that in **Appendix 1 - Fig. 13**, with certain pairs corresponding highly between the the ICA and PCA solutions. The coefficient maps for each of the five ICs are shown in **Appendix 1 - Fig. 13**. To compare the ICA solution to the PCA solution, we performed a correlation between the individual PCA and ICA scores across subjects, as well as between parcel coefficients of the 5 PCA maps and 5 ICA maps across parcels.

##### Note 2 - Establishing the Reproducibility of the Data-reduced Symptom-Neural Mapping Solution

Here we expand on the specific details regarding the cross-validation and reproducibility of the symptom-neural mapping result described in **Fig. 4**. Specifically, **Fig. 4D** shows a summary of a 5-fold bootstrapping symptom-neural map validation. Here the patient sample was randomly split into 5 folds and each fold was held out once as the regression was performed between symptom scores and parcel-level GBC in the other 4 folds. The resulting GBC coefficient map was then correlated with the full sample map performed in all 436 patients. Notably, the correlations between maps for each of the *k*-fold runs and the full model reached effects between *r*=0.8-0.9. The coefficient map for one such fold is shown in **Fig. 4E** for and the correlation between the coefficients in this map and the coefficients of the full regression model (performed in all 436 subjects) is shown in **Fig. 4F**. **Fig. 4G** shows a similar summary for leave-one-site-out validation for the regression of all five PC dimensions and traditional symptom scales onto neural GBC. Here, patients from one of the sites were left out while a regression was performed in all remaining patients. In turn, this effect was correlated with the coefficient map from the full patient sample. Again, the maps are highly similar for all sites, showing that the relationships between behavior and neural data are not driven by any one site in particular. Here, the coefficient map for one example is shown in **Fig. 4H** for PC3. Specifically, the results performed without Site 3 are shown because this is the site with the greatest number of patients and therefore may be the most likely to impact the final effect. The correlation between the coefficients in the resulting map without Site 3 and the coefficients of the full sample regression model are shown in the scatterplot in **Fig. 4I**. Finally, we performed a stratified split-half replication. Here, the full sample of patients was randomly divided into two halves (H1 & H2) with the proportion of each diagnostic group (BPP, SADP, SZP) preserved within each half. Then, PCA on the symptom data was computed in each half independently. As noted in the main text, this resulted in the "observed" PCA values for H1 and H2. In turn, we used loadings from H1 PCA to compute the "predicted" PC scores for subjects in H2, and vice versa. These "predicted" and "observed" PC scores were then independently regressed against parcellated GBC for subjects a given half. This yielded two maps for each PC: one for the "predicted" PC scores based on H1 data; and one for the "observed" PC scores from H1 data. These two symptom-neural maps were then correlated to evaluate reproducibility of the effects. This process was performed for both H1-to-H2 reproducibility as well as H2-to-H1 reproducibility. We performed 1,000 runs of the entire split-half replication analysis for the PC1-5 maps (**Fig. 4J**. Example data for *PC3* H1 and H2 are shown in **Fig. 4K-M**). These results highlight that the behavioral PC-to-GBC mapping yields a highly reproducible "brain-behavior" effect.

#### Note 3 - Multivariate Canonical Correlation Analysis of Symptom and Neural Features in PSD

CCA solves for transformation matrices $\Psi$ and $\Theta$ such that the correlations between the linearly transformed 'latent' matrices ***U*** and ***V*** are maximal (see **Methods** for computational details). Matrix ***U*** contains the latent "behavioral" subject scores for all CV and is a linear composite of behavioral feature matrix ***B*** scaled by the loadings in $\Psi$; similarly, ***V*** is the latent neural score matrix and is the linear composite of neural features ***N*** transformed by $\Theta$. Each column in ***U*** and ***V*** is known as a canonical variate (CV); each corresponding pair of canonical variates (e.g. ***U1*** and ***V1***) is referred to as a canonical mode. The correlations between the transformed latent matrices ***U*** and ***V*** are maximized (**Fig. 5B**), under the constraint that the canonical modes are orthogonal and independent to each other. Hence the correlation between each subsequent pair of CVs is computed from the residuals of the previous pair, until the number of columns in the smaller of the two matrices is reached.

In the main text we show that our CCA solution is not reproducible. We also examined whether the size of the neural feature space affects the CCA solution. To this end we computed CCA using:i) 180 symmetrized cortical parcels; ii) 359 bilateral subcortex-only parcels; iii) 192 symmetrized subcortical parcels; iv) 12 functional whole-brain networks, from the recently developed cortico-subcortical CAB-NP parcellation (***Glasser et al., 2016***; ***Ji et al., 2019c***). We show results for subcortical (192 neural features) in **Appendix 1 - Fig. 14**, as well as network-level (12 features neural features) in **Appendix 1 - Fig. 15**. We evaluated CCA significance via permutation testing whereby patient order was randomly shuffled 5,000 times (such that the association between behavioral and neural features was violated) and a CCA solution was computed for each shuffle; see **Methods**).

For each CCA solution, we examined how much symptom variance can be accounted for by the neural feature variance. Specifically, the correlation between behavioral features

***B*** and the latent neural matrix ***V*** (i.e. ***N****Θ*) reflects the amount of variance in ***B*** that can be explained by the neural CVs in ***V*** (**Fig. 5E**). In the main text, **Fig. 5F** shows the proportion of behavioral variance explained by each of the neural CVs in a CCA performed between 180 neural features and all 36 symptom measures. **Fig. 5G** shows the result for the 5 PC-derived symptom dimensions. Given that the symptom PCs accounted for ∼50.93% of total symptom variance (see **Fig. 1**), the percentages here were scaled to reflect the total proportion of behavioral variance explained by each CV. The amount of neural variance explained by each behavioral CV is shown in **Appendix 1 - Fig. 17**.

Next, we examined the proportion of symptom variance explained by each neural canonical variate (CV). Because the PCs accounted for ∼50.93% of total symptom variance (**Fig. 1**), the percentages here were scaled accordingly. While the CCA using PC scores has fewer features than the one using item-level symptom measures, each neural CV explained more behavioral variance (**Fig. 5G** versus **5F**), suggesting that the PCs may capture neurally-relevant symptom variation with fewer features. Each CV has a pair of associated symptom and neural profiles, which can inform interpretation. This is shown for *CV3* in **Fig. 5H–K**. Specifically, the latent scores for an example behavioral canonical variate (CV3) across diagnostic groups normalized to controls are shown in **Fig. 5H**. The profile of PC loadings onto *CV3* are shown in **Fig. 5I**. Full results are shown in **Appendix 1 - Fig. 17**. CCA loadings are computed as the correlation of the latent matrix ***U*** (or ***V***) and the input data matrix ***B*** (or ***N***). These CCA loadings reflect the amount of variance in the original data that is extracted by the canonical variates. The high negative loadings of *CV3* onto *PCs 1,4,5* versus the high positive loadings of *CV3* onto *PC3* in **Fig. 5I** indicate that *CV3* captures complex symptom variation. This can also be seen by the projecting 36 symptom measure loadings from a given PC onto *CV3* **Fig. 5J**. Here the most positively loaded items were #7 "Delusions" (PANSS P1), #9 "Hallucinations" (PANSS P3), #21 "Somatic Concern" (PANSS G1) and #29 "Unusual Thought Content" (PANSS G9), while negatively-loaded items include #16 "Poor Rapport" ((PANSS N3), #19 "Lack of Spontaneity" (PANSS N6), and #28 "Uncooperativeness" (PANSS G8). Conversely, the corresponding neural factor loadings for *CV3* are shown in **Fig. 5K**, illustrating GBC patterns associated with the *CV3* profile. Data for all 5 CVs are shown in **Appendix 1 - Fig. 16**.

Interestingly, the cross-validation of the CCA effect was reproducible when using overlapping cross-validation strategies **Appendix 1 - Fig. 18**. However, as noted in the main text, the CCA solution was not reproducible with out-sample cross-validation strategies even though the effect passed p<0.05 permutation testing. This highlights the importance of rigorously testing for out-of-sample reproducibility of brain-behavioral effects across independent sub-samples (***Helmer et al., 2020***).

#### Note 4 - Establishing the Reproducibility of the CCA Solution

Here we outline the details of the split-half replication for the CCA solution. Specifically, the full patient sample was randomly split (referred to as "H1" and "H2" respectively), while preserving the proportion of patients in each diagnostic group. Then, CCA was performed independently for H1 and H2. While the loadings for behavioral PCs and original symptom measures are somewhat similar (mean $r \approx 0.5$) between the two CCAs in each run, the neural loadings were not stable across H1 and H2 CCA solutions. Critically, CCA results did not perform well for leave-one-subject-out cross-validation (**Fig. 5M**). Here, one patient was held out while CCA was performed using all data from the remaining 435 patients. The loadings matrices $\Psi$ and $\Theta$ from the CCA were then used to calculate the "predicted" neural and behavioral latent scores for all 5 CVs for the patient that was held out of the CCA solution. This process was repeated for every patient and the final result was evaluated for reproducibility. As described in the main text, this did not yield reproducible CCA effects (**Fig. 5M**).

Of note, CCA may yield higher reproducibility if the neural feature space were to be further reduced. As noted, our approach was to first parcellate the BOLD signal and then use GBC as a data-driven method to yield a neuro-biologically and quantitatively interpretable neural data reduction, and we additionally symmetrized the result across hemispheres. Nevertheless, in sharp contrast to the PCA univariate feature selection approach, the CCA solutions were still not stable in the present sample size of $N = 436$. Indeed, a multivariate power analysis (***Helmer et al., 2020***) estimates that the following sample sizes will be required to sufficiently power a CCA between 180 neural features and 5 symptom features, at different levels of true canonical correlation ($r_t rue$):

- if $r_{true} = 0.1$: $n_{required} = 103,545$
- if $r_{true} = 0.3$: $n_{required} = 8,145$
- if $r_{true} = 0.5$: $n_{required} = 2,497$

To test if further neural feature space reduction may be improve reproducibility, we also evaluated CCA solutions with neural GBC parcellated according to 12 brain-wide functional networks derived from the recent HCP-driven network parcellation (***Ji et al., 2019c***). Again, we computed the CCA for all 36 symptom measures as well as 5 *PCs* (**Appendix 1 - Fig. 15**). As with the parcel-level effects, the network-level CCA analysis produced significant results (for *CV1* when using 36 symptom measure scores and for all 5 CVs when using the 5 PC-derived scores). Here the result produced much lower canonical correlations (~0.3-0.5; however, these effects (for *CV1*) clearly exceeded the 95% confidence interval generated via random permutations, suggesting that they may reflect the true canonical correlation. We observed a similar result when we evaluated CCAs computed with neural GBC from 192 symmetrized subcortical parcels and 36 symptoms or 5 PCs (**Appendix 1 - Fig. 14**). In other words, data-reducing the neural signal to 12 functional networks likely averaged out parcel-level information that may carry symptom-relevant variance, but may be closer to capturing the true effect. Indeed, the power analysis suggests that the current sample size is closer to that needed to detect an effect with 12 vs. 5 features:

- if $r_{true} = 0.1$: $n_{required} = 8,855$
- if $r_{true} = 0.3$: $n_{required} = 696$
- if $r_{true} = 0.5$: $n_{required} = 213$

Note that we do not present a CCA conducted with parcels across the whole brain, as the number of variables would exceed the number of observations. However, the multivariate power analysis using 718 neural features and 5 symptom features estimates that the following sample sizes would be required to detect the following effects:

- if $r_{true} = 0.1$: $n_{required} = 421,608$
- if $r_{true} = 0.3$: $n_{required} = 33,166$
- if $r_{true} = 0.5$: $n_{required} = 10,168$

This analysis suggests that even the lowest bound of ~10k samples exceeds the present available sample size by two orders of magnitude.

#### Note 5 - Evaluating Individual Patients in Discovery and Replication Samples via Similarity to Target Brain-Behavior Spaces

We tested a hybrid neuro-behavioral patient selection strategy for single *PC* symptom axes by first imposing a PC-based symptom threshold, followed by a target neural similarity threshold driven by the most highly predictive symptom-neural map features. Specifically, the “neural similarity prediction index (NSPI)" computes a patient-specific Spearman’s $\rho$ between that patient’s $\Delta GBC^{obs}$ and the group reference $\beta_{PC3}GBC^{obs}$ map using the maximally predictive $P = 39$ parcels (see **Appendix 1 - Fig. 28** for whole-brain results and alternative similarity metrics). **Appendix 1 - Fig. 27A** shows a significant relationship between each patient’s *PC3* symptom score (X-axis) and the neural similarity index (Y-axis). In turn, **Appendix 1 - Fig. 27B** shows binned results, which provides a visual intuition for patient segmentation across both the neural and behavioral indices (**Appendix 1 - Fig. 27A**, right: binned by $\rho = 0.1$ & $PC3_{score} = 0.5$). For patients at either tail, the neuro-behavioral relationship was robust. Conversely, patients with a low absolute *PC3* score showed a weak relationship with symptom-relevant neural features. This is intuitive because these individuals do not vary along the selected PC symptom axis.

**Appendix 1 - Fig. 27C** shows the mean NSPI across subjects within each PC symptom bin along the X-axis. The resulting sigmoid captures that patients exhibit greater neural similarity if their PC symptom scores are more extreme. To evaluate if this relationship can yield a personalized patient selection, we computed the absolute NSPI **Appendix 1 - Fig. 27D**). The effect was approximated by a quadratic function, highlighting that patients with extreme *PC3* scores (either positive or negative) exhibited a stronger NSPI (i.e. personalized neural effects that strongly resembled the reference $(\beta_{PC3}GBC^{obs})^P = 39$ map). **Appendix 1 - Fig. 27E** shows the application of this neuro-behavioral selection procedure, demonstrating that PSD patients with extreme *PC3* scores (defined at the top/bottom 10th percentile of the “discovery” sample, $+2.17 < PC3_{score} < -2.41$) exhibit high NSPI values. We observed an inherent trade-off such that if the PC score threshold was raised then neural target similarity confidence goes up, but fewer patients will be selected. In the discovery PSD sample, all patients were accurately selected above the following neuro-behavioral thresholds: $90^{th}\%tile < PC3_{score} < 10^{th}\%tile$ and $|\rho| > 0.4$ (**Appendix 1 - Fig. 27E**, 34/436 patients selected, green line). We show consistent effects for when the selection was applied to *PC5* (**Appendix 1 - Fig. 27F**; results for all PCs in **Appendix 1 - Fig. 29**).

To test if the neuro-behavioral selection is generalizable, we used an independent cross-diagnostic sample of 30 patients diagnosed with SZP and 39 diagnosed with obsessive-compulsive disorder (OCD) (**Appendix 1 - Fig. 27G**, see **Methods** and **Appendix 1 - Table 4** for details). Applying the “discovery” selection thresholds yielded similar results for ∼6% of the cross-diagnostic “replication” sample for *PC3* (**Appendix 1 - Fig. 27G**, full analyses in **Appendix 1 - Fig. 28C**). Notably, no replication sample patients were selected along the neuro-behavioral thresholds for *PC5* (**Appendix 1 - Fig. 27H**). While there are SZP patients in the replication cross-diagnostic sample, few scored highly on *PC5* and none met the neural similarity threshold, emphasizing that not all patients within the same DSM-based diagnosis will exhibit variation along the same neuro-behavioral axis. Collectively, these results show that data-driven symptom scores can pinpoint individual patients for whom their neural variation strongly maps onto a target neural reference map. These data also highlight that both symptom and neural information for an independent patient can be quantified in the reference ‘discovery’ BBS using their symptom data alone.

**Appendix 1 - Table 1 | Glossary of Terms and Abbreviations**

| Abbreviation / Term | Definition |
|---|---|
| | **General & Behavioral Terms** |
| PSD | Psychosis spectrum disorder; or patients diagnosed with a PSD. |
| BPP | Bipolar disorder with psychosis; or patients diagnosed with BPP. |
| SADP | Schizoaffective disorder with psychosis; or patients diagnosed with SADP. |
| SZP | Schizophrenia with psychosis; or patients diagnosed with SZP. |
| CON | Control subjects. |
| BACS | Brief Assessment of Cognition in Schizophrenia. |
| PANSS | Positive and Negative Syndrome Scale. |
| Symptom measures | The 36 behavioral items from the BACS and PANSS. |
| PCA, PC | Principal component analysis, and principal component. |
| ICA, IC | Independent component analysis, and independent component. |
| | **Neural Terms** |
| FC | Functional connectivity (Fisher's r-to-Z transformed Pearson correlation values). |
| GBC | Global brain connectivity; computed for each brain location as the the mean FC across the whole brain. |
| $\beta_{PC}GBC$ | The beta coefficient map from a mass univariate regression of PC scores on to GBC across subjects. |
| $\beta_{PC}GBC^{obs}$ | The beta coefficient map from a mass univariate regression of observed (measured) symptom PC scores on to observed (measured) GBC across subjects. |
| $\beta_{PC}GBC^{pred}$ | The beta coefficient map from a mass univariate regression of predicted symptom PC scores on to observed (measured) GBC across subjects. |
| GBC-PCA | A PCA performed on the neural GBC data across all N=638 PSD and CON subjects. |
| $G\hat{B}C_{woPC1}$ | The GBC matrix reconstructed without the first PC of the GBC-PCA. |
| $G\hat{B}C_{woPC1-2}$ | The GBC matrix reconstructed without the first PC of the GBC-PCA. |
| $G\hat{B}C_{woPC1-3}$ | The GBC matrix reconstructed without the first two PCs of the GBC-PCA. |
| $G\hat{B}C_{woPC1-4}$ | The GBC matrix reconstructed without the first three PCs of the GBC-PCA. |
| | **Multivariate Analyses Terms** |
| CCA | Canonical correlation analysis. |
| CV | Canonical variate from a CCA solution. |
| 180 vs. 36 CCA | The CCA computed between 180 neural features and 36 symptom measures. |
| 180 vs. 5 CCA | The CCA computed between 180 neural features and 5 symptom PCs. |
| N, B | The input neural and behavioral data matrices in a CCA, respectively. |
| $\hat{N}$, $\hat{B}$ | The predicted neural and behavioral data matrices in the CCA cross-validation, respectively. |
| $\theta$,$\Psi$ | The neural and behavioral transformation matrices in a CCA, respectively. |
| U, V | The transfomed neural and behavioral data matrices in a CCA, respectively. |
| | **Single-Subject Prediction Analyses Terms** |
| $\Delta$GBC | The difference between an individual's measured GBC map and the mean GBC map computed for the whole group; reflects individual differences in GBC, which may otherwise be conflated by the group mean. |
| $\beta_{PC}GBC^{obs}{}_{N-1}$ | The beta coefficient map computed across N-1 subjects, using observed symptom PC and GBC data. |
| $S^{obs}$ | The vector of observed symptom scores for a single PC (i.e. from a PCA computed using symptom measures), with length N-1 (as subject i is left out). |
| $S^{pred}$ | The predicted symptom PC for subject i, using the loadings from a leave-one-out PCA computed on N-i subjects. |
| dpGBC | Dot product between a subject's $\Delta$GBC map and a $\beta_{PC}GBC$ map. |
| $dpGBC^{obs}{}_{i}$ | Dot product between a subject i's $\Delta$GBC map and the $\beta_{PC}GBC^{obs}{}_{N-1}$ map computed without that subject. |
| $dpGBC^{pred}{}_{i}$ | The predicted dpGBC for a subject i, predicted from a regression model using the $dpGBC^{obs}$ from all other subjects and the predicted $S^{pred}$ for subject i. |
| $P_{select}$ | The number of parcels in the subset of the brain map used in a particular analysis. $P_{select}$=718 for the full whole-brain map. |
| BBS | Brain-Behavior Space; the symptom and neural mapping for a particular dimension. |

**Appendix 1 Table 1. Glossary of the key terms and abbreviations used in the study.**

**Appendix 1 - Table 2 | BSNIP Dataset Clinical and Demographic Characteristics**

| Characteristic | HCS (N =202) | | PSD (N=436) | | BPP (N=150) | | SADP (N=119) | | SZP (N=167) | |
|---|---|---|---|---|---|---|---|---|---|---|
| | Mean | S.D. | Mean | S.D | Mean | S.D. | Mean | S.D. | Mean | S.D. |
| **Age (years)** | 37.18 | 12.18 | 35.33 | 12.31 | 36.29 | 12.94 | 35.86 | 11.87 | 34.16 | 11.99 |
| **Gender (% male)** | 42.08 | - | 48.62 | - | 31.33 | - | 44.54 | - | 67.07 | - |
| **Parental Education (years)** | 13.35 | 3.42 | 13.67 | 3.43 | 14.16 | 3.44 | 13.05 | 3.95 | 13.62 | 2.93 |
| **Participant's Education** | 14.76 | 2.27 | 13.47 | 2.34 | 14.22 | 2.31 | 13.25 | 2.22 | 12.93 | 2.28 |
| **Handedness (% right)** | 85.64 | - | 85.78 | - | 83.33 | - | 88.24 | - | 86.23 | - |
| **Signal-to-noise (SNR)** | 231.03 | 82.70 | 218.34 | 93.44 | 229.78 | 100.09 | 230.93 | 93.83 | 198.10 | 83.13 |
| **% Frames Flagged** | 2.81 | 3.99 | 6.17 | 9.09 | 6.41 | 9.67 | 5.32 | 9.32 | 6.57 | 8.35 |
| **Medication (CPZ equivalents)** | - | - | 443.20 | 402.92 | 318.61 | 321.06 | 514.43 | 462.70 | 490.64 | 395.76 |
| **PANSS Positive Symptoms** | 7.03 | 0.30 | 15.66 | 5.35 | 12.87 | 4.35 | 18.19 | 5.20 | 16.37 | 5.15 |
| **PANSS Negative Symptoms** | 7.01 | 0.16 | 14.59 | 5.10 | 12.01 | 3.66 | 15.55 | 4.52 | 16.22 | 5.68 |
| **PANSS General Psychopathology** | 16.04 | 0.36 | 31.45 | 8.67 | 28.72 | 8.20 | 34.57 | 8.70 | 31.67 | 8.30 |
| **PANSS Total Psychopathology** | 30.08 | 0.61 | 61.67 | 16.32 | 53.59 | 13.69 | 68.22 | 15.72 | 64.26 | 16.04 |
| **BACS Cognitive Score (Z)** | 0.02 | 1.10 | -1.22 | 1.20 | -0.83 | 1.19 | -1.30 | 1.10 | -1.52 | 1.19 |

**Appendix 1 Table 2. Clinical and demographic details for the BSNIP sample.** Clinical and demographic measures are shown at the group level for the BSNIP sample used for initial "discovery" analyses. Means and standard deviations were computed for each of the diagnostic groups. Abbreviations: CON, controls; PSD, psychosis spectrum disorder patients; BPP, bipolar disorder with psychosis; SADP, schizoaffective disorder with psychosis; SZP, schizophrenia; CPZ, chlorpromazine; PANSS, Positive and Negative Syndrome Scale for Schizophrenia, BACS, Brief Assessment of Cognition in Schizophrenia.

Appendix 1 - Table 3 | BSNIP Dataset Scan Acquisition Parameters

| BOLD | TR (ms) | TE (ms) | FA (deg.) | Slices (N) | Slice Order | Matrix (mm) | Voxel (mm) | Vendor |
|---|---|---|---|---|---|---|---|---|
| Baltimore | 2210 | 30 | 70 | 36 | I-A | 64 x 64 | 3.4 x 3.4 x 3 | Siemens Trio |
| Hartford | 1500 | 27 | 70 | 29 | S-A | 64 x 64 | 3.4 x 3.4 x 5 | Siemens Allegra |
| Detroit | 1570 /1720 | 22 | 60 | 29 | S-A | 64 x 64 | 3.4 x 3.4 x 4 | Siemens Trio |
| Dallas | 1500 | 27 | 60 | 29 | S-A | 64 x 64 | 3.4 x 3.4 x 4 | Philips |
| Chicago | 1775 | 27 | 60 | 29 | S-A | 64 x 64 | 3.4 x 3.4 x 4 | GE Signa HDX |
| Boston | 3000 | 27 | 60 | 30 | S-A | 64 x 64 | 3.4 x 3.4 x 5 | GE Signa HDX |
| **T1w** | **TR (ms)** | **TE (ms)** | **FA (deg.)** | **Slices (N)** | **Slice Order** | **Matrix (mm)** | **Voxel (mm)** | **Vendor** |
| Baltimore | 2300 | 2.91 | 9 | 160 | — | 256 x 240 | 1 x 1 x 1.2 | Siemens Trio |
| Hartford | 2300 | 2.91 | 9 | 160 | — | 256 x 240 | 1 x 1 x 1.2 | Siemens Allegra |
| Detroit | 2300 | 2.94 | 9 | 160 | — | 256 x 240 | 1 x 1 x 1.2 | Siemens Trio |
| Dallas | 6.6 | 2.8 | 8 | 170 | — | 256 x 256 | 1 x 1 x 1.2 | Philips |
| Chicago | 6.98 | 2.84 | 8 | 166 | — | 256 x 256 | 1 x 1 x 1.2 | GE Signa HDX |
| Boston | 6.98 | 2.84 | 8 | 166 | — | 256 x 256 | 1 x 1 x 1.2 | GE Signa HDX |

**Appendix 1 Table 3. Scan acquisition parameters for the BSNIP dataset across sites.** Scan acquisition details are shown for the 6 BSNIP acquisition sites. Adapted from (***Meda et al., 2015***).

**Appendix 1 - Table 4 | Replication Cross-diagnostic Dataset Clinical and Demographic Characteristics**

| Characteristic | CON (N=34) | | PTT (N=69) | | OCD (N=39) | | SCZP (N=30) | |
|---|---|---|---|---|---|---|---|---|
| | Mean | S.D. | Mean | S.D | Mean | S.D. | Mean | S.D. |
| **Age (years)** | 29.91 | 8.11 | 33.7 | 12.89 | 32.23 | 12.32 | 35.63 | 13.63 |
| **Gender (% male)** | 61.7 | - | 59.4 | - | 46.1 | - | 76.7 | - |
| **Parental Education (years)** | 15.05 | 3.31 | 13.98 | 3.45 | 14.02 | 3.78 | 13.93 | 3.01 |
| **Participant's Education** | 16.37 | 2.34 | 14.16 | 2.12 | 15.22 | 1.95 | 12.79 | 2.34 |
| **Handedness (% right)** | 93.9 | - | 91.3 | - | 84.6 | - | 100.0 | - |
| **Signal-to-noise (SNR)** | 53.45 | 5.59 | 51.86 | 5.63 | 52.29 | 5.28 | 51.33 | 6.08 |
| **% Frames Flagged** | 0.21 | 0.53 | 7.27 | 15.53 | 5.11 | 13.81 | 10.07 | 17.37 |
| **PANSS Positive Symptoms** | 7.58 | 1.06 | 11.40 | 4.23 | 8.97 | 2.74 | 14.56 | 6.16 |
| **PANSS Negative Symptoms** | 7.75 | 1.57 | 11.06 | 3.59 | 8.80 | 2.21 | 14.00 | 5.39 |
| **PANSS General Psychopathology** | 32.62 | 2.62 | 48.80 | 10.96 | 42.71 | 9.07 | 56.72 | 13.42 |
| **PANSS Total Psychopathology** | 47.95 | 5.25 | 71.26 | 18.78 | 60.48 | 14.02 | 85.28 | 24.97 |
| **BACS Cognitive Score (Z)** | 0.00 | 1.00 | -0.67 | 1.45 | -0.07 | 1.03 | -1.45 | 1.55 |

**Appendix 1 Table 4. Clinical and demographic details for the independent replication cross-diagnostic sample.** Clinical and demographic measures are shown at the group level for the independent clinical sample used for replication analyses presented in main text (**Fig. 7**). Means and standard deviations were computed for each of the diagnostic groups. Abbreviations: CON, controls; PTT, all patients; OCD, obsessive-compulsive disorder; SZP, schizophrenia; PANSS, Positive and Negative Syndrome Scale for Schizophrenia, BACS, Brief Assessment of Cognition in Schizophrenia.

### Appendix 1 - Figures

| Question | Results | Corresponding Figures |
|---|---|---|
| Can data-reduction methods (e.g. principal component analysis, PCA) reliably map symptom axes across PSD that include both canonical symptoms and cognitive deficits? | The PCA on 36 behavioral measures and 436 PSD patients produced five significant components (PCs) spanning key symptom domains and oblique to conventional symptom factors and diagnostic categories. | Main Fig. 1<br>Supplementary Fig. S3-S4 |
| Are the derived dimensionality-reduced symptom axes in PSD stable and replicable? | The behavioral PCA solution was stable across split-half replication, leave-site-out, and k-fold cross-validations, and not driven by medication effects. | Main Fig. 2<br>Supplementary Fig. S5-S7 |
| Do dimensionality-reduced symptom axes yield stronger statistical neural maps across PSD, relative to conventional scales & diagnoses? | PC symptom scores yielded a much stronger statistical effect with neural features (as measured by global brain connectivity, GBC) across patients than did conventional symptom scores. | Main Fig. 3<br>Supplementary Fig. S8-S10 |
| Are the PC brain-behavior maps stable and replicable? | PC brain-behavior maps are stable and reproducible across split-half, leave-site-out and k-fold cross-validations. | Main Fig. 4<br>Supplementary Fig. S11 |
| Can a multivariate approach (e.g. canonical correlation analysis, CCA) be used to derive a stable neuro-behavioral mapping? | CCA produced a stable within-sample solution, but performed poorly on out-of-sample cross-validations. | Main Fig. 5<br>Supplementary Fig. S12-S16 |
| Is the computed symptom-to-neural mapping actionable for individual patient selection? | A majority of neural variance in PSD patients is not symptom-relevant. However, brain-behavioral maps can be optimized to select specific patients along PC axes. This method can also be applied to select patients from independent datasets. | Main Fig. 6 & 7<br>Supplementary Fig. S17-S22 |
| Can we inform molecular mechanism and treatment decisions by relating patient-specific brain-behavioral maps to independently-acquired pharmacological neuroimaging maps? | Pharmacological neuroimaging maps targeting mechanisms of interest (e.g. ketamine and LSD) can be used as independent benchmarks against which patient-specific brain-behavioral maps can be evaluated. | Main Fig. 7<br>Supplementary Fig. S23 |
| Can we inform molecular mechanism and potential novel therapeutic targets by relating brain-behavior maps to gene expression maps? | Gene expression maps for genes implicated in PSD neuropathology (e.g. serotonin and GABA receptors, interneuron markers) can inform mechanism and potential novel targets relating to specific axes of neuro-behavioral variation. | Main Fig. 8<br>Supplementary Fig. S24 |

**Appendix 1 Figure 1. Summary of the key questions, results, and supporting data in this study.**

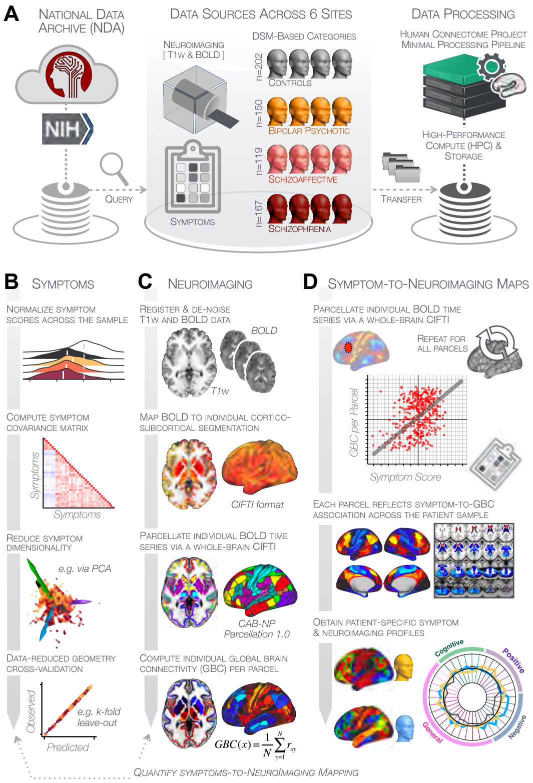


**Appendix 1 Figure 2. Study workflow used study to quantify shared neural and behavioral variation in individuals diagnosed with PSD. (A)** Data from the BSNIP study were acquired from the National Institute of Health Data Archive (NDA). T1-weighted structural and resting-state BOLD neuroimaging data were obtained for a total of 638 individuals (202 controls, 150 patients with a diagnosis of bipolar disorder with psychosis, 119 with a diagnosis of schizoaffective disorder, and 167 patients with schizophrenia). Data were processed through the Human Connectome Project's (HCP) Minimal Preprocessing Pipeline with modifications made for "legacy" BOLD and T1w data, which are now featured as a standard option in the HCP pipelines provided by our team (https://github.com/Washington-University/HCPpipelines/pull/156) using Yale High Performance Computing clusters. **(B)** Symptom data were first normalized across the sample. The correlation matrix across all 436 PSD patients and 36 symptom measures was computed followed by dimensionality reduction (e.g. using PCA or ICA). The dimensionality-reduced solution was then cross-validated to assess stability and reproducibility across sites, *k*-fold cross-validations, leave-subject-out and split-half approaches. **(C)** In parallel, all neuroimaging data were processed as noted above. T1w and resting-state BOLD images were preprocessed using a modified version of the HCP Minimal Preprocessing Pipeline, including individual-subject registration of structural and function data, de-noising, and mapping of BOLD data on a hybrid surface-volume cortico-subcortical format (Connectivity Informatics Technology Initiative [CIFTI] format, see **Methods** for details). After registration to a standard CIFTI template BOLD data were parcellated at the individual subject level using the Cole-Anticevic Brain-wide Network Partition (CAB-NP), which is a functionally-defined network and parcel-level partition in the CIFTI space encompassing both cortex and subcortex (***Glasser et al., 2016***; ***Ji et al., 2019c***). Lastly, a global brain connectivity (GBC) map for each subject was computed by taking the mean functional connectivity of each parcel with all other parcels in the brain at the single subject level. **(D)** After symptom and neural data were fully processed in tandem, the symptom-to-neuroimaging mapping was quantified across subjects. Specifically, the relationship between data-reduced symptom scores and GBC was computed for each parcel across all patients. This produced a group-level symptom-neural map, which was subsequently cross-validated using leave-site-out, *k*-fold, leave subject-out as well as split-half approaches to assess reproducibility of the effect. Finally, the stable symptom-neural mapping result was further feature-optimized for single-subject prediction. This yielded a set of parcels for quantifying patient-specific symptom-neural prediction based on a cross-validated group reference map as well as comparison of the selected parcels with independent molecular neuroimaging maps (i.e. pharmacological (***Anticevic et al., 2012c***; ***Preller et al., 2018***) and gene expression maps (***Burt et al., 2018***)).

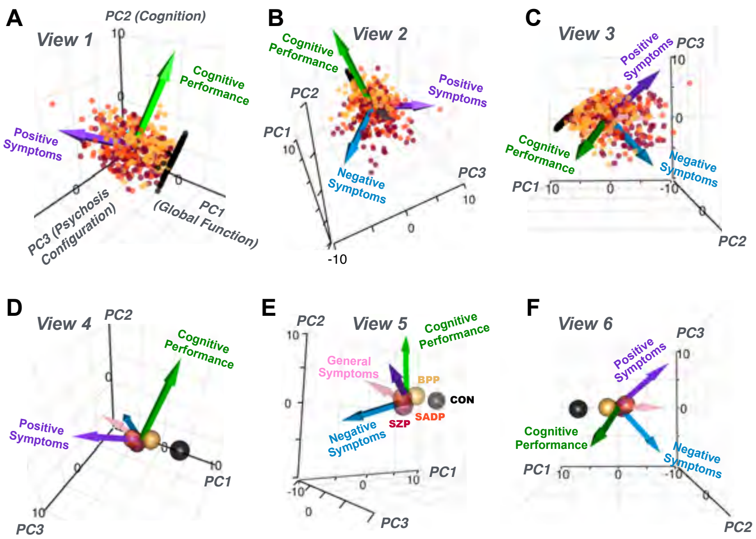


**G** Individual Items & PCA Loadings

| Group | Item | PC1 Global Functioning | PC2 Cognitive Function | PC3 Psychosis Configuration | PC4 Affective Valence | PC5 Agitation/ Excitement |
|---|---|---|---|---|---|---|
| Cognitive | 1 BACS Digital Sequence | 0.089 | 0.320 | -0.152 | 0.084 | -0.192 |
| | 2 BACS Sequence Coding | 0.070 | 0.375 | -0.102 | 0.149 | -0.135 |
| | 3 BACS Token Motor | 0.059 | 0.250 | -0.089 | 0.083 | -0.024 |
| | 4 BACS Tower | 0.062 | 0.258 | -0.161 | 0.074 | -0.187 |
| | 5 BACS Verbal Fluency | 0.085 | 0.320 | -0.053 | 0.087 | -0.202 |
| | 6 BACS Verbal Memory | 0.090 | 0.322 | -0.127 | 0.134 | -0.135 |
| Positive | 7 PANSS P1 - Delusions | -0.221 | -0.003 | 0.129 | -0.053 | -0.402 |
| | 8 PANSS P2 - Conceptual Disorganization | -0.208 | -0.059 | 0.138 | 0.211 | -0.131 |
| | 9 PANSS P3 - Hallucinations | -0.188 | -0.076 | 0.173 | -0.158 | -0.256 |
| | 10 PANSS P4 - Excitement | -0.142 | 0.199 | 0.235 | 0.076 | 0.268 |
| | 11 PANSS P5 - Grandiosity | -0.146 | 0.141 | 0.235 | 0.125 | -0.165 |
| | 12 PANSS P6 - Suspiciousness/Persecution | -0.225 | 0.081 | 0.074 | -0.180 | -0.090 |
| | 13 PANSS P7 - Hostility | -0.160 | 0.235 | 0.125 | 0.060 | 0.222 |
| Negative | 14 PANSS N1 - Blunted Affect | -0.147 | -0.169 | -0.322 | 0.061 | -0.140 |
| | 15 PANSS N2 - Emotional Withdrawal | -0.212 | 0.034 | -0.311 | -0.113 | 0.037 |
| | 16 PANSS N3 - Poor Rapport | -0.171 | -0.033 | -0.264 | 0.250 | 0.077 |
| | 17 PANSS N4 - Passive/Apathetic Social Withdrawal | -0.164 | -0.006 | -0.252 | -0.254 | -0.059 |
| | 18 PANSS N5 - Difficulty in Abstract Thinking | -0.145 | -0.201 | 0.103 | 0.032 | -0.098 |
| | 19 PANSS N6 - Lack of Spontaneity and Flow of Conversation | -0.152 | -0.136 | -0.297 | 0.200 | 0.042 |
| | 20 PANSS N7 - Stereotyped Thinking | -0.236 | -0.002 | 0.000 | 0.174 | -0.039 |
| General | 21 PANSS G1 - Somatic Concern | -0.164 | 0.077 | 0.123 | -0.163 | -0.108 |
| | 22 PANSS G2 - Anxiety | -0.156 | 0.199 | 0.031 | -0.315 | 0.110 |
| | 23 PANSS G3 - Guilt Feelings | -0.105 | 0.163 | -0.022 | -0.214 | -0.016 |
| | 24 PANSS G4 - Tension | -0.175 | 0.175 | 0.017 | -0.103 | 0.241 |
| | 25 PANSS G5 - Mannerisms and Posturing | -0.178 | -0.052 | -0.151 | 0.203 | 0.030 |
| | 26 PANSS G6 - Depression | -0.160 | 0.148 | -0.116 | -0.354 | 0.015 |
| | 27 PANSS G7 - Motor Retardation | -0.164 | -0.125 | -0.313 | 0.042 | 0.023 |
| | 28 PANSS G8 - Uncooperativeness | -0.151 | 0.113 | 0.036 | 0.250 | 0.255 |
| | 29 PANSS G9 - Unusual Thought Content | -0.209 | -0.065 | 0.131 | 0.024 | -0.388 |
| | 30 PANSS G10 - Disorientation | -0.160 | -0.075 | 0.135 | -0.009 | 0.112 |
| | 31 PANSS G11 - Poor Attention | -0.195 | 0.015 | 0.128 | 0.218 | 0.085 |
| | 32 PANSS G12 - Lack of Judgment and Insight | -0.164 | -0.013 | 0.059 | 0.217 | -0.125 |
| | 33 PANSS G13 - Disturbance of Volition | -0.199 | -0.004 | -0.127 | 0.043 | 0.137 |
| | 34 PANSS G14 - Poor Impulse Control | -0.189 | 0.159 | 0.144 | 0.048 | 0.221 |
| | 35 PANSS G15 - Preoccupation | -0.218 | 0.075 | 0.019 | 0.120 | -0.053 |
| | 36 PANSS G16 - Active Social Avoidance | -0.201 | 0.075 | -0.199 | -0.247 | 0.058 |

0.5 r -0.5

**Appendix 1 Figure 3. Alternative views of the behavioral PCA triplot. (A–C)** Alternative views of the triplot in **Fig. 1F** showing the relationship between the three principal axes of variation in behavior and standard clinical symptom factors. Each point represents an individual subject projected into the geometry defined by the first 3 principal components (PC). Vectors show the projections of standard symptoms factors. **(D–F)** Alternative views of the triplot in panels **A–C**, where each sphere represents the mean of each a priori clinical group. Vectors show the projections of standard symptoms factors [PANSS positive (purple), negative (blue), general (pink) symptoms and BACS cognitive performance (green)]. BPP, bipolar disorder; SADP, schizoaffective disorder; SZP, schizophrenia; CON, controls. **(G)** Heatmap of the loadings of each of the 36 symptom measures on the five significant PCs (also seen in radarplot in **Fig. 1E**). Positive loadings are indicated in red; negative loadings are shown in blue. Each PC is named based on its most strongly loaded items.

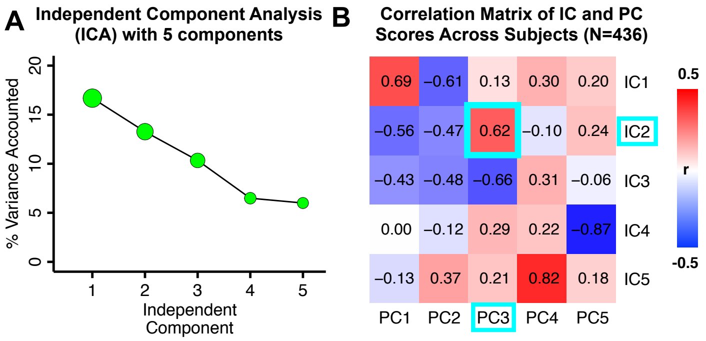


**Appendix 1 Figure 4. Independent component analysis (ICA) as an alternative method of dimensionality-reduction of symptom data. (A)** Screeplot showing the total proportion of variance explained by each independent component (IC) in a five-component solution performed across all 36 behavioral measures in 436 patients. The size of each point is proportional to the variance explained by that IC. **(B)** Correlation matrix showing correlations of individual subject scores for the 5 significant principal components (PCs) from the PCA solution shown in **Fig.** 1 and the five ICs from the ICA solution, across all 436 subjects. Neural maps for each IC are shown in **Appendix 1 - Fig.** 13.

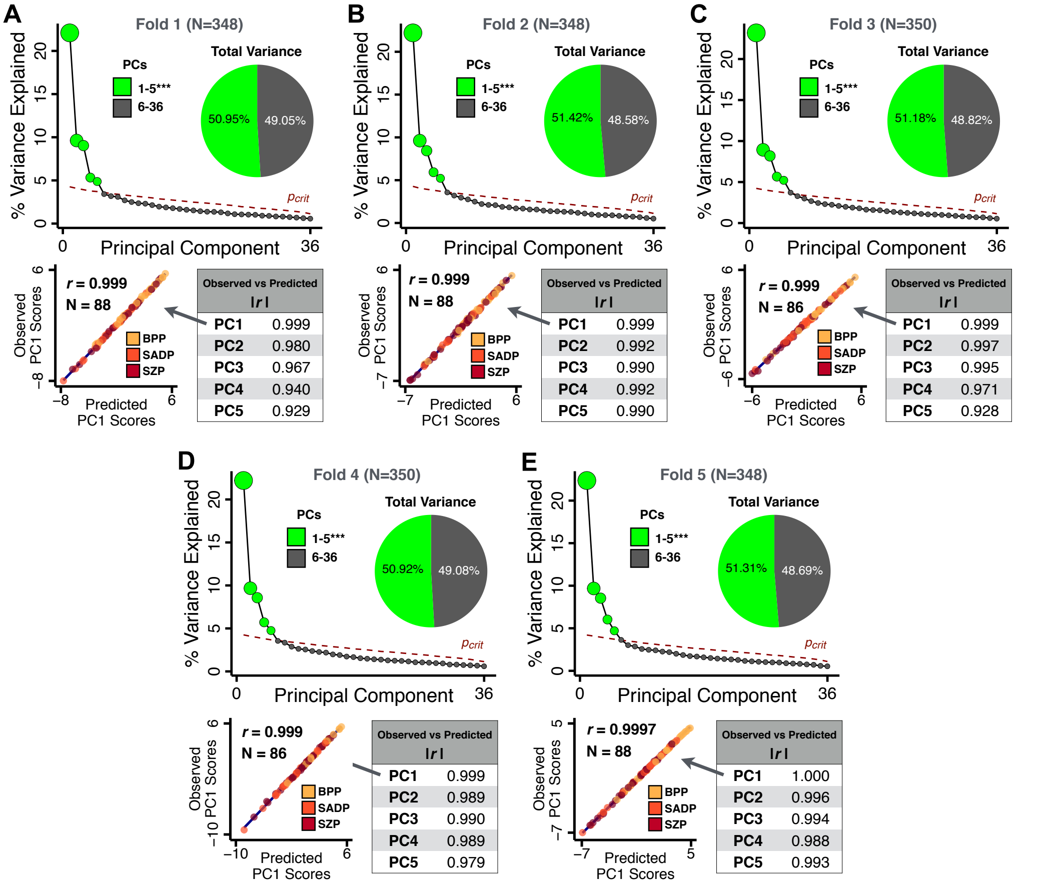


**Appendix 1 Figure 5. k-Fold cross-validation for symptom-driven principal component analysis (PCA).** These results show a 5-fold cross-validation analysis to test the stability of the PCA solution. The full patient sample was first randomly split into 5 sets and patients were randomly assigned to one of 5 subsets. Each subset of patients was then used as an independent 'test sample' in a PCA that was derived from the other 4 subsets. Screeplot shows proportion of variance explained by each of the PCs in a PCA of all 36 behavioral measures, excluding a subset of 88 patients. The number of significant PCs determined via a permutation test and the total proportion of variance explained by these PCs are all comparable to the full model shown in the main text . To obtain a 'predicted' PC score for the 88 patients in the excluded subset, the loadings from the model obtained from the other 348 patients were used. The 'observed' PC scores are the scores from the full model of the same 88 patients. The scatterplot shows that the predicted and observed scores for *PC1* are highly correlated (*r*=0.999), suggesting that the PCA solution is stable and predictive at the individual-subject level. Similarly, predicted and observed scores are highly correlated for all five PCs. **(B–E)** The results of the PCA are also highly comparable and predictive for the other four folds.

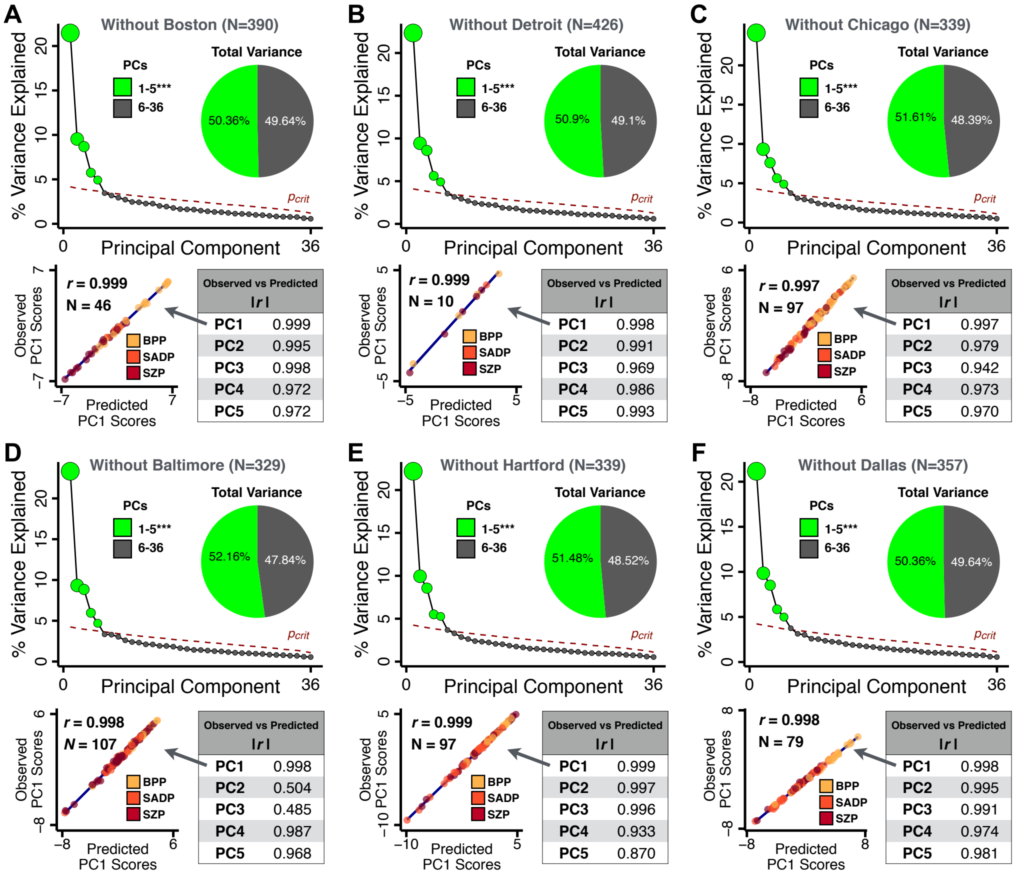


**Appendix 1 Figure 6. Leave-site-out cross-validation for symptom-driven principal component analysis (PCA).** These results show a 6-fold leave-site-out cross-validation analysis to test the stability of the PCA solution when a given site is excluded from the model. The full patient sample was first split into 6 sets according to data collection site. Each held-out site was then used as an independent 'test sample' in a PCA that was derived from the other 5 sites. **(A)** Proportion of variance explained by each of the PCs in a PCA of all 36 behavioral measures, excluding one of the six sites at which data was collected (here we excluded the Boston site). The number of significant PCs determined via a permutation test and the total proportion of variance explained by these PCs are all comparable to the full model shown in **Fig. 1C**. To obtain a 'predicted' PC score for the 46 patients in the excluded site, the loadings from the model obtained from the other 390 patients were used. The 'observed' PC scores are the scores from the full model of the same 88 patients. The scatterplot shows that the predicted and observed scores for *PC1* are highly correlated ($r$=0.999), suggesting that the PCA solution is stable and robustly predictive at the individual patient level. Predicted and observed scores are highly correlated for all five PCs. **(B–F)** The results of the PCA are also highly comparable for the other 5 sites, suggesting that possible site differences in evaluating patient symptoms or patient sample composition are not impacting the obtained PCA solutions.

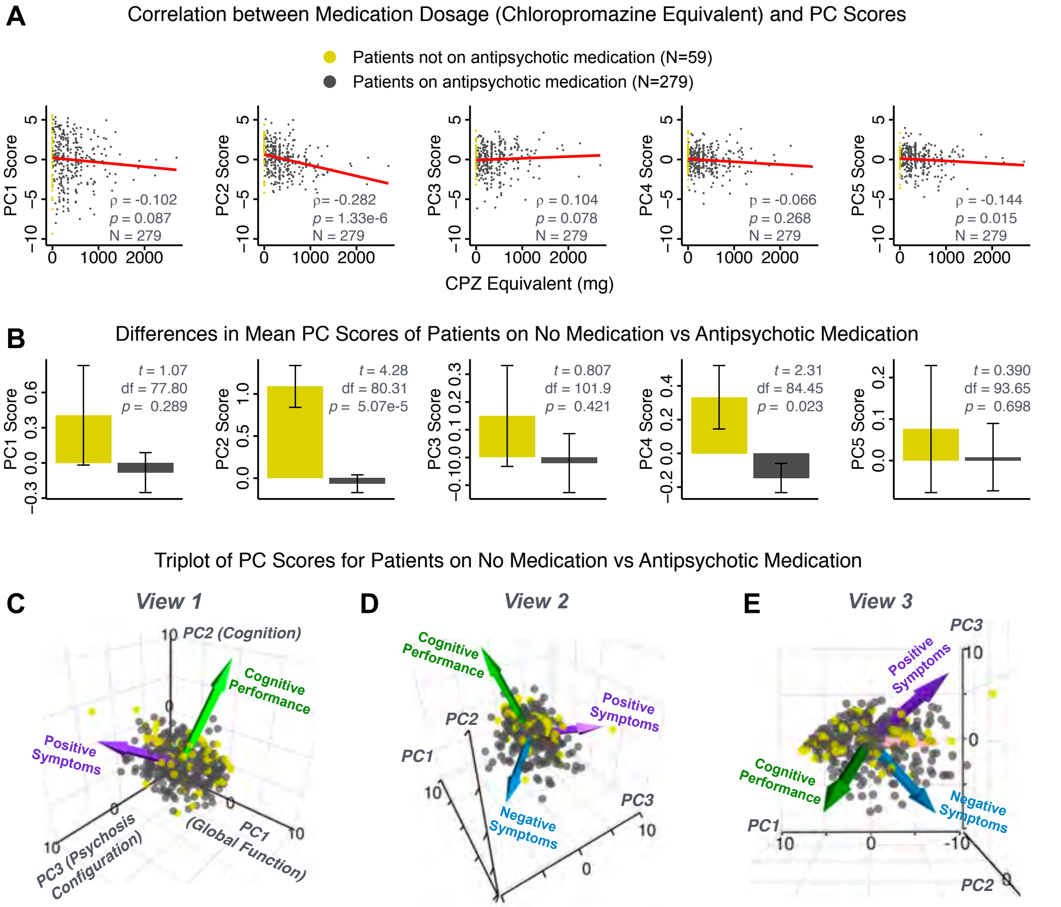


**Appendix 1 Figure 7. PCA solution is not driven by medication status or dosage.** Antipsychotic medication dosages were available for N=338 out of 436 PSD patients, including 59 patients not on antipsychotic medication. Antipsychotic dosages were converted to chlorpromazine (CPZ) equivalents (***Lencer et al., 2015***). **(A)** Spearman's $\rho$ between medication dosage (CPZ equivalents) and PC scores for each of the 5 significant principal components (PCs), for medicated patients (grey points). Patients not on antipsychotic medication are also shown (yellow points, CPZ=0mg), however they were not included in the calculation of Spearman's $\rho$ as they contain no rank information. **(B)** Bar plots show the mean PC scores of unmedicated (yellow) versus medicated (grey) PSD patients for each of the 5 PCs. Error bars show standard error of the mean. Note that only PC2 "Cognitive Performance" scores appear to show a significant relationship with medication consistently; this could be because antipsychotic medication (particularly first-generation antipsychotics) are related to symptom variance across some PCA-derived symptom dimensions but do not effectively treat cognitive deficits. Also, cognitive deficits (such as reaction time and fluency) may be exacerbated due to the neuroleptic effects of first-generation antipsychotics. **(C–E)** Alternative views of the triplot (as seen in **Appendix 1 - Fig. 3A–C**) showing the relationship between the three principal axes of variation in symptoms and aggregate scores from the PANSS and BACS symptom scales (i.e. vectors show the projections of standard symptoms factors). Each point represents an individual patient projected into the geometry defined by the first 3 principal components (PC). Each patient is colored according to medication status (yellow=unmedicated, grey=with medication). As evident from this plot there is no apparent clustering of patients according to their medication status in the 3D PCA-derived space.

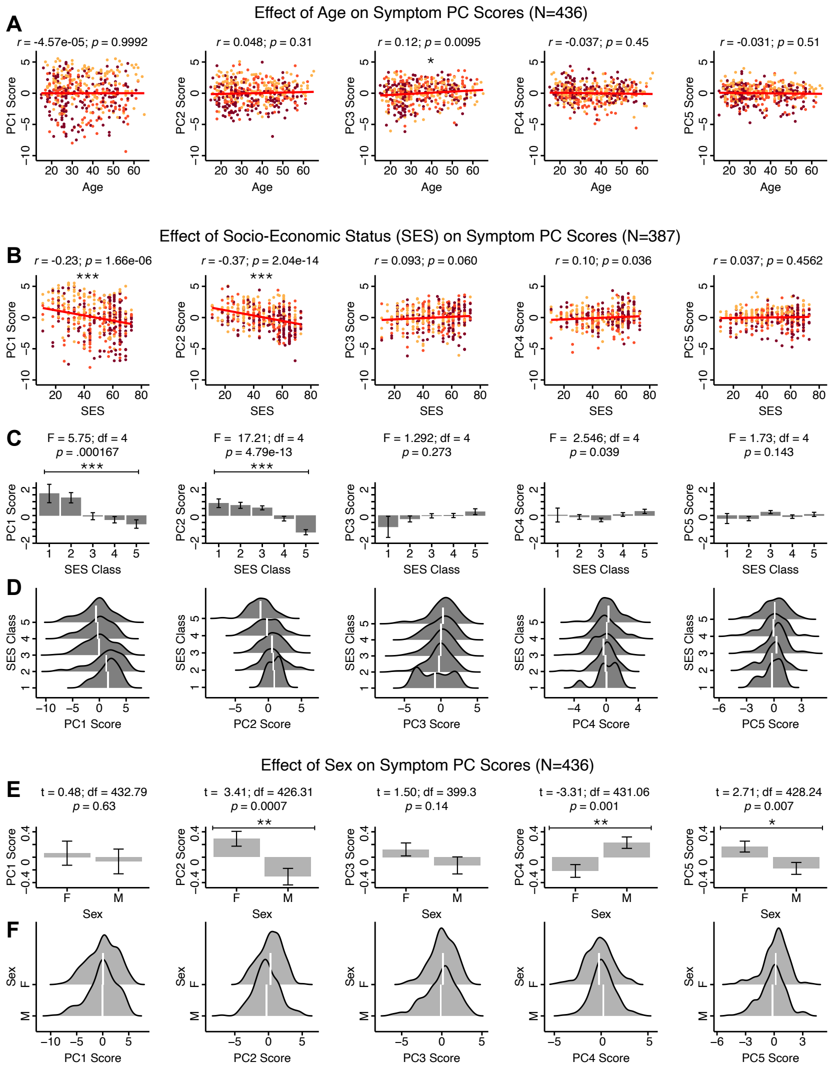


**Appendix 1 Figure 8. Effects of age, socio-economic status, and sex on symptom PCA solution. (A)** Correlations between symptom PC scores and age (years) across N=436 PSD. Pearson's correlation value and uncorrected p-values are reported above scatterplots. After Bonferroni correction, we observed a significant positive relationship between age and *PC3* score. This may be because older patients have been ill for a longer period of time and exhibit more severe symptoms along the positive *PC3* dimension. **(B)** Correlations between symptom PC scores and socio-economic status (SES) as measured by the Hollingshead Index of Social Position (***Hollingshead et al., 1975***), across N=387 PSD with available data. The index is computed as (Hollingshead occupation score * 7) + (Hollingshead education score * 4); a higher score indicates lower SES (***Padmanabhan et al., 2015***). We observed a significant negative relationship between Hollingshead index and *PC1* and PC2 scores. Lower *PC1* and *PC2* scores indicate poorer general functioning and cognitive performance respectively, which is consistent with higher Hollingshead indices (i.e. lower-skilled jobs or unemployment and fewer years of education). **(C)** The Hollingshead index can be split into five classes, with 1 being the highest and 5 being the lowest SES class (***Hollingshead et al., 1975***). Consistent with **(B)** we found a significant difference between the classes after Bonferroni correction for *PC1* and *PC2* scores. **(D)** Distributions of PC scores across Hollingshead SES classes show the overlap in scores. White lines indicate the mean score in each class. **(E)** Differences in PC scores between (M)ale and (F)emale PSD subjects. We found a significant difference between sexes in *PC2 – Cognitive Functioning*, *PC4 – Affective Valence*, and *PC5 – Agitation/Excitement* scores. **(F)** Distributions of PC scores across M and F subjects show the overlap in scores. White lines indicate the mean score for each sex.

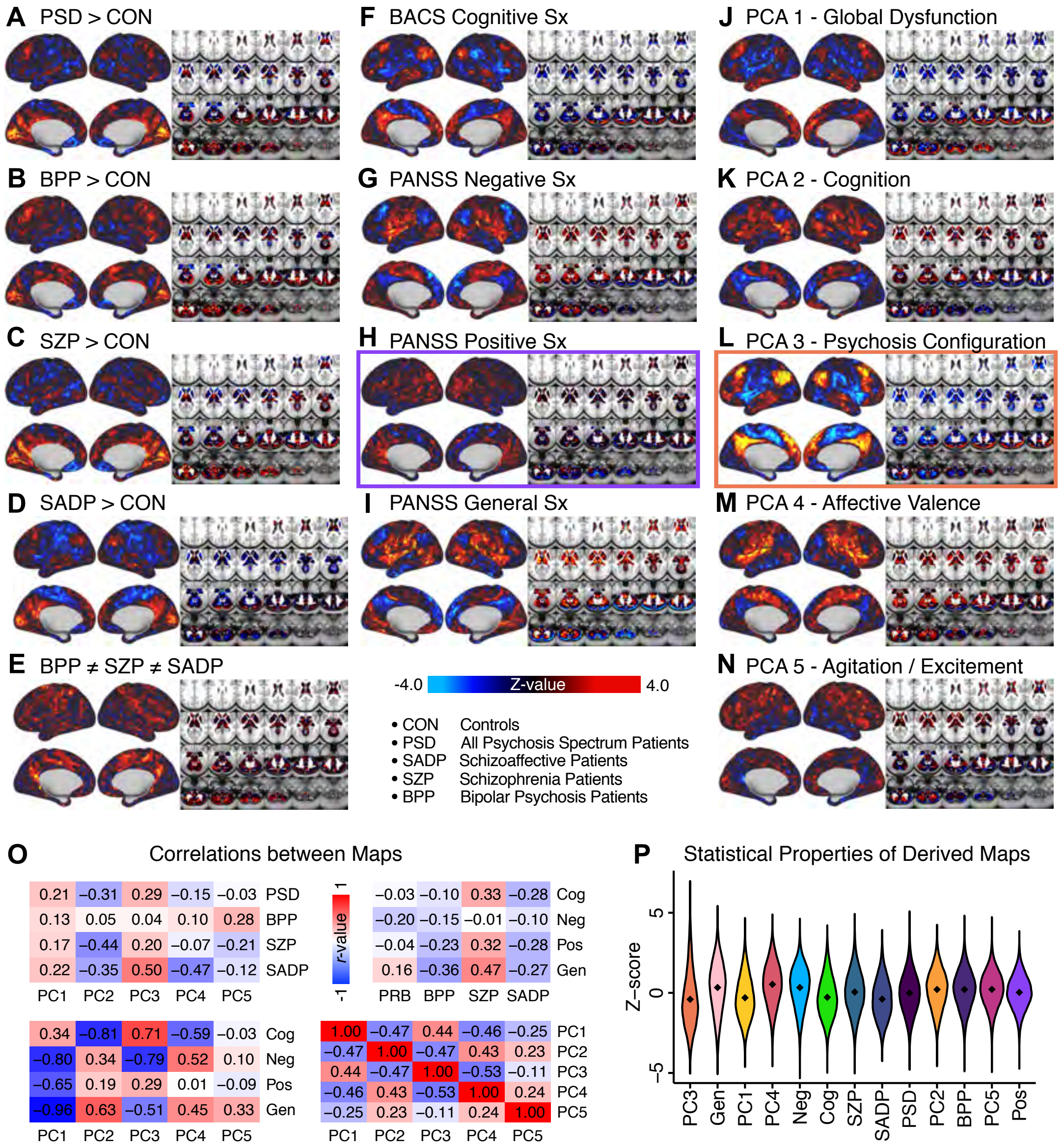


**Appendix 1 Figure 9. Similarity across *a priori*, categorical and PCA-derived brain-behavioral GBC maps.** Z-scored maps of t-test for the difference in group mean GBC between traditional diagnostic groups: **(A)** all patients (PSD) versus all healthy controls (CON); **(B)** patients with bipolar disorder (BPP) versus CON; **(C)** patients with schizophrenia (SZP) versus CON; **(D)** patients with schizoaffective disorder (SADP) versus CON. **(E)** Z-scored map of the F-test for the difference in group mean GBC between patients in all 3 diagnostic groups (BPP, SZP, SADP). Z-scored map of the regression against GBC, across all patients (PSD), of traditional symptom/behavioral scales: **(F)** BACS cognitive composite performance score; **(G)** PANSS total negative symptom score; **(H)** PANSS total positive symptom score; **(I)** PANSS total general symptom score. Z-scored map of the regression against GBC, across all patients, of data-derived behavioral dimension scores: **(J)** *PC1* score; **(K)** PC2 score; **(L)** *PC3* score; **(M)** *PC4* score; **(N)** *PC5* score. **(O)** Correlation matrices showing the similarity between brain-behavioral maps in **A-N**. **(P)** Violin plots of the distribution of Z-values in all phenotype maps. Note that although there are strong correlations between the PC maps and the *a priori* symptom maps, the statistical properties of some of the PC maps are improved (i.e. the range of Z-scores is greater, with more extreme values), suggesting a stronger mapping between neural and behavioral variation.

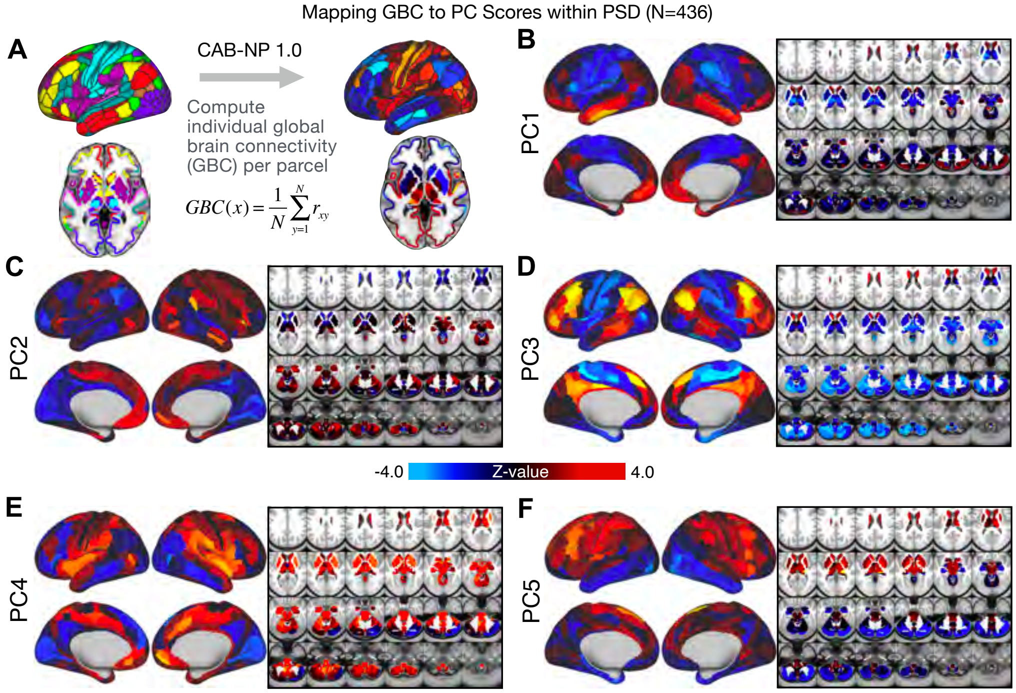


**Appendix 1 Figure 10. Parcellated symptom-neural GBC maps across all PSD patients derived from PCA dimensionality reduction of symptom measures. (A)** All of the maps shown here were parcellated at the single patient level using the Cole-Anticevic Brain Network Parcellation (CAB-NP) parcellation (***Ji et al., 2019c***), which defines functional networks and regions across cortex and subcortex that leveraged the Human Connectome Project's Multi-Modal Parcellation (MMP1.0) (***Glasser et al., 2016***; ***Ji et al., 2019c***). The final published CAB-NP 1.0 parcellation solution can be visualized via the Brain Analysis Library of Spatial maps and Atlases (BALSA) resource (https://balsa.wustl.edu/rrg5v) and downloaded from the public repository (https://github.com/ColeLab/ColeAnticevicNetPartition). **(B–F)** Relationships across all patients (N=436) at each parcel location between global brain connectivity (GBC) and PC score for each of the significant PCs. Values shown in each brain parcel is the Z-scored regression coefficient of PC score on to parcel GBC, across all 436 subjects.

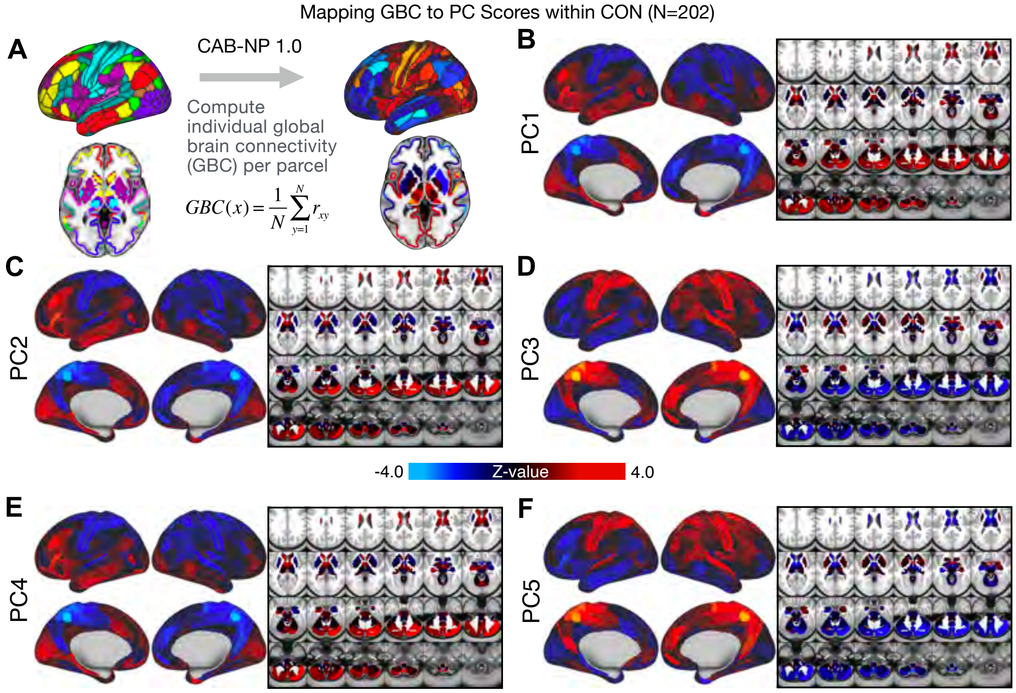


**Appendix 1 Figure 11. Parcellated symptom-neural GBC maps across all CON subjects derived from PCA dimensionality reduction of PSD symptom measures. (A)** As with PSD maps, all of the maps shown here were parcellated at the single patient level using the Cole-Anticevic Brain Network Parcellation (CAB-NP) parcellation (***Ji et al., 2019c***), which defines functional networks and regions across cortex and subcortex that leveraged the Human Connectome Project's Multi-Modal Parcellation (MMP1.0) (***Glasser et al., 2016***; ***Ji et al., 2019c***). The final published CAB-NP 1.0 parcellation solution can be visualized via the Brain Analysis Library of Spatial maps and Atlases (BALSA) resource (https://balsa.wustl.edu/rrg5v) and downloaded from the public repository (https://github.com/ColeLab/ColeAnticevicNetPartition). **(B–F)** Relationships across all controls (N=202) at each parcel location between global brain connectivity (GBC) and PC score for each of the significant PCs. Values shown in each brain parcel is the Z-scored regression coefficient of PC score on to parcel GBC, across all 202 control subjects.

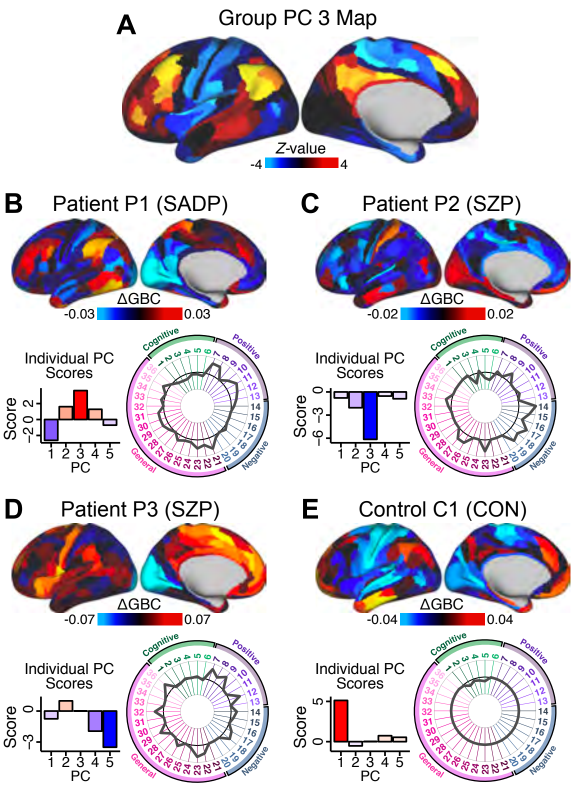


**Appendix 1 Figure 12. Individual patients exhibit complex projections into the PCA-derived brain-behavioral space (BBS) geometry (A)** The *PC3* group-level cortical map is shown here for comparison purposes. **(B)** Neural and behavioral profile for one PSD patients, “Patient P1”. Neural data shows the *Δ*GBC for Patient P1 (i.e. GBC demeaned relative to group mean of the entire PSD patient sample) to reflect the pattern of changes in their GBC relative to the ’average’ of the entire sample. The bar plot shows five behavioral PC scores for Patient P1. The radarplot shows original scores on the 36 individual symptom measures for Patient P1 (black circle indicates zero; positive values are indicated by outward deviation of the grey line). **(C–D)** Data is shown for two other individual Patients P2 & P3. **(E)**. Data is shown for one Control C1 participant. Note the difference between the symptom-neural profiles of Patient P1, a positive *PC3* ‘loading’ individual, and Patient P2, a negative *PC3* ‘loading’ individual, both of whom were diagnosed with SZP under a conventional categorical DSM approach. These data further illustrate the need to consider the complex relationships in the symptom-neural mapping and highlight that individual-level precision in BBS mapping can be obtained.

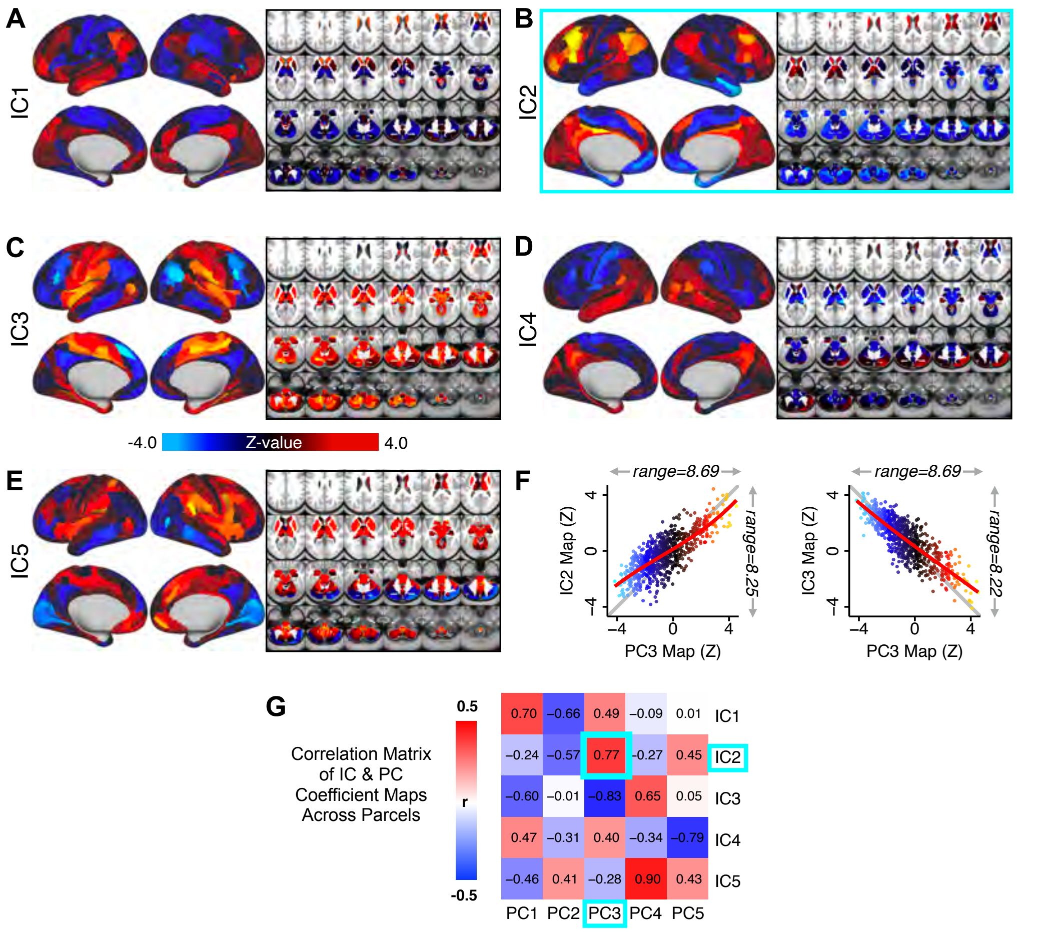


**Appendix 1 Figure 13. Independent component analysis (ICA) as an alternative method of dimensionality-reduction for symptom-neural mapping. (A–E)** Relationships across all patients (N=436) at each brain location between global brain connectivity (GBC) and IC score, for ICs 1-5. Values shown in each brain parcel is the Z-scored regression coefficient of IC score on to parcel GBC, across all 436 subjects. **(F)** Scatterplots showing the relationships across parcels between the PC3 map and the IC2 and IC3 maps. Sigmoids and a greater range (max - min) indicate an improvement in the Z-statistics of the PC3 map relative to the IC maps. **(G)** Correlation matrix showing correlations of individual parcel regression coefficients for the 5 significant PCs and the five ICs (shown in **A–E**), across all 718 neural parcels. Of note, *PC3 "Psychosis Configuration"* discovered via the PCA solution appears to be oblique to both *IC2* and *IC3*, suggesting that these two *ICs* may capture diverging tails of the *PC3* axis; however, the orthogonality of the PCA solution yielded maximally separated symptom dimensions that mapped on to unique neural variance, as shown by the superior statistics of the PC3 map versus the *IC2/IC3* maps.

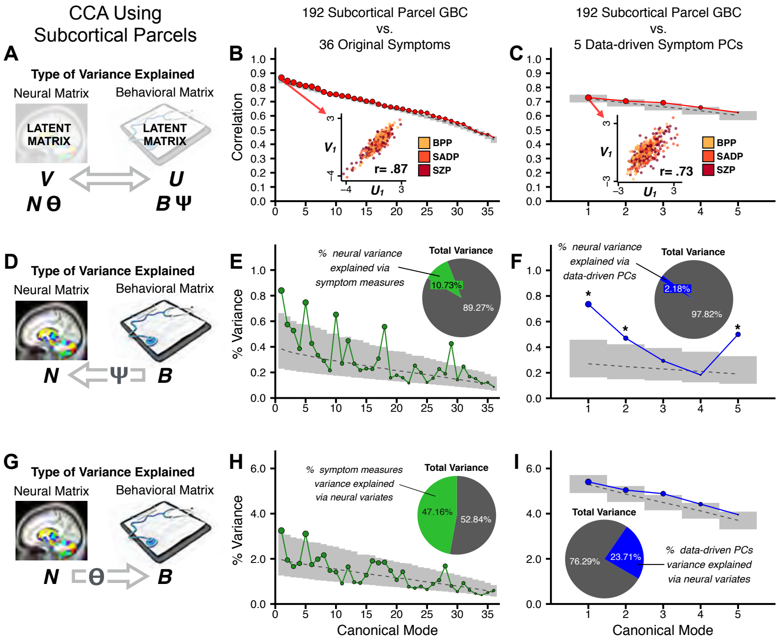


**Appendix 1 Figure 14. Canonical correlation analysis (CCA) of behavioral and subcortical neural features. (A)** As noted in the main text, CCA maximizes correlations between canonical variates (CVs), i.e. matrices $\boldsymbol{U}$ and $\boldsymbol{V}$. Here we evaluated a version of CCA using subcortical neural parcel features in relation to item-level symptom measures or dimensionality-reduced PC symptom measures. **(B)** Screeplot showing canonical modes for the CVs obtained from 192 subcortical neural features (GBC of parcels from a neurobiologically-derived functional parcellation of the subcortex) and 36 single-item PANSS and BACS symptom measures. Inset illustrates the correlation ($r$=0.85) between the CV of the first mode, $\boldsymbol{U_1}$ and $\boldsymbol{V_1}$ (note that the correlation was not driven by a separation between diagnoses). **(C)** CCA was obtained from 192 subcortical neural features and 5 low-dimensional symptom scores derived via the PCA analysis. Here all modes remained significant after FDR correction. Dashed black line shows the null calculated via a permutation test with 5,000 shuffles; grey bars show 95% confidence interval. **(D)** Correlation between the neural data matrix $\boldsymbol{N}$ and the behavioral data matrix weighted by the transformation matrix ($\boldsymbol{B\Psi}$) reflects the amount of variance in $\boldsymbol{N}$ that can be explained by the final latent neural matrix $\boldsymbol{U}$. Put differently, this transformation calculates how much of the neural variation can be explained by the latent behavioral features. **(E)** Proportion of symptom variance explained by each of the neural CVs in a CCA performed between 192 subcortical neural features and all 36 behavioral measures. Inset shows the total proportion of behavioral variance explained by the neural variates. **(F)** Proportion of total behavioral variance explained by each of the neural CVs in a CCA performed between 192 subcortical neural features and the 5 low-dimensional symptom scores derived via the PCA analysis. Dashed black line shows the null calculated via a permutation test with 5,000 shuffles; grey bars show 95% confidence interval. **(G)** Correlation between the behavioral data matrix $\boldsymbol{B}$ and the neural data matrix weighted by the transformation matrix ($\boldsymbol{N\Theta}$) reflects the amount of variance in $\boldsymbol{B}$ that can be explained by the final latent neural matrix $\boldsymbol{V}$. Put differently, this transformation calculates how much of the symptom variation can be explained by the latent neural features. **(H)** Proportion of symptom variance explained by each of the neural CVs in a CCA performed between 192 subcortical neural features and all 36 behavioral measures. Inset shows the total proportion of behavioral variance explained by the neural variates. **(I)** Proportion of total behavioral variance explained by each of the neural CVs in a CCA performed between 192 subcortical neural features and the 5 low-dimensional symptom scores derived via the PCA analysis. While CCA using symptom PCs has fewer dimensions and thus lower total variance explained (see inset), each neural variate explains a higher amount of symptom variance than seen in **F**, suggesting that CCA could be further optimized by first obtained a principled low-rank symptom solution. Dashed black line shows the null calculated via a permutation test with 5,000 shuffles; grey bars show 95% confidence interval.

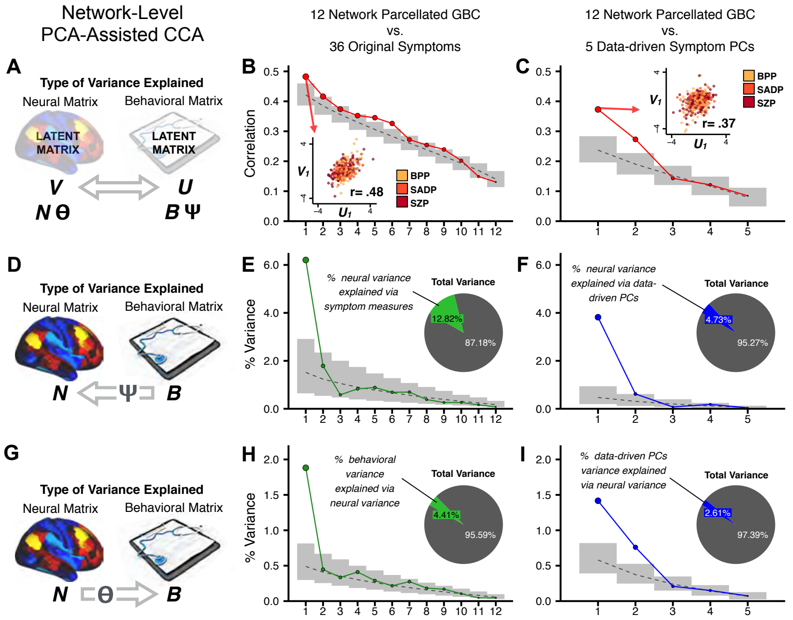


**Appendix 1 Figure 15. Canonical correlation analysis (CCA) of behavioral and network-level neural features. (A)** As noted in the main text, CCA maximizes correlations between canonical variates (CVs), i.e. matrices ***U*** and ***V***. Here we evaluated a version of CCA using network-level neural parcel features in relation to item-level symptom measures or dimensionality-reduced PC symptom measures. **(B)** Screeplot showing canonical modes for the CVs obtained from 192 subcortical neural features (GBC of parcels from a neurobiologically-derived functional parcellation of the subcortex) and 36 single-item PANSS and BACS symptom measures. Inset illustrates the correlation (*r*=0.85) between the CV of the first mode, $\boldsymbol{U_1}$ and $\boldsymbol{V_1}$ (note that the correlation was not driven by a separation between diagnoses). **(C)** CCA was obtained from 192 subcortical neural features and 5 low-dimensional symptom scores derived via the PCA analysis. Here all modes remained significant after FDR correction. Dashed black line shows the null calculated via a permutation test with 5,000 shuffles; grey bars show 95% confidence interval. **(B)** Screeplot showing the correlations between the canonical variates of a CCA performed between 12 neural features (mean GBC of 12 whole-brain networks) and all 36 behavioral measures. Dashed black line shows the null calculated via a permutation test with 5,000 shuffles; grey bars show 95% confidence interval. Inset illustrates the correlation (*r*=0.48) between the canonical variates of the first mode, $\mathbf{U}_1$ and $\mathbf{V}_1$. Note that the correlation is not driven by a separation between categorical diagnoses). **(C)** Screeplot showing the correlations between the canonical variates of a CCA performed between 12 network neural features and the 5 behavioral principal components. Dashed black line shows the null calculated via a permutation test with 5,000 shuffles; grey bars show 95% confidence interval. Inset illustrates the correlation (*r*=0.37) between the canonical variates of the first mode, $\mathbf{V}_1$ and $\mathbf{V}_1$. Note that the strength of the correlations is greatly reduced compared to the parcel-level CCA shown in **Fig. 5**. **(D)** The correlation between the neural data N and the transformed behavioral data matrix $\mathbf{B}\Psi$ reflects the amount of variance in **N** that can be explained by behavioral canonical variates **V**. **(E)** Screeplot showing the proportion of neural variance explained by each of the behavioral canonical variates in a CCA performed between 12 network neural features and all 36 behavioral measures. Dashed black line shows the null calculated via a permutation test with 5,000 shuffles; grey bars show 95% confidence interval. Inset shows the total proportion of neural variance explained by the behavioral variates. **(F)** Screeplot of the proportion of neural variance explained by each of the behavioral canonical variates in a CCA performed between 12 network neural features and the 5 PCs of behavior. **(G)** The correlation between the behavioral data **B** and the transformed neural data matrix $\mathbf{N}\Theta$ reflects the amount of variance in **B** that can be explained by neural canonical variates **U**. **(H)** Screeplot showing the proportion of behavioral variance explained by each of the neural canonical variates in a CCA performed between 12 network neural features and all 36 behavioral measures. Dashed black line shows the null calculated via a permutation test with 5,000 shuffles; grey bars show 95% confidence interval. Inset shows the total proportion of behavioral variance explained by the neural variates.t **(I)** Screeplot of the proportion of total behavioral variance explained by each of the neural canonical variates in a CCA performed between 12 network neural features and the 5 PCs of behavior. Note that although the CCA using behavioral PCs has far fewer dimensions, each neural variate explains a higher amount of total behavioral variance than neural variates in **H**, suggesting that the identified PCs of behavior capture variance with far fewer features.

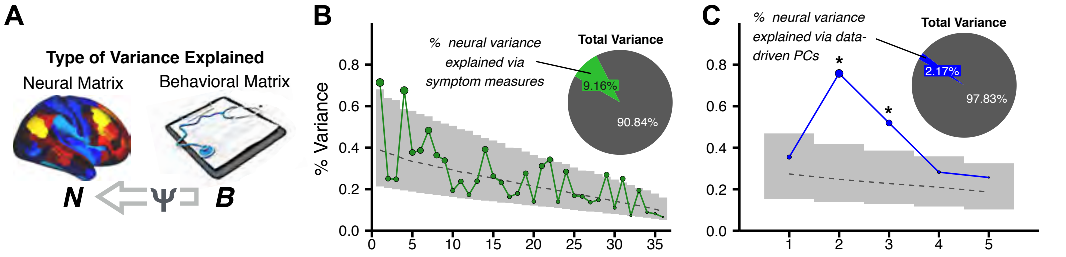


**Appendix 1 Figure 16. Canonical correlation analysis (CCA) results showing the amount of variance explained in the neural features via behavioral canonical variates. (A)** The correlation between the neural data N and the transformed behavioral data matrix ***BΨ*** reflects the amount of variance in ***N*** that can be explained by behavioral canonical variates. **(B)** Screeplot showing the proportion of neural variance explained by each of the behavioral canonical variates in a CCA performed between 180 neural features (symmetrized cortical parcel GBC) and all 36 behavioral measures. Dashed black line shows the null calculated via a permutation test with 5,000 shuffles; grey bars show 95% confidence interval. Inset shows the total proportion of neural variance explained by the behavioral variates. **(C)** Screeplot of the proportion of neural variance explained by each of the behavioral canonical variates in a CCA performed between 180 neural features (symmetrized cortical parcel GBC) and the 5 PCs of behavior.

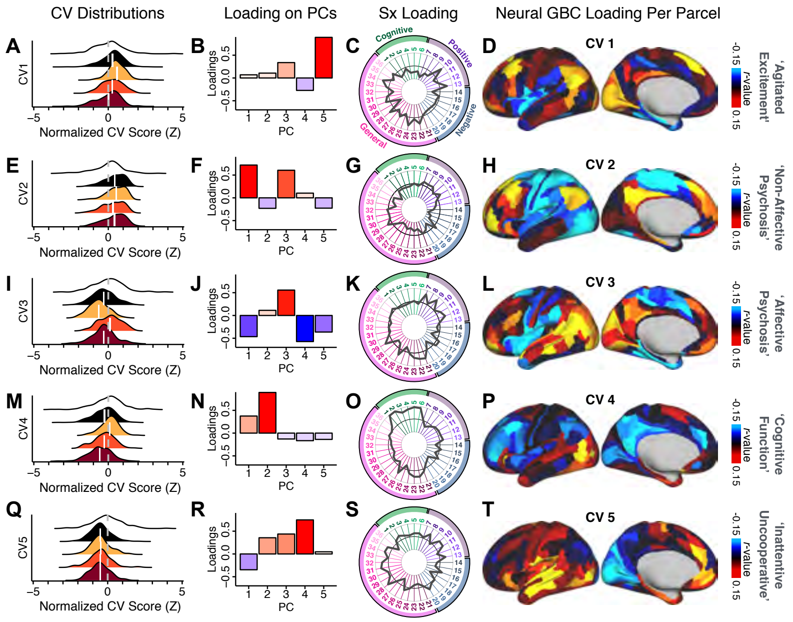


**Appendix 1 Figure 17. Canonical correlation analysis (CCA) symptom and neural configurations. (A)** Distributions of scores for the first canonical variate (*CV1*) by diagnostic group (white: controls; black: all patients; yellow: bipolar disorder with psychosis; orange: schizoaffective disorder with psychosis; red: schizophrenia). All scores are normalized to controls. **(B)** Loadings of the PC-derived symptom scores for *CV1*. **(C)** Loadings of the original item-level symptom measures for *CV1*. **(D)** Loadings of each neural parcel for *CV1*. **(E–T)** The same analyses are shown for canonical variates 2-5. As noted in the main text, these CCA effects were not stable when tested via out-of-sample cross-validation methods (see **Fig. 5**).

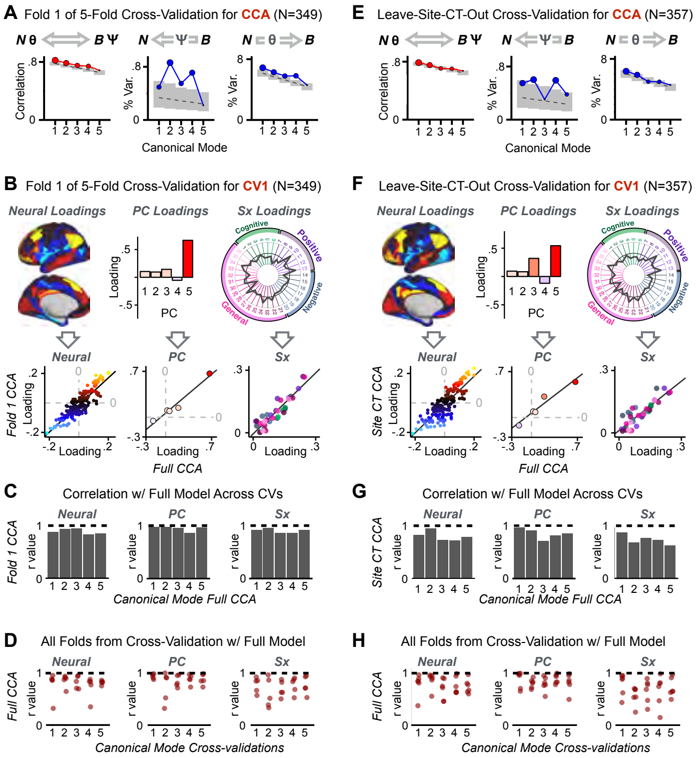


**Appendix 1 Figure 18. K-Fold cross-validation of canonical correlation analysis (CCA).** In the main text we show that CCA effects do not reproduce when tested via out-of-sample cross-validation methods (see **Fig. 5**). In contrast, here we show that using within-sample K-fold methods yields effects that appear reproducible but in fact reflect an overfit CCA solution. Panels **(A-E)** illustrate the results from a 5-fold cross-validation analysis to test the reproducibility of the CCA solution. Subjects were first randomly assigned to one of 5 subsets. Each subset of subjects was then used as an independent 'test sample' in a CCA that was derived from subjects in the other 4 subsets. **(A)** Results of CCA performed in a subsample of N=349 subjects. Left: screeplot showing the correlations between the canonical variates of a CCA performed between 180 neural features (symmetrized cortical parcel GBC) and the 5 behavioral principal components in N=349 subjects. Middle: Screeplot of the proportion of neural variance explained by each of the behavioral canonical variates in a CCA performed between 180 neural features (symmetrized cortical parcel GBC) and the 5 PCs of behavior. Right: Screeplot of the proportion of total behavioral variance explained by each of the neural canonical variates in a CCA performed between 180 neural features (symmetrized cortical parcel GBC) and the 5 PCs of behavior. Dashed black line shows the null calculated via a permutation test with 5,000 shuffles; grey bars show 95% confidence interval. Note that the outcome of this CCA is highly similar to the full sample CCA shown in the main text, but it is not similar at all to the split-half cross-validation effects, which reveal how overfit CCA effects seem to be with this number of features **Fig. 5**. **(B)** Comparison of the first canonical variate (CV1) in a CCA performed in a subsample of N=349 subjects with *CV1* from the full model performed with N=436 subjects. Left: neural factor loadings for *CV1* obtained from the subsample (Fold 1) CCA and from the full sample CCA are highly correlated, at r=0.85. Middle: Behavioral factor loadings are also highly correlated between the Fold 1 CCA and the full CCA, at r=0.85. Right: Additionally, absolute values of individual symptom measure loadings associated with *CV1* are highly correlated between the Fold 1 CCA and the full CCA. **(C)** Summary of correlation values between Fold 1 CCA and full CCA results, as in panel **B**, across all 5 CVs. Note that the neural and behavioral factor loadings as well as individual symptom measure loadings are highly preserved between the sample CCA and the full model. **(D)** Summary of correlation values between all 5 subsample CCAs and the full sample CCA, for all 5 CVs. Each of the 5 CVs is plotted along the X-axis; each point represents the correlation between one of the 5-fold subsample CCAs and the full CCA, hence there are 5 points (in the 5-fold cross-validation) for each CV. **(E)** Results of a CCA performed in a subsample of N=357 subjects, with subjects from one site left out (leave-one-site-out, LOSO). Panels as described in **A**. **(F)** Comparison of the first canonical variate (CV1) in a LOSO CCA and the *CV1* from the full model performed with N=436 subjects. Panels as described in **B**. **(G)** Summary of correlation values between the LOSO CCA and full CCA results, as in panel **B**, across all 5 CVs. Panels as described in **C**. **(H)** Summary of correlation values between all 6 LOSO CCAs and the full sample CCA, for all 5 CVs. Panels as described in **D**; each point represents the correlation between one of the 6 LOSO CCAs and the full CCA.

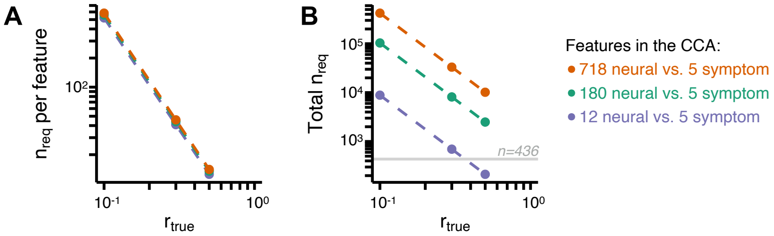


**Appendix 1 Figure 19. Multivariate power analysis for CCA.** Sample sizes were calculated according to (***Helmer et al., 2020***), see also **Methods** and https://gemmr.readthedocs.io/en/latest/. We computed the multivariate power analyses for three versions of CCA reported in this manuscript: i) 718 neural vs. 5 symptom features; ii) 180 neural vs. 5 symptom features; iii) 12 neural vs. 5 symptom features. **(A)** At different levels of features, the ratio of samples (i.e. subjects) required per feature to derive a stable CCA solution remains approximately the same across all values of $r_{true}$. As discussed in (***Helmer et al., 2020***), at $r_{true} = 0.3$ the number of samples required per feature is about 40, which is much greater than the ratio of samples to features available in our dataset. **(B)** The total number of samples required ($n_{\mathrm{req}}$) for a stable CCA solution given the total number of neural and symptom features used in our analyses, at different values of $r_{true}$. In general these required sample sizes are much greater than the N=436 (horizontal grey line) PSD in our present dataset, consistent with the finding that the CCA solutions computed using our data were unstable. Notably, the '12 vs. 5' CCA assuming $r_{true} = 0.3$ requires only $\sim$700 subjects, which is closest to the N=436 used in the present sample. This may be in line with the observation of the CCA with 12 neural vs 5 symptom features (**Appendix 1 - Fig. 15C**) that the canonical correlation ($r = 0.38$ for CV1) clearly exceeds the 95% confidence interval, and may be closer to the true effect. However, to to confidently detect effects in such an analysis (particularly if $r_{true}$ is actually less than 0.3), a larger sample would likely still be needed.

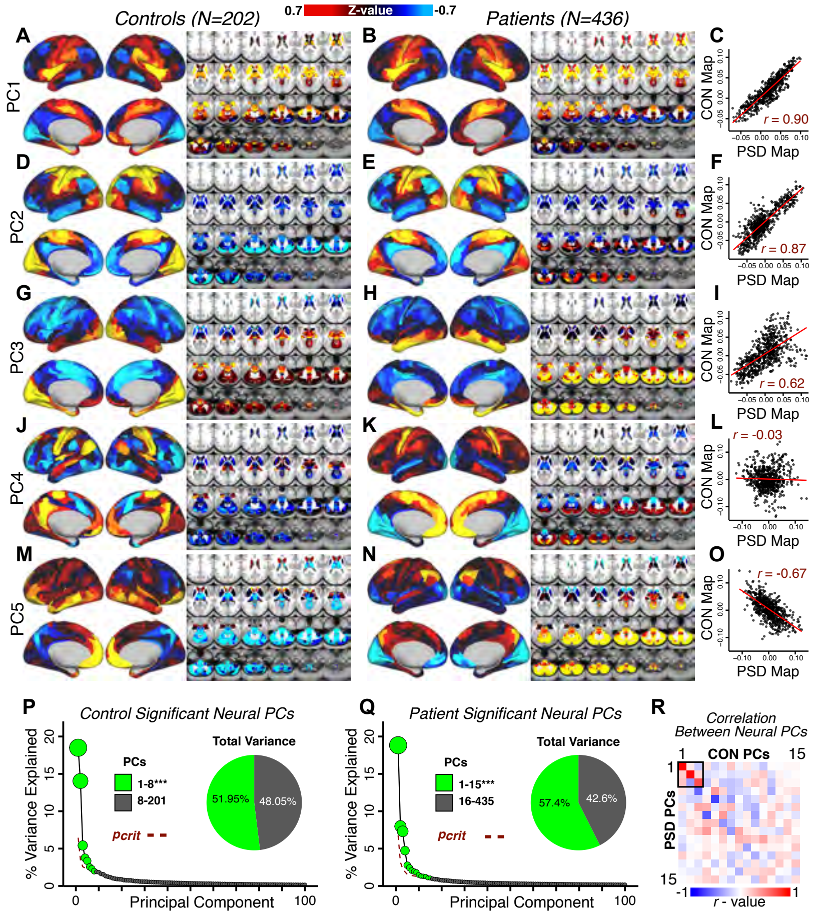


**Appendix 1 Figure 20. Principal component analysis (PCA) of neural features in control and patient subjects. (A)** Results of PCA performed on neural features (718 whole-brain parcel GBC) across all control subjects (N=202). **(B)** Results of PCA performed on neural features (718 whole-brain parcel GBC) across all patient subjects (N=436). **(C)** Note that the first neural PC in control subjects (panel A) and in patient subjects (panel **B**) are highly similar ($r$=0.90 across parcels). This may reflect a component of neural variance common across all humans. **(D–I)** The second and third PCs are also similar between control and patient subjects. **(J–L)** The fourth PC is markedly dissimilar between control and patient subjects, suggesting that this component may reflect neural variance which deviates in individuals with psychiatric symptoms. **(M–O)** The fifth PC is also highly dissimilar between controls and patients, also possibly reflecting diagnosis-relevant differences in neural variance. **(P)** Screeplot showing the total proportion of variance explained by the first 100 PCs from the PCA performed across all 718 neural features in 202 control subjects. The size of each point is proportional to the variance explained by that PC. The first 8 PCs (green) were determined to be significant using a permutation test. Inset shows the proportion of variance both accounted and not accounted for by the 8 significant PCs. Together, these 8 PCs capture 51.95% of the total variance in neural GBC in the sample. **(Q)** Screeplot showing the total proportion of variance explained by the first 100 PCs from the PCA performed across all 718 neural features in 436 patient subjects. The size of each point is proportional to the variance explained by that PC. The first 15 PCs (green) were determined to be significant using a permutation test. Inset shows the proportion of variance both accounted and not accounted for by the 15 significant PCs. Together, these five PCs capture 57.4% of the total variance in neural GBC in the sample. **(R)** Correlations between the first 15 neural PCs in controls (CON) and patients (PSD). The first 3 PCs are common across both control and patient subjects, possibly reflecting common components of human neural variance. After this, the PCs diverge, possibly reflecting differences in neural variance of healthy individuals and those with psychiatric symptoms.

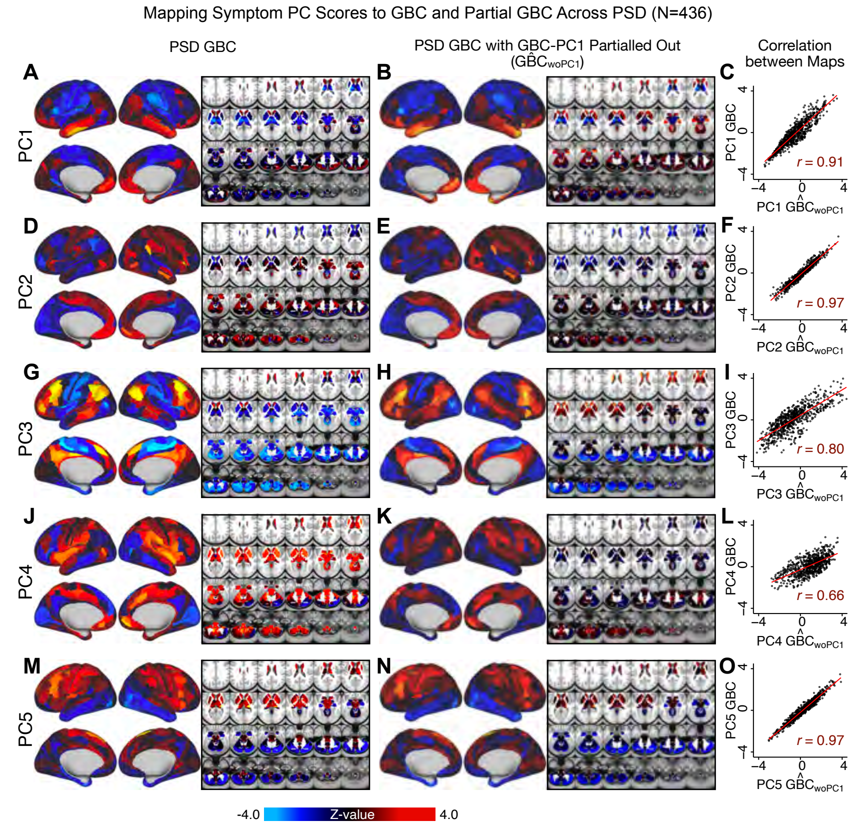


**Appendix 1 Figure 21. Comparison between the PSD $\beta_{PC}GBC$ maps computed using GBC and GBC with the first neural PC parsed out.** If a substantial proportion of neural variance is not be clinically relevant, then removing the shared neural variance between PSD and CON should not drastically affect the reported symptom-neural univariate mapping solution, because this common variance will not map to clinical features. We therefore performed a PCA on CON and PSD GBC to compute the shared neural variance (see **Methods**), and then parsed out the first GBC-PC from the PSD GBC data ($G\hat{B}C_{woPC1}$). We then reran the univariate regression as described in **Fig. 3**, using the same five symptom PC scores across 436 PSD. **(A)** The $\beta_{PC1}GBC$ map, also shown in **Appendix 1 - Fig. 10**. **(B)** The first GBC-PC accounted for about 15.8% of the total GBC variance across CON and PSD. Removing GBC-PC1 from PSD data attenuated the $\beta_{PC1}GBC$ statistics slightly (not unexpected as the variance was by definition reduced) but otherwise did not strongly affect the univariate mapping solution. **(C)** Correlation across 718 parcels between the two $\beta_{PC1}GBC$ map shown in **A** and **B**. **(D-O)** The same results are shown for $\beta_{PC2}GBC$ to $\beta_{PC5}GBC$ maps.

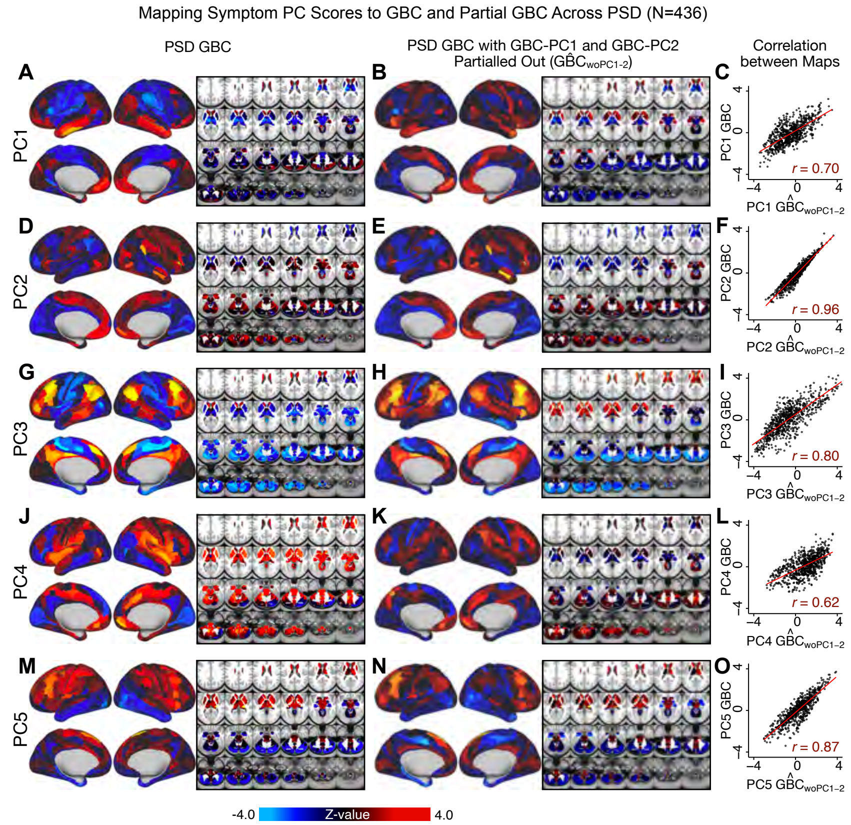


**Appendix 1 Figure 22. Comparison between the PSD $\beta_{PC}GBC$ maps computed using GBC and GBC with the first two neural PCs parsed out.** We performed a PCA on CON and PSD GBC and then parsed out the first three GBC-PC from the PSD GBC data ($G\hat{B}C_{woPC1-2}$, see **Methods**). We then reran the univariate regression as described in **Fig.** 3, using the same five symptom PC scores across 436 PSD. **(A)** The $\beta_{PC1}GBC$ map, also shown in **Appendix 1 - Fig.** 10. **(B)** The second GBC-PC accounted for about 9.5% of the total GBC variance across CON and PSD. **(C)** Correlation across 718 parcels between the two $\beta_{PC1}GBC$ map shown in **A** and **B**. **(D-O)** The same results are shown for $\beta_{PC2}GBC$ to $\beta_{PC5}GBC$ maps.

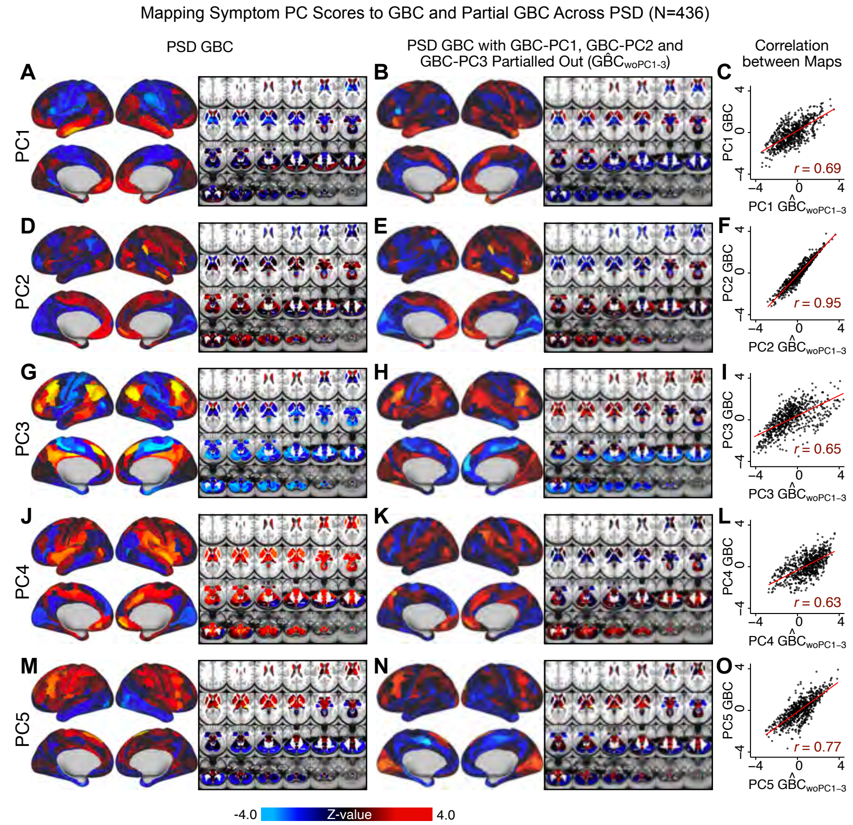


**Appendix 1 Figure 23. Comparison between the PSD $\beta_{PC}GBC$ maps computed using GBC and GBC with the first three neural PCs parsed out.** We performed a PCA on CON and PSD GBC and then parsed out the first three GBC-PC from the PSD GBC data ($G\hat{B}C_{woPC1-3}$, see **Methods**). We then reran the univariate regression as described in **Fig.** 3, using the same five symptom PC scores across 436 PSD. **(A)** The $\beta_{PC1}GBC$ map, also shown in **Appendix 1 - Fig.** 10. **(B)** The second GBC-PC accounted for about 9.5% of the total GBC variance across CON and PSD. **(C)** Correlation across 718 parcels between the two $\beta_{PC1}GBC$ map shown in **A** and **B**. **(D-O)** The same results are shown for $\beta_{PC2}GBC$ to $\beta_{PC5}GBC$ maps.

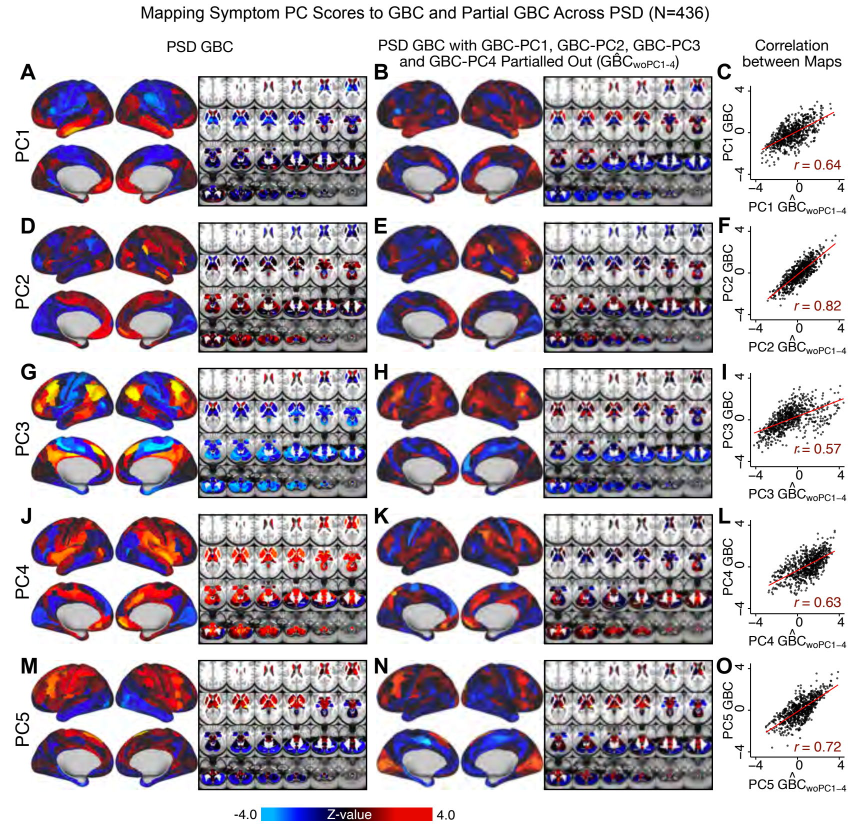


**Appendix 1 Figure 24. Comparison between the PSD $\beta_{PC}GBC$ maps computed using GBC and GBC with the first four neural PCs parsed out.** We performed a PCA on CON and PSD GBC and then parsed out the first four GBC-PC from the PSD GBC data ($G\hat{B}C_{woPC1-4}$, see **Methods**). We then reran the univariate regression as described in **Fig.** 3, using the same five symptom PC scores across 436 PSD. **(A)** The $\beta_{PC1}GBC$ map, also shown in **Appendix 1 - Fig.** 10. **(B)** The second GBC-PC accounted for about 9.5% of the total GBC variance across CON and PSD. **(C)** Correlation across 718 parcels between the two $\beta_{PC1}GBC$ map shown in **A** and **B**. **(D-O)** The same results are shown for $\beta_{PC2}GBC$ to $\beta_{PC5}GBC$ maps.

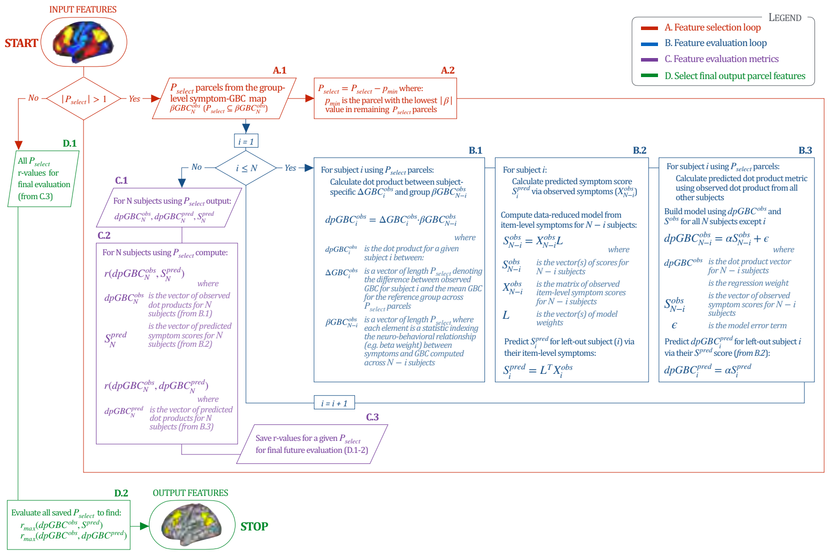


**Appendix 1 Figure 25. Workflow for optimizing the neural feature selection for a predictive symptom-neural subject specific model.** We optimized the selection of a subset of neural parcels from the group $\beta GBC_N^{obs}$ map (where $N$ is the total number of subjects) that could be predicted via a single symptom score across subjects (e.g. for the *PC3* beta map and *PC3* symptom scores). (**START**) We start with $P_{select}$ set as the full parcellated map,$P_{select} = \beta GBC_N^{obs}$. Hence initially there are 718 elements in the vector $P_{select}$, each representing the value of a parcel ($|P_{select}| = 718$). Next, since $|P_{select}| > 1$, the selection process enters the **A. Feature Selection Loop** (in red). Selects subset of parcels $P_{select}$ (**A.1**) for which the predictive model will be evaluated. This loop iteratively removes parcels from the $P_{select}$ vector (**A.2**) and each time passes the updated $P_{select}$ as input in the Feature Evaluation Loop (in blue). **B. Feature Evaluation Loop** (shown in blue). Using a multi-step algorithm, this loop produces the set of metrics to be evaluated at the final output using a leave-one-subject out approach. **B.1**: For each subject $i$, this first step is to compute the "observed" dot product metric $dpGBC_i^{obs}$ for that subject. This is the dot product between $\Delta GBC_i^{obs}$ and $\beta GBC_{N-i}^{obs}$ using the $P_{select}$ features. $\Delta GBC_i^{obs}$ is the vector of the parcel-wise difference between subject $i$'s GBC map and the group mean GBC map; $\beta GBC_{N-i}^{obs}$ denotes group-level relationship between symptom variation and GBC excluding subject $i$. This group $\beta GBC_{N-i}^{obs}$ is used as a reference for the dot-product. **B.2**: Next for subject $i$ we compute their predicted symptom score $S_i^{pred}$. First, a symptom-prediction model is built using $X_{N-i}^{obs}$ (all item-level symptom measures excluding subject $i$). This yields the “observed” model scores and coefficients excluding subject $i$ ($S_{N-i}^{obs}$). The symptoms for excluded subject $i$ are then used to obtain the predicted symptom score $S_i^{pred}$, as done in **Fig. 6A**. **B.3**: Lastly, for excluded subject $i$ the loop computes the "predicted" dot product metric $dpGBC_i^{pred}$ to compare it to the "observed" value for model evaluation. Here, a regression model is computed relating $dpGBC_{N-i}^{obs}$ to $S_{N-i}^{obs}$, excluding subject $i$. This yields regression coefficients $\alpha$, which are then used to predict the $dpGBC_i^{pred}$ for the excluded subject $i$ using their predicted symptom scores $S_i^{pred}$. Once $i > N$ (i.e. $dpGBC_i^{pred}$ and $S_i^{pred}$ have been computed for all subjects), the algorithm moves to step "C". **C. Feature Evaluation Metrics** (shown in purple). **C.1**: This step captures feature evaluation metrics across N subjects for the model that was computed using the selected features ($P_{select}$). Specifically, three vectors (outputs from "B") are computed: $dpGBC_N^{obs}$ (the vector of observed $dpGBC$ values for all $N$ subjects); $dpGBC_N^{pred}$ (the vector of predicted $dpGBC$ values for all $N$ subjects); and $S_N^{pred}$ (the vector of predicted model symptom scores for all $N$ subjects). **C.2**: Generates two feature evaluation metrics across the entire sample ($r(dpGBC_N^{obs}, S_N^{pred})$ and $r(dpGBC_N^{obs}, dpGBC_N^{pred})$). **C.3**: The feature evaluation metrics for the selected $P_{select}$ are generated, such that they can be evaluated across all $P_{select}$ iterations (red loop). **D. Final Feature Selection** (in green). This step evaluates the metrics computed for all $P_{select}$ iterations, starting from from $|P_{select}| = 718$ to $|P_{select}| = 1$. **D.1**: The input is the vector of $r(dpGBC_N^{obs}, S_N^{pred})$ and the vector of $r(dpGBC_N^{obs}, S_N^{pred})$ for all 718 models. **D.2**: The most predictive feature subset is obtained by finding the model for which $r(dpGBC_N^{obs}, S_N^{pred})$ and $r(dpGBC_N^{obs}, dpGBC_N^{pred})$ were maximized, which leads to the final output features (**STOP**). *Note*: that the algorithm imposed a threshold of $|P_{select}| > 30$ to ensure that the correlation metrics were not unstable based on a small feature set.

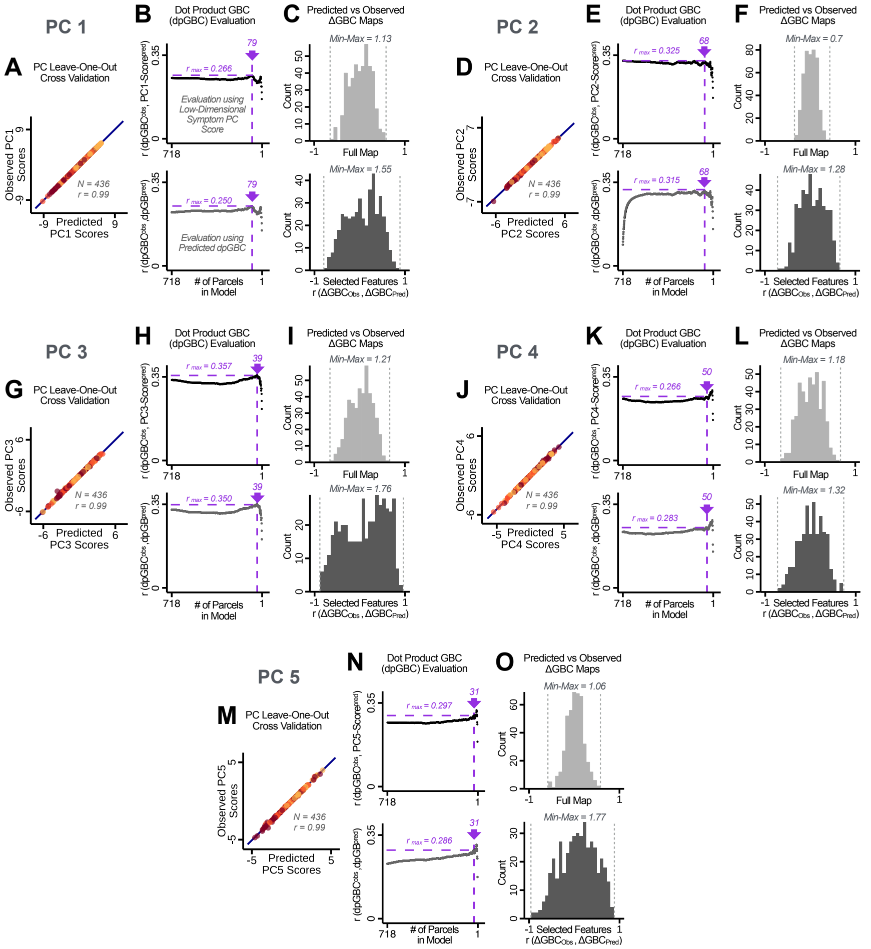


**Appendix 1 Figure 26. Optimizing neural feature selection to inform statistically reliable single-subject prediction via low-dimensional symptom scores only. (A)** Individual PC-derived symptom scores for *PC1* can be robustly predicted, as shown by leave-one-out cross-validation for the symptom PCA analyses. Here for the leave-one-out cross-validation, a PCA was computed in all except one subject (N=435) and the PCA loadings were then used to compute the predicted PC scores for the left-out subject. This was repeated such that predicted scores were calculated for every subject, using loadings from a PCA computed in the other 435 subjects. Observed scores were obtained from a PCA computed using the full sample of subjects (N=436). Top panel: Scatterplot at left shows the correlation between each subject's predicted *PC1* score from leave-one-out PCA model and their observed *PC1* score from the full-sample PCA model, *r*=0.99. Red line shows perfect prediction, *r*=1. **(B)** Top panel indicates that PC symptom scores can yield robust neural feature selection. *Δ*GBC denotes an individual subject GBC demeaned parcel-wise relative to the group mean GBC. The process is described in detail in the **Methods** and in **Appendix 1 - Fig.** 25. The correlation between *PC1* behavioral score and [ΔGBC · *PC1* beta] across all 436 subjects was maximal for $P = 79$, at *r*=0.27 (purple arrow). Bottom panel: For each value of *p*, the regression was performed in 435 subjects and was then used to predict the [ΔGBC · *PC1* beta] of the selected parcels in the left-out subject. The mean correlation between the predicted [ΔGBC · *PC1* beta] and the actual observed [ΔGBC · *PC1* beta] across all 436 subjects was highest for $P = 79$, with *r*=0.19 across all subjects, highly consistent with the across-subject analysis in the top panel. **(C)** Comparing the correlation between $\Delta GBC_{Obs}$, $\Delta GBC_{Pred}$ for the full map ($P = 718$) versus the selected map ($P = 79$) shows that the predictability of $\Delta$ GBC is improved by using the selected parcels in the selected map.s **(D–O)** Similar results for PCs 2-5. Note that for *PC4* and *PC5* a local maximum ($P = 50$ and $P = 31$ respectively) was selected after imposing a threshold of $P > 30$ to ensure that the correlation metrics used for evaluation were not made unstable from a small number of parcels in computing the $dpGBC$.

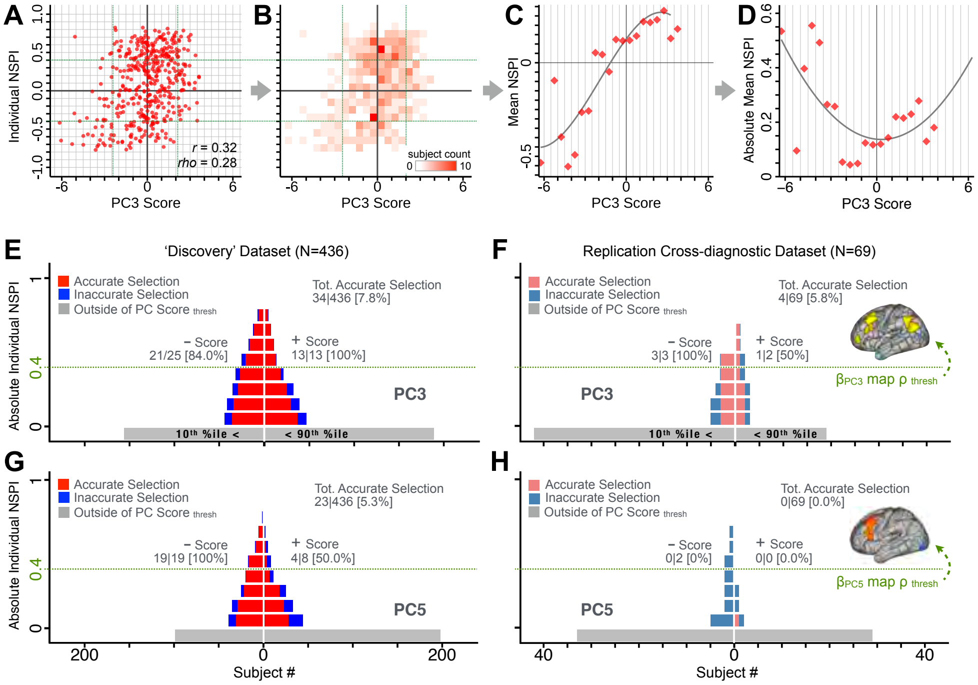


**Appendix 1 Figure 27. Evaluating patient-specific similarity to derived symptom-neural targets via data-reduced symptom scores.** **(A)** Scatterplot of *PC3* symptom score (X-axis) versus *PC3* neural similarity prediction index (NSPI, Y-axis) for all 436 PSD subjects. The NSPI is defined as the Spearman's correlation between $\Delta GBC^{obs}$ and the $\beta_{PC3} GBC^{obs}$ map of the maximally-predictable "selected" $P = 39$ parcels. Alternative metrics are shown in **Appendix 1 - Fig.** 28. **(B)** Bins across axes express subject counts within each cell as a heatmap, indicating a high similarity between symptom PC score and *PC3* NSPI for a number of patients. **(C)** Mean NSPI is computed for a given bin along the the X-axis to visualize patient clustering. Note the sigmoidal shape of the distribution reflecting greater neural similarity at more extreme values of the *PC3* score. **(D)** The absolute value of the mean NSPI reflects the magnitude irrespective of neural similarity direction. This highlights a quadratic effect, showing that patients with higher *PC3* symptom scores (either positive/negative) exhibited higher neural correspondence of their maps with the target neural reference map. **(E)** Using the NSPI and PC scores we demonstrate one possible brain-behavioral patient selection strategy. We first imposed a PC score symptom threshold to select patients at the extreme tails (i.e. outside of the $10th - 90th\%tile$ behavioral range [>+2.17 or <-2.41]). Note that this patient selection strategy excludes patients (shown in grey) below the PC symptom score threshold. This yielded n=38 patients. Next, for each patient we predicted the sign of their individual NSPI based on their individual PC3 score, which served as the basis for the neural selection. Next, at each NSPI threshold we evaluated the proportion of patients correctly selected until there were no inaccurately selected patients in at least one PC3 direction (green line or higher). The number of accurately (A) vs. inaccurately (I) selected patients within each bin is shown in red and blue respectively. Note that as the neural $\rho$ threshold increases the A/I ratio increases. **(F)** The neural and behavioral thresholds defined in the "discovery" sample were applied to an independent "replication" dataset (N=69, see **Methods**), yielding a similar final proportion of accurately selected patients. **(G)** The same brain-behavioral patient selection strategy was repeated for *PC5* in the discovery sample (thresholds of $10^{th}\%tile$=-1.89 and $90^{th}\%tile$= +1.47; NSPI threshold of $\rho$=0.4 optimized for PC5). Results yielded similar A/I ratios as found for PC3. **(H)** The neural and behavioral thresholds for PC5 defined in the discovery sample were applied to the replication sample. Here the results failed to generalize due true clinical differences between the discovery and replication samples.

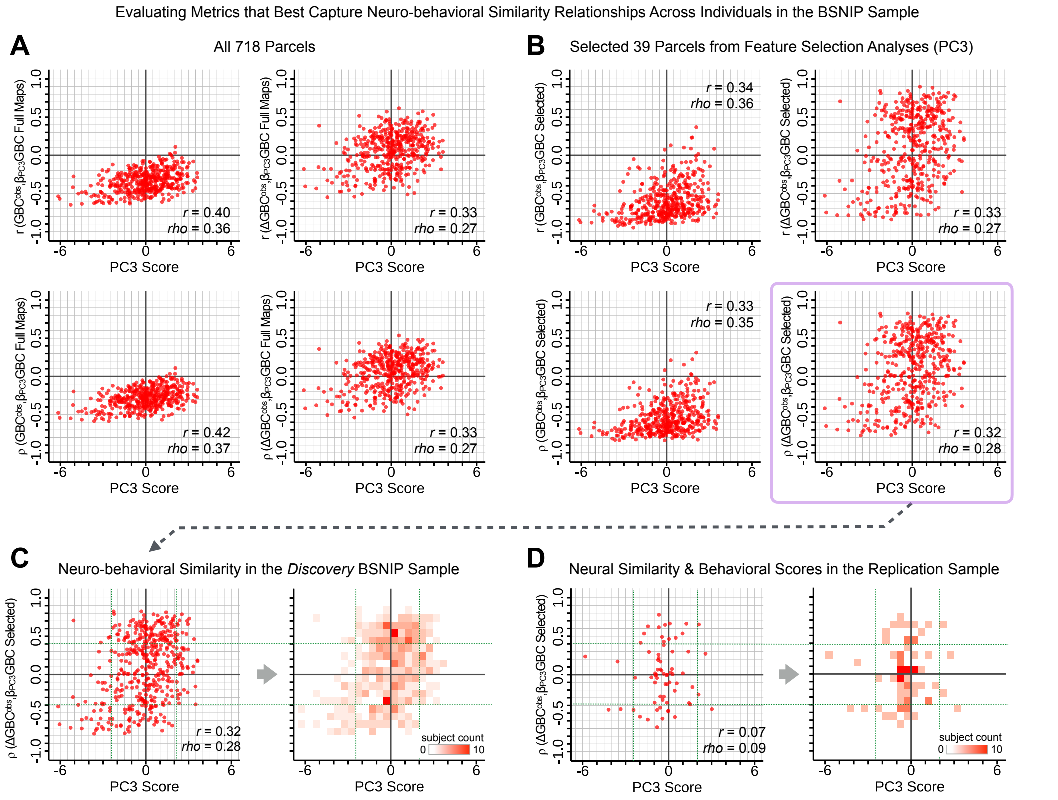


**Appendix 1 Figure 28. Metrics quantifying the relationship between individual-level neural effects relative to the group mean and individual-level PC-derived symptom scores.** The relationship between various symptom-neural similarity metrics and the behavioral *PC3* symptom scores across N=436 PSD patients are shown. The goal was to evaluate metrics that maximize the spread of the Y-axis, while maintaining a centered distribution of scores around zero to yield an interpretable symptom-neural variation pattern across subjects. Specifically, along the Y-axis we examined either the Pearson's correlation (top panels in A and B) or Spearman's $\rho$ (bottom panels in A and B) between two neural maps at the individual patient level: i) the group-level $\beta_{PC3}GBC$ map and ii) an individual subjects' raw GBC map which can be expressed as either a raw map (left panels in A and B) or as $\Delta GBC^{obs}$ relative to the group mean map (right panels in A and B). **(A)** These four panels show group-level symptom-neural relationships for sample using the entire 718-parcel neural map. As noted, the neural similarity metrics on each Y-axis show the relationship between individual-patient neural GBC (or $\Delta GBC^{obs}$) and the group-level $\beta_{PC3}GBC$ map across 718 parcels. Note that the *r* and *rho* values reported in the corner of each plot reflect the *group-level* relationship between the neural similarity metric and the individual-level PC3 symptom score computed across across 436 patients. **(B)** These four parcels show group-level symptom-neural relationships using only the selected 39 *PC3* parcels derived from the feature selection workflow (**Appendix 1 - Fig.** 25). The metric used in the main text is the $\rho$ within selected parcels (shown with a light purple envelope) because the spread along Y-axis was maximized and the values centered around zero for both the x and the Y-axis. Furthermore, the $\rho$ metric is desirable given violations of normality that the $r$ metric assumes. **(C)** The neural-behavioral similarity plot for the Discovery BSNIP sample was binned by $\rho = 0.1$ & $PC3_{score} = 0.5$ to provide a visual intuition for patient segmentation across both the neural and symptom indices. For patients with highly positive or negative scores, the neuro-behavioral relationship was robust. Conversely, patients with a low absolute *PC3* score showed a weak relationship with symptom-relevant neural features. **(D)** The neural-behavioral similarity plot for the independent replication sample (see **Methods** for sample details) was binned by $\rho = 0.1$ & $PC3_{score} = 0.5$, using the same neural similarity index ($\rho[\Delta GBC^{obs}, \beta_{PC3}GBC^{obs}]$ within selected 39 parcels) and projected *PC3* symptom scores. Although the grid is far more sparse (total N=69 for this sample), a similar pattern between PC3 symptom score and neural similarity index emerges.

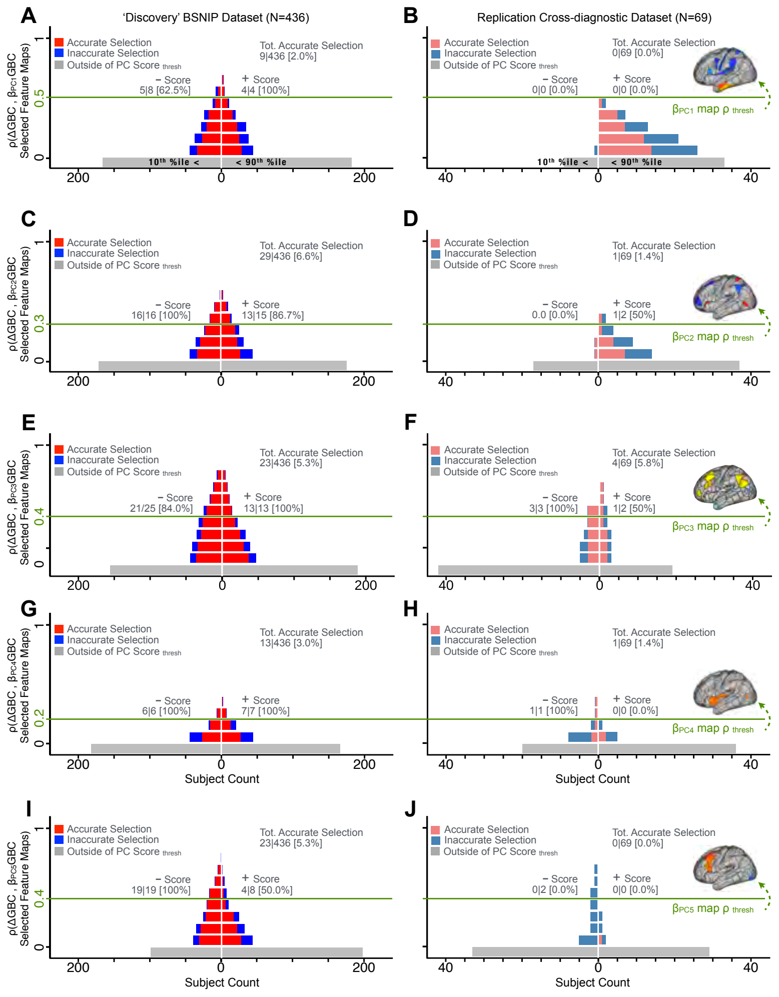


**Appendix 1 Figure 29. Evaluating patient-specific similarity to derived symptom-neural targets via data-reduced symptom scores for all *PCs*.** As noted in the main text for *PC3*, here we plot the *PC* symptom score (X-axis) versus the neural similarity prediction index (NSPI, **Appendix 1 - Fig.** 27) for all 436 PSD patients. The NSPI is defined as the Spearman's correlation between $\Delta GBC^{obs}$ and the $\beta_{PC3} GBC^{obs}$ map of the maximally-predictable "selected" $P$ parcels (Y-axis for each plot). Alternative metrics were evaluated in **Appendix 1 - Fig.** 28. **(A)** A parameter sweep was performed for 0<|rho|<1 in bins of 0.1, for all subjects with *PC1* score<-3.97 ("negative behavioral threshold" set at the 10th percentile ($P_{10\text{-Disc}}$) of the distribution and for all patients with *PC1* score>+3.60 ("positive behavioral threshold" set at the 90th percentile ($P_{90\text{-Disc}}$) of the distribution). The number of patients accurately selected within each bin is shown in red; the number of inaccurately selected patients is shown in blue. Note that this selection process excludes a number of patients (shown in grey) who do not meet the PC threshold cutoff. Note that as the neural $\rho$ threshold increases the proportion of accurate to inaccurate selections increases and the number of inaccurately selected subjects decreases. For illustrative purposes, a neural threshold of $\rho$=0.5 is applied at the point where no inaccurate selections occur for the positive behavioral threshold. **(B)** The neural and symptom score thresholds defined in discovery patients were applied on an independent external "replication" dataset (N=69, see **Methods**). For the symptom score ($P_{10\text{-Disc}}$=-3.97 and $P_{90\text{-Disc}}$=+3.60 ) and neural ($\rho$=0.5) thresholds as those defined in the discovery sample, no patients are selected for *PC1*. **(C–D)** Similar data to **A–B** for *PC2*, using PC score thresholds of $P_{10\text{-Disc}}$=-2.43 and $P_{90\text{-Disc}}$=+2.32 and a PC map threshold of $\rho$=0.3 on a subset of $P = 68$ optimized parcels. **(E–F)** Similar data to **A–B** for *PC3* using PC score thresholds of $P_{10\text{-Disc}}$=-2.41 and $P_{90\text{-Disc}}$=+2.17 and a PC map threshold of $\rho$=0.4 on a patients of $P = 39$ optimized parcels. **(G–H)** Similar data to **A–B** for *PC4*, using PC score thresholds of $P_{10\text{-Disc}}$=-1.69 and $P_{90\text{-Disc}}$=+1.72 and a PC map threshold of $\rho$=0.2 on a subset of $P = 50$ optimized parcels. **(I–J)** Similar data to **A–B** for *PC5*, using PC score thresholds of $P_{10\text{-Disc}}$=-1.89 and $P_{90\text{-Disc}}$=+1.47 and a PC map threshold of $\rho$=0.4 on a subset of $P = 31$ optimized parcels.

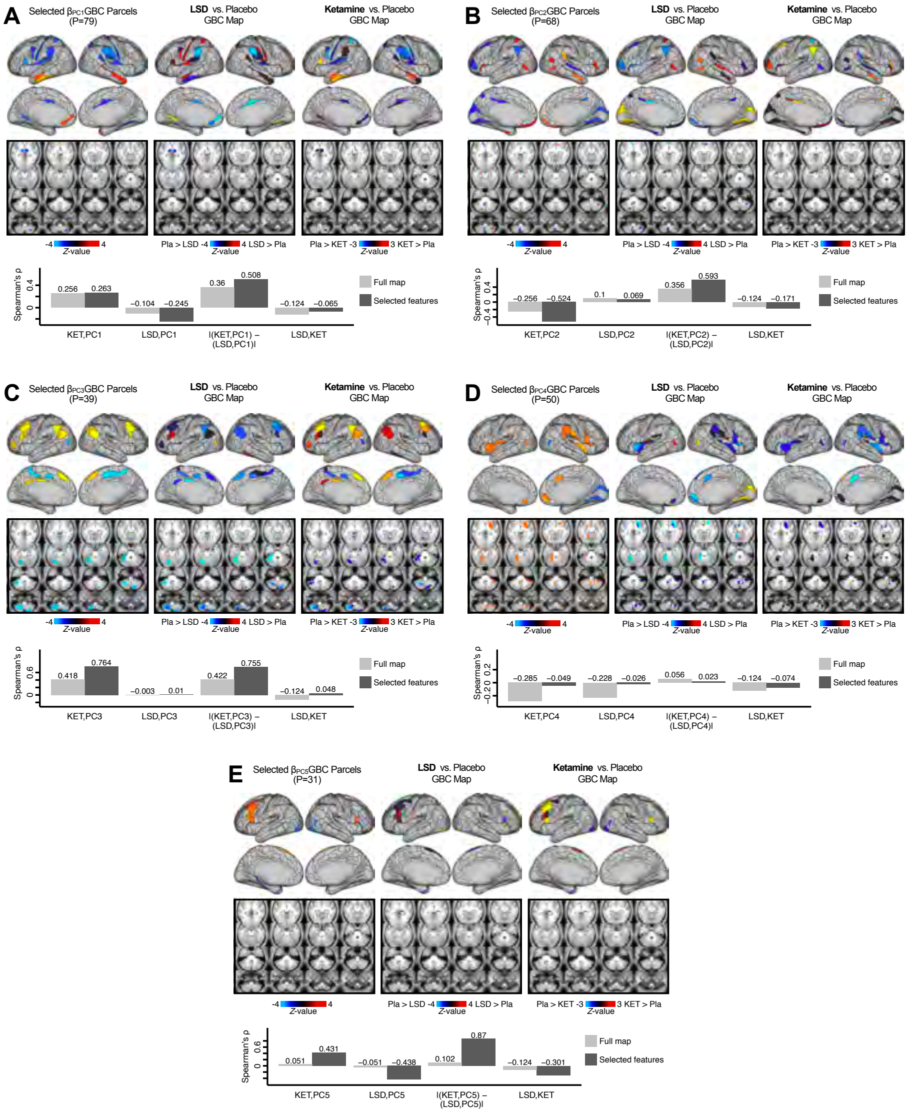


**Appendix 1 Figure 30. Examining most predictive parcels for all *PC*-driven symptom-neural effects and their relationship with pharmacological neuroimaging maps. (A)** (Left) The selected group-level $\beta_{PC1}GBC$ map showing the 79 selected parcels derived from the feature selection workflow (**Appendix 1 - Fig.** 25). (Middle) The same parcels are highlighted from the group-level $\Delta GBC$ LSD map reflecting the pharmacological effect relative to placebo. (Right) the group-level ketamine $\Delta GBC$ map reflecting the pharmacological effect relative to placebo. (Bottom bar plot) Here we show four values: i) Spearman's $\rho$ between the ketamine $\Delta GBC$ and $\beta_{PC1}GBC$ maps (KET,PC1); ii) Spearman's $\rho$ between the LSD $\Delta GBC$ and $\beta_{PC1}GBC$ maps (LSD,PC1); iii) Absolute value of the difference for the (i) and (ii) (i.e. differential similarity; |(KET,PC1) - (LSD,PC1)|); iv) Spearman's $\rho$ between the ketamine $\Delta GBC$ response and LSD ketamine $\Delta GBC$ response maps (KET,LSD). These relationships are quantified within both the selected 79 features (dark grey bars) and the full *P*=718 neural map (light grey bars). **(B–E)** These analyses are repeated and shown for *PCs 2-5*. Note that a strong differential similarity for a pair of pharmacological maps may indicate a strategy for patient treatment selection along a particular brain-behavioral axis. Specifically, in this framework patients loading on *PC3* behaviorally (or neurally) would be indicated as highly similar to neural variation induced by ketamine but not LSD. Such metrics may be clinically useful for selecting one pharmacological target over another in the process making individual patient segmentation decisions.

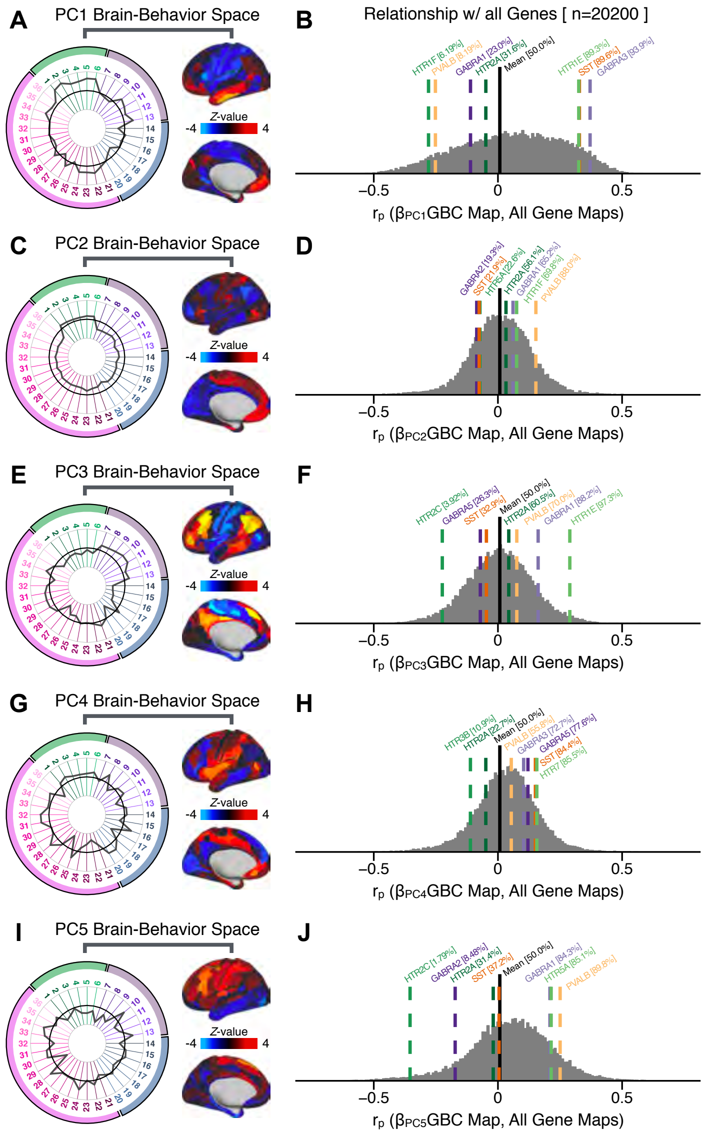


**Appendix 1 Figure 31. Quantifying the relationship between *PC*-derived symptom-neural effects and expression maps for genes implicated in psychosis spectrum disorder (PSD) neurobiology. (A)** The symptom loading profile (radarplot) and neural loading map derived in the main analyses are shown for the *PC1* BBS. **(B)** The distribution shows the Pearson's correlation coefficients between the *PC1* symptom-neural map and all 20,200 gene expression maps derived from the Allen Human Brain Atlas (AHBA, see **Methods** and for details on gene expression analyses see (***Burt et al., 2018***)). In this analysis we focused on a select number of PSD-relevant genes, as highlighted in the main text for PC3 specifically **Fig. 8**. Seven genes of interest are shown. Specifically, these include two interneuron marker genes, somatostatin (SST) and parvalbumin (PVALB); two $GABA_A$ receptor subunit genes: GABRA1 and GABRA3; and three serotonin receptor subunit genes: HTR1E, HTR1F, and HTR2A. The percentile for each highlighted gene relative to the entire distribution of 20,200 genes is reported in square brackets. Note that the gene expression maps of HTR1F and PVALB are at the negative tail of the entire distribution, i.e. anti-correlated with *PC1* symptom-neural map. Conversely, GABRA3, SST, and HTR1E are on the far positive end, reflecting a highly similar spatial pattern of the gene expression maps with the *PC1* symptom-neural map. **(C–J)** *PC* symptom-neural map profiles and distributions of gene expression map similarities are also shown for PCs 2-5. Note that some of these exemplar genes show a strong spatial similarity (or anti-similarity) with the PC-derived neural maps relative to the entire distribution of all gene-PC map pairs.
